# Mimicking opioid analgesia in cortical pain circuits

**DOI:** 10.1101/2024.04.26.591113

**Authors:** Corinna S. Oswell, Sophie A. Rogers, Justin G. James, Nora M. McCall, Alex I. Hsu, Gregory J. Salimando, Malaika Mahmood, Lisa M. Wooldridge, Meghan Wachira, Adrienne Jo, Raquel Adaia Sandoval Ortega, Jessica A. Wojick, Katherine Beattie, Sofia A. Farinas, Samar N. Chehimi, Amrith Rodrigues, Jacqueline K. Wu, Lindsay L. Ejoh, Blake A. Kimmey, Emily Lo, Ghalia Azouz, Jose J. Vasquez, Matthew R. Banghart, Kevin T. Beier, Kate Townsend Creasy, Richard C. Crist, Charu Ramakrishnan, Benjamin C. Reiner, Karl Deisseroth, Eric A. Yttri, Gregory Corder

**Author notes:** Correspondence (G.C.) & (E.A.Y.). Co-First Authors (Corinna S. Oswell, Sophie A. Rogers, Justin G. James). Co-Senior Authors (Benjamin C. Reiner, Karl Deisseroth, Eric A. Yttri, Gregory Corder).

## Abstract

The anterior cingulate cortex is a key brain region involved in the affective and motivational dimensions of pain, yet how opioid analgesics modulate this cortical circuit remains unclear. Uncovering how opioids alter nociceptive neural dynamics to produce pain relief is essential for developing safer and more targeted treatments for chronic pain. Here we show that a population of cingulate neurons encodes spontaneous pain-related behaviors and is selectively modulated by morphine. Using deep-learning behavioral analyses combined with longitudinal neural recordings in mice, we identified a persistent shift in cortical activity patterns following nerve injury that reflects the emergence of an unpleasant, affective chronic pain state. Morphine reversed these neuropathic neural dynamics and reduced affective-motivational behaviors without altering sensory detection or reflexive responses, mirroring how opioids alleviate pain unpleasantness in humans. Leveraging these findings, we built a biologically inspired gene therapy that targets opioid-sensitive neurons in the cingulate using a synthetic mu-opioid receptor promoter to drive chemogenetic inhibition. This opioid-mimetic gene therapy recapitulated the analgesic effects of morphine during chronic neuropathic pain, thereby offering a new strategy for precision pain management targeting a key nociceptive cortical opioid circuit with safe, on-demand analgesia.

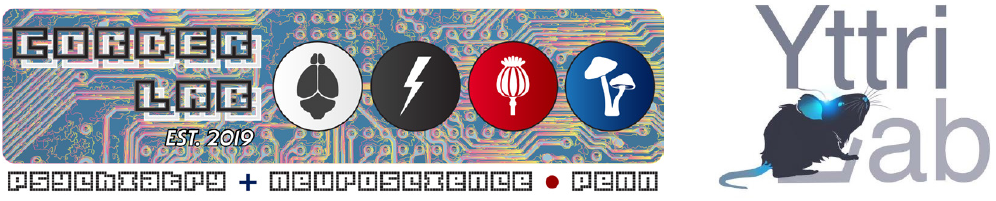

## Introduction

Pain is a complex, aversive perception, fundamental to adaptive survival ^1^. Understanding the brain neural circuits and cell-types underlying the affective and motivational features of pain experiences is crucial for advancing precision therapeutics for individuals with pathological, chronic pain conditions ^2,3^. Current analgesic drugs, such as opioids, bathe the body in chemical compounds, acting on widely expressed molecular targets to that reduce pain unpleasantness but also can promote serious and fatal side effects. By pinpointing specific brain neural circuits for pain-associated aversion, which intersect with the expression of the mu-opioid receptor (MORs) ^4^—the molecular target of morphine—a new class of effective analgesics can be developed that interfere with the unpleasantness of pain rather than pain sensation, with reduced addiction and respiratory depression side effects ^5,6^.

Coordinated neural activity in the anterior cingulate cortex (ACC) is essential for encoding the emotional and motivational dimensions of pain ^7–13^, guiding behavioral choices in real time and promoting future avoidance of harmful stimuli ^14,15^. According to Gate Control Theory, pain perception is not a passive relay of nociceptive input but is actively shaped by spinal and supraspinal circuits that integrate sensory, cognitive, and emotional information to either amplify or inhibit pain signals. Within this framework, the ACC plays a central role in evaluating nociceptive input in relation to valence, context, and internal state. This processing supports the selection of adaptive responses such as escape, protection, and recuperative behaviors ^16^ —responses that act as a form of negative feedback to reduce further injury and promote healing. By quantitatively tracking the durations, frequencies, and transitions between these spontaneous pain-related behaviors, we can infer the latent computations underlying pain affect. This approach provides a window into identifying single neurons and distributed ensembles in the ACC that represent the affective-motivational components of pain, offering candidate targets for next-generation therapeutic strategies.

Ablation of the ACC alleviates the emotional suffering of chronic pain patients, but it does not affect their ability to evaluate the intensity of their pain ^17–20^. Similarly in preclinical models, lesions ^21,22^, opioids ^23–26^, and optogenetic manipulation ^27–33^ of ACC neural circuits attenuate aspects of the affective-motivational component of the pain, as shown by a reduction in conditioned place avoidance and aversive behaviors induced by noxious stimuli. In total, these studies point to the same neural circuits in the ACC that drive adaptive behavior to acute noxious stimuli might also be maladaptive during chronic pain ^34,35^. Thus, leveraging personalized deep-brain stimulation ^36–38^ or cell-type-specific gene therapies ^39,40^ to modulate neural activity in MOR-expressing neurons ^41^ may mimic the prefrontal cortical actions of opioid analgesia without the associated risks of off-target pharmacotherapies ^1^.

To functionally link ACC neural ensembles to affective-motivational aspects of pain, we developed LUPE (Light aUtomated Pain Evaluator), a deep-learning platform that quantifies spontaneous behaviors across timescales ranging from sub-seconds to minutes. LUPE enables the identification of latent behavioral states affected by pain and analgesia, and the formation of a quantitative index for pain—the Affective-Motivational Pain Scale (AMPS). When paired with single-cell calcium imaging using head-mounted miniscopes, LUPE-AMPS facilitates the investigation of individual ACC neurons encoding pain behaviors, and states, and responding responses to analgesic interventions.

This integrated system provides the first longitudinal tracking of cortical pain neural dynamics across the onset, maintenance, and treatment phases of chronic pain.

Last, we introduce a biologically inspired, circuit-based gene therapy that mimics the cortical effects of morphine. Using a synthetic mu-opioid receptor promoter, we selectively express inhibitory chemogenetic receptors in ACC neurons responsive to opioids. This enables precise modulation of endogenous nociceptive processes—achieving targeted analgesia without altering nociceptive sensitivity or reinforcing behavior in healthy subjects. By acting directly and specifically on the circuits we identified to mediate the affective unpleasantness of pain, this approach offers a path toward non-addictive, cell-type-specific pain relief.

## Results

### Mapping a nociceptive hotspot in µ-opioidergic ACC neurons

To identify pain-responsive neurons in the ACC, we used the TRAP2 activity-dependent transgenic mouse for nociceptive neural tagging (*pain*TRAP) ^1,42^. Repeated noxious 55°C stimuli to the hindpaw activated *Fos*-driven Cre recombination, permanently labeling nociceptive neurons with tdTomato throughout the nervous system (**Fig. S1a–b**). Labeled *pain*TRAP+ neurons were distributed across the dorsal (Cg1) and ventral (Cg2) ACC but concentrated within a ∼700-μm segment of the ACC (+1.69 to +0.97 mm AP; **Fig. S1b-d**), anterior to known mid-cingulate circuits for reflexive pain responses ^43,44^. Two weeks later, mice were re-exposed to the same noxious stimulus, and we immunostained for FOS protein (*pain*FOS) to assess reactivation of the *pain*TRAP+ neural population. Compared to no-stimulus controls (**Fig. S1e-g**), *pain*TRAP mice showed a 2.3-fold increase in double-labeled painTRAP+/painFOS+ neurons (**Fig. S1c,f**), indicating stable recruitment of a pain-reactive ACC neural ensemble.

Given the dense expression of mu-opioid receptors (MORs) in the ACC ^45–48^ and their known role in analgesia ^25,49^, we next asked whether pain-responsive neurons in this region express the MOR-encoding gene *Oprm1*. Using fluorescent in situ hybridization and HALO deep-learning histology quantification, we measured co-expression of *Oprm1* and noxious stimulus-induced *Fos* mRNA (*painFos*) in male and female mice with either uninjured or neuropathic pain conditions (**Fig. S2**). On average, ∼30% of *painFos*+ neurons expressed *Oprm1* in uninjured mice, increasing to ∼50% during chronic pain (**Supp. Fig. 2c–d**), indicating enhanced recruitment of MOR-expressing ACC neurons in persistent pain states.

To further refine the identity and distribution of these opioidergic nociceptive neurons, we mapped *Oprm1* expression across ACC layers. Approximately 71% of *Oprm1*+ cells in Cg1 co-expressed the excitatory pyramidal neuron marker *Slc17a7* (VGLUT1; 9,488 of 13,047 cells) and were evenly distributed across cortical layers (**Fig. S2b-d**). However, both the proportion of *Oprm1*+ cells and mRNA transcript density per cell increased down the layers in a graded manner, similar to quantifications in MOR-mCherry mice ^50^, but in contrast, we did not detect sex differences. Consistent with *pain*TRAP counts (**Fig. S1**), *painFos* expression was highest in Cg1 Layer 2/3, where ∼25% of *Oprm1*+ cells also expressed *painFos*, suggesting that specific subsets of excitatory MOR-expressing neurons in superficial *vs*. deep layers may be preferentially engaged by noxious events.

### Molecular profiling of opioid and nociceptive ACC cell-types over the development of chronic pain

To identify ACC cell types involved in acute *vs*. chronic neuropathic pain processing and opioid receptor-mediated analgesia, we performed single-nucleus RNA sequencing (snRNA-seq) from neurons within the nociceptive hotspot defined by *pain*TRAP (**Fig. S3**). We collected bilateral ACC punches from mice at 3 days, 3 weeks, and 3 months after bilateral Spared Nerve Injury (SNI ^51^), along with uninjured controls (n = 4 mice per injury condition/timepoint), for subsequent Differential Expressed Gene (DEG) analysis over the development of chronic pain ^52^. To enrich for allodynia-relevant activity, we applied a light-touch (0.16 g) mechanical stimulus to the hypersensitive, injured paws 5 minutes before tissue collection to induce immediate early gene (IEG) mRNA expression (**Fig. S3a**). We analyzed 104,302 single nuclei from the n = 16 mice and identified 23 distinct cell-type clusters (**Fig. S3b**), including 14 glutamatergic, 3 GABAergic, and 6 non-neuronal populations (**Fig. S3c,d**). Cluster distributions were consistent across all injury conditions (**Fig. S3c,i**).

We next assessed the expression of opioid receptors (*Oprm1, Oprd1, Oprk1, Oprl1*) and their peptide ligands (*Penk, Pdyn, Pomc, Pnoc*; **Fig. S3e**) ^53^. In alignment with others ^50^ and our fluorescence *in situ* mapping (**Fig. S2**), *Oprm1* showed a graded increase across glutamatergic neurons, from Layer 2/3 intratelencephalic (IT) types to Layer 6 corticothalamic (CT) types (**Fig. S3e, h**). In contrast with some reports ^50,54,55^, we observed limited expression in *Sst* or *Vip* GABAergic interneurons. To identify nociceptive cell types, we used two orthogonal IEG-based activity measures: a weighted activity score and Nebulosa density mapping across 25 IEGs (*pain*IEGs; **Fig. S3f–g**). Only three cell types—L2/3 IT-3 (Cluster 3), L5 IT-1 (Cluster 6), and L6 CT-2 (Cluster 13)—showed consistent *pain*IEG activation across all three SNI timepoints – all of which expressed Oprm1 (**Fig S3h**).

To determine whether chronic pain alters ACC gene expression over time, we performed differential expression analysis across all 23 clusters using a pseudo-bulk edgeR approach (FDR < 0.05, log_2_FC > |0.25|). At 3 days post-SNI, we identified 3,583 DEGs, with 43, 208, and 184 DEGs for the *pain*IEG+ clusters, L2/3 IT-3, L5 IT-1 and LC CT-2, respectively (**Fig. S3j**). Synaptic gene ontology (SynGO) analysis of these DEGs revealed changes in synaptic translation, vesicle cycling, and synapse organization (**Fig. S3k**) ^56^. Contrary to bulk-sequencing reports without cell-type resolution ^57–59^, few DEGs remained at 3 weeks (n = 635 total) and 3 months (n = 180 total) across all clusters (**Fig. S3j**), making cluster-specific pathway analysis statistically infeasible. These findings suggest that despite persistent pain behaviors, ACC gene expression largely returns to a baseline state over time. This mirrors findings in peripheral dorsal root ganglia neurons, where transcriptional changes resolve by 2–8 weeks post-injury ^60^. While some reports show decreased MOR expression in the mid-cingulate cortex with chronic pain ^61,62^, *Oprm1* expression in the anterior ACC remained stable across all conditions and cell types, with no change detected at any timepoint post-SNI—confirmed by both snRNAseq and FISH in male and female mice (**Fig. S2**) ^63^. This stability supports the feasibility of targeting ACC MOR+ neurons for MOR promoter-based cell-type-specific therapies, independent of chronic pain-induced transcriptomic changes.

### ACC MORs are necessary and sufficient for morphine analgesia

Given prior evidence for the role of ACC MORs in mediating opioid pain relief ^25^, we next sought to test the necessity of ACC MORs expressed by ACC neurons to facilitate morphine analgesia on both the sensory-reflexive and affective-motivational components of pain behaviors ^1^ (**Fig. S4a,b**). First, we knocked out *Oprm1* in ACC neurons by injecting AAV9-*hSyn*-eGFP-Cre into *Oprm1*^Flox/Flox^ mice^64^ (**Fig. S4c,e**), leaving MORs intact elsewhere in the nervous system. qPCR confirmed >50% reduction in *Oprm1* (**Fig. S4d**). Following a systemic dose of morphine (0.5 mg/kg, i.p. ^25,26,65^) shown to selectively mediating affective analgesic responses without altering sensory thresholds, mice were tested on a battery of evoked-sensory stimuli tests to calculate changes in analgesia. Deletion of ACC MORs did not affect mechanical withdrawal thresholds, but significantly impaired morphine’s effect to reduce affective-motivational behaviors (*e*.*g*., paw licking, biting, guarding, and escape ^42,64,66^; **Fig. S4a**) in response to noxious 55°C water and 6°C acetone stimuli, relative to controls with intact MOR expression (**Fig. S4f–g**). One week later, the same mice were tested on a 60-second exposure to an inescapable 50°C hotplate. ACC MOR deletion blunted morphine-induced analgesia, as reflected in increased duration of licking and guarding behaviors (**Fig. S4h**). In controls, morphine increased the latency between the initial reflexive paw withdrawal and the onset of paw licking behavior—from ∼1 second to ∼7.2 seconds. This dissociation between reflexive and affective responses was abolished in mice lacking ACC MORs (**Fig. S4i**). Moreover, once bouts of affective-motivational behaviors began, ACC MOR-deleted mice spent more time engaged in licking (**Fig. S4j**), indicating a loss of morphine suppression of aversive motivational drive.

To test sufficiency, we re-expressed human MOR (hMOR) in the ACC of global MOR knockout mice (MOR KO; *Oprm1*^Cre/Cre^) ^67^ using a Cre-dependent AAV.DJ-*hSyn*-FLEx-mCherry-2A-hMOR ^68^ (**Fig. S4k**). qPCR and mCherry fluorescence confirmed successful hMOR expression (**Fig. S4l,m**). In MOR KO mice, morphine (0.5 mg/kg) had no effect on affective-motivational pain behaviors, but hMOR re-expression restored morphine analgesia in response to noxious 55°C water and 6°C acetone stimuli (**Fig. S4n–o**). On the inescapable hotplate assay, morphine in hMOR-re-expressing mice increased the latency to initiate licking and reduced total engagement in affective-motivational behaviors (**Fig. S4p**). This also restored the temporal dissociation between reflexive and affective responses (**Fig. S4q–r**), mirroring the wild-type phenotype.

These rapid shifts in affective-motivational and pain-recuperative behaviors suggest that morphine modulates ACC MOR-expressing neurons to suppress affective nociception and thereby deprioritize pain-related action selection, underscoring the need for more precise, unbiased tools to quantify subsecond behavioral dynamics beyond the limits of manual scoring.

### A standardized machine-vison and deep-learning system for complex pain behaviors

The complexity of pain cannot be fully captured by reflexive withdrawal responses—the current standard for preclinical analgesic evaluation—which assess only evoked responses and fail to reflect the ongoing, affective experience most relevant to patients ^69–71^. While assays such as conditioned place preference or aversion measure memory-based responses to prior pain or relief ^21,72,73^, they do not capture the dynamic, moment-to-moment motivational behaviors driven by spontaneous or ongoing pain ^1,74^.

To address these limitations, we developed LUPE (Light aUtomated Pain Evaluator; **Fig. 1**), a behavioral analysis platform designed to resolve fine-scale, naturalistic pain-related behaviors across multiple timescales. LUPE enables a more nuanced and translationally relevant assessment of affective-motivational pain states in freely moving mice. Named after the Greek daemon of pain and suffering (Lýpē, λŪπ́η), LUPE provides a standardized, dark, and safe environment optimized for the behavior of nocturnal prey animals. Critically, it eliminates the presence of the human experimenter—both as a looming threat and as a subjective observer—allowing for objective quantification of spontaneous and nocifensive behaviors. This setup prioritizes the detection of behavioral transitions as a primary metric for inferring pain or analgesia, capturing the dynamic and ethologically grounded nature of pain expression.

**Fig. 1.**
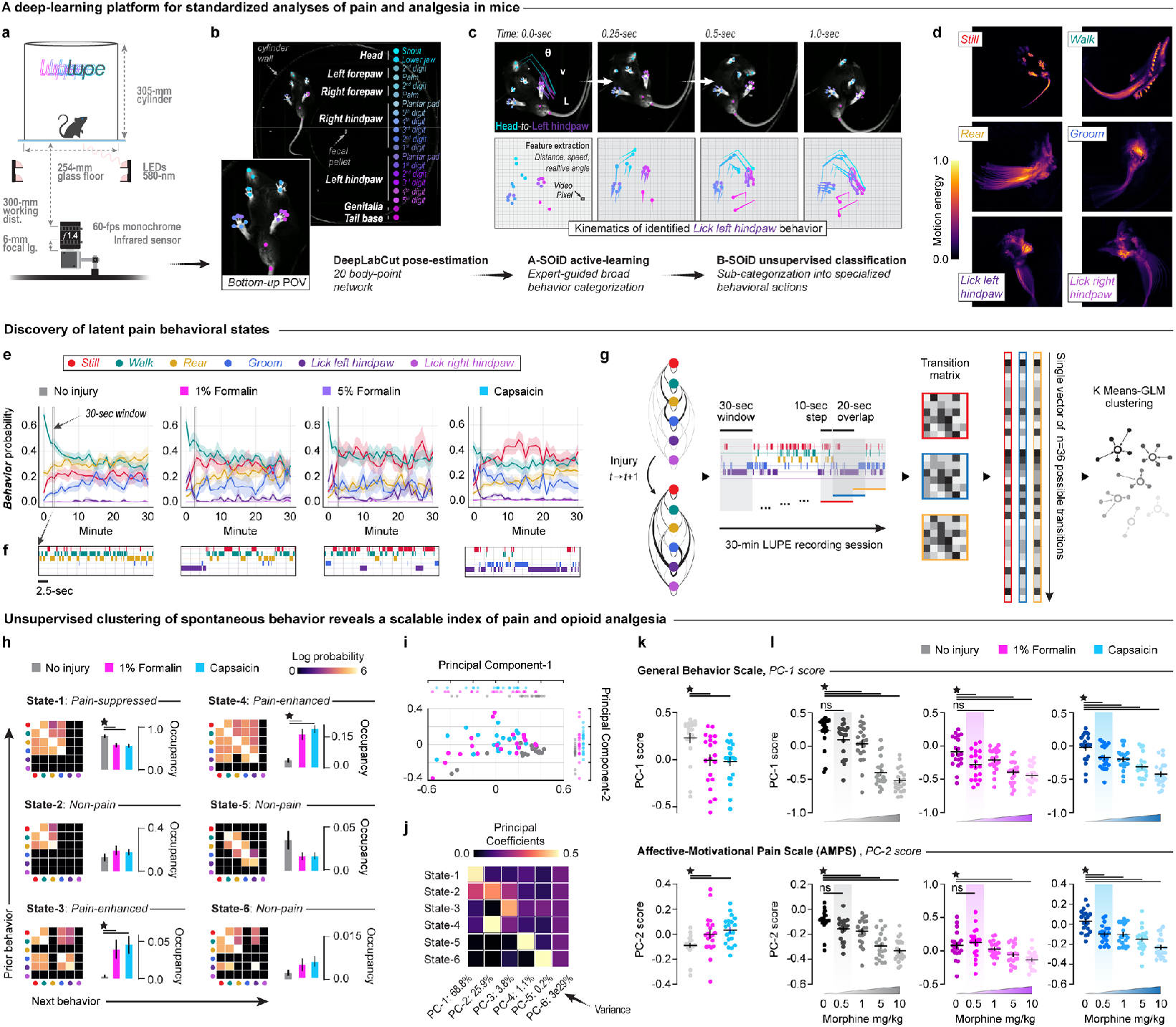
Deep learning analysis of natural behavior reveals how pain and opioids shape internal affective-motivational states. **(a)** Schematic of the standardized LUPE platform and chamber for high-speed infrared videography. **(b)** A 20-body point DeepLabCut pose-tracking model was built from a bottom-up camera for male and female mouse pain behavior. **(c)** Behavior segmentation models trained iteratively on supervised annotations of holistic behaviors (*e*.*g*., licking *vs*. grooming *vs*. walking *vs*. rearing etc.) with A-SOiD, followed by further unsupervised sub-clustering with B-SOiD. **(d)** Motion energy heat maps illustrating spatial trajectories and intensity distributions for each of the six primary behavioral repertoires. **(e)** Temporal probability plots for each of the six primary behavior repertoires in 1-min bins, comparing mice in uninjured conditions to those injured by left hindpaw injections of 1% formalin, 5% formalin, or capsaicin. **(f)** Raster plots of rapid behavioral transitions within a 30-sec window from uninjured, formalin, and capsaicin injuries. **(g)** Procedure for behavioral state inference from statistical structure of spontaneous behavior. **(h)** *Left panels*: Model centroid transition matrices characterizing each of six inferred states. *Right panels*: Comparing fraction occupancy of mice in each state between uninjured (gray), formalin (magenta), and capsaicin (Cyan) pain models (n=20/group; 1-way ANOVA + Tukey: *p*State 1 = 0.0007, *p*State 3 = 0.0078, *p*State 4 < 0.0001). **(i)** Two-dimensional visualization of PCA of state occupancies across pain models. Dots are individual animals. **(j)** Magnitude of coefficients of each state in each PCA. **(k)** Scores of each animal along PC1 (top) and PC2 (bottom) across pain models (n=20/group; One-way ANOVA, Tukey correction: pPC1 = 0.0027, pPC2 = 0.0082). **(l)** Dose-response of morphine on PC1 (top) and PC2 (bottom) scores in uninjured, formalin, and capsaicin administered mice (n=20/group and dose; 1-way ANOVA + Tukey: *p*PC1, uninjured < 0.0001, *p*PC1 formalin < 0.0001, *p*PC1 capsaicin < 0.0001, *p*PC2, uninjured < 0.0001, *p*PC2, formalin < 0.0001, *p*PC2, capsaicin < 0.0001). ⋆= P < 0.05. Bars, lines, or dots are mean; error bars and shaded areas are SEM. See ***Supplementary Table 1*** for statistics.

In addition to the standardized LUPE chamber and high-speed infrared videography recorded from below through a glass floor (**Fig. 1a, S5**), behavioral classification is driven by a multi-layered analysis pipeline. Using DeepLabCut ^75^ to track 20 body key points, LUPE extracts detailed posture dynamics that are processed through both semi-supervised (A-SOiD ^76^) and unsupervised (BSOiD ^77^) algorithms to identify six holistic behavioral repertoires: *Still, Walk, Rear, Groom, Lick Left Hindpaw*, and *Lick Right Hindpaw* (**Fig. 1b,c, S6**). These repertoires are assembled from sub-second behavioral syllables and allow quantitative analysis of transitions across time. Motion energy plots, which quantify the displacement of tracked body points, visualize the movement patterns that define each behavior and distinguish similar actions such as grooming and paw-directed licking (**Fig. 1d**).

To evaluate LUPE’s sensitivity to dynamic changes in pain-related behavior, we applied the formalin and capsaicin models of acute pain ^78^. Male and female C57Bl/6J mice were habituated to the LUPE chamber for two consecutive days, then injected into the left hindpaw with 1% or 5% formalin, 2% capsaicin, or left uninjured as controls (**Fig. 1e, S5c**). LUPE computed behavioral probabilities for all six repertoires in 1-minute bins over 30-minute sessions for all 60 mice in under two hours—compared to 50–150 minutes for manual BORIS ^79^ scoring of a single behavior in a single mouse (estimated upper bound = 54,000 minutes for the full dataset; **Fig. S5k**). By automating behavior classification with high temporal resolution, LUPE increases the speed, rigor, and reproducibility of preclinical pain behavior analysis. It also generates archival-quality datasets that include video logs and computer-scored results, facilitating transparent cross-laboratory comparison, long-term record-keeping, and future reanalysis.

### Discovery of latent affective-motivational pain states sensitive to morphine

A central challenge in interpreting spontaneous behavior is that an animal’s internal state—such as pain—is not directly observable. For example, an injury to the left hindpaw may or may not elicit licking at a given moment, yet the animal may still be experiencing pain. Thus, we hypothesized that latent cognitive-affective states could be inferred from patterns in sparse, spontaneous, and multivalent behavior. To identify latent pain states in acute and chronic pain models (**Fig. 1g, S7**), we analyzed LUPE-scored behaviors from 58 male and female mice subjected to formalin (n=19), capsaicin (n=20), or spared nerve injury (SNI; n=19). Behavioral transitions were modeled as Markov processes using thirty-second sliding windows, from which we generated transition matrices for each animal (**Fig. 1g, S7a**). Next, transition matrices were clustered using k-means (k=6; 100-fold cross-validation) based on silhouette and elbow method optimization (**Fig. 1g, S7b**). Classification of individual animals’ behavior to these clusters significantly exceeded chance across all conditions, as measured by Euclidean distance between cluster centroids from real versus shuffled data (**Fig. S7c–h**). Importantly, clustering did not rely disproportionately on any single behavior, as systematic removal of individual behaviors disrupted classification less than expected by chance (**Fig. S7c**).

The centroids of each cluster are represented by mean transition matrices that define distinct behavioral states (**Fig. 1h**). *State 1* is characterized by transitions among stillness, walking, rearing, and grooming, while *State 2* includes only stillness, walking, and rearing. *State 3* adds licking the injured paw to the repertoire of *State 1*, and *State 4* includes all behaviors except licking the uninjured paw. *State 5* involves transitions between all behaviors except stillness, whereas *State 6* includes stillness, walking, rearing, and licking the injured paw, but not grooming. States evolved slowly, on the order of seconds to minutes, and exhibited unique dynamics that were conserved across pain models (**Fig. S7i-k**).

Consistent with our hypothesis, behavioral states distinguished injured from uninjured conditions, but did not differentiate between injury types (**Fig. 1h, Supp. Table 1, Rows 1–6**). Uninjured animals most frequently occupied *States 1* and *2*. In contrast, both capsaicin and formalin increased occupancy of *States 3* and *4*, while reducing time spent in *State 1*. Among these, *State 4* was uniquely and dose-dependently suppressed by morphine, suggesting it may represent a selectively opioid-sensitive dimension of spontaneous pain **Fig. S8g, Supp. Table 1, Rows 46-48**). However, all states were dose-dependently modulated by morphine in various directions and extents, across injury conditions (**Fig. S8g, Supp. Table 1, Rows37-54**). Thus, latent states spanning affective-motivational relevant timescales inferred from sparse, spontaneous behavior track pain and analgesia.

### A quantitative index for pain captures the bidirectional effects of injury and analgesia

To compress the robust yet diverse effects of pain and morphine across all six states, we applied principal component analysis (PCA) to the fraction of time each mouse spent in each state across experiments (**Fig. 1i**). This revealed two principal axes of variation in behavior across pain conditions. Both capsaicin and formalin shifted scores along these axes—reducing the first component and increasing the second—regardless of injury model (**Fig. 1k, Supp. Table 1 Rows 7-8**).

The first principal component (PC1), driven primarily by States 1 and 2, reflects a baseline behavioral structure disrupted by both injury and high-dose morphine, and is termed the General Behavior Scale (**Fig. 1j, 1l top, Supp. Table 1 Rows 9-11**). The second component (PC2), weighted by States 2 and 4, was selectively increased by injury and dose-dependently suppressed by morphine (**Fig. 1l, bottom, Supp Table 1 Rows 12-14**), capturing the presence and relief of affective pain. Because PC2 responds bidirectionally to both injury and analgesia, we define it as the *Affective-Motivational Pain Scale* (AMPS)—a data-driven, continuous index of pain-related behavioral states.

### Licking as a structured motivational response to affective acute and chronic pain

Gate-control theory proposes that volitional, recuperative behaviors such as rubbing or licking injured tissue serve as antinociceptive responses, where activation of touch-sensitive afferents inhibits nociceptive signaling at the spinal level (**Fig. S9a**). In this framework, the aversive quality of pain—its affective unpleasantness—drives the motivation to engage in innate behaviors like licking the injured site. This behavior, in turn, reduces pain, forming a negative feedback loop. Thus, motivated licking is expected to increase with affective pain and decline as analgesia—or recuperation—is achieved.

Our modeling approach takes a neutral stance on the functional significance of licking, treating latent behavioral states as Markovian processes in which the probability of each behavior remains stable over time within a given state, independent of behavioral history (**Fig. S9b, top**). To assess whether licking dynamics aligned with theoretical predictions, we examined the probability of each behavior as a function of time elapsed within *Pain State-4*—a latent state consistently enhanced by injury and dose-dependently suppressed by morphine (**Fig. 1h**). Across all injury models (spared nerve injury, formalin, and capsaicin), licking of the injured paw followed a consistent temporal pattern: it began near zero at the onset of *Pain State-4*, accumulated during the latter half of the state, and declined just before the transition out of *Pain State-4* (**Fig. S9b,c, Supp Table 1 Rows 55-58**). While licking occurred in other states, this temporal structure was unique to *Pain State-4* and not observed in other behaviors or in licking across all states (**Fig. S9d,e**). These findings suggest that paw licking is not merely a reflexive pain behavior, but an innate affective-motivational response engaged to negatively modulate pain—consistent with gate-control theory and its proposed role as a functional and motivated antinociceptive behavior.

### Neural dynamics in ACC track nociceptive stimuli and behaviors

To identify the neural correlates of morphine-suppressed pain behavior in the ACC, we performed single-cell calcium imaging in freely behaving mice inside the LUPE chamber. We expressed AAV9-hSyn-jGCaMP8m in ACC neurons and implanted 1.0-mm GRIN lenses at nociceptive hotspot coordinates (n = 5 male mice; **Fig. 2a,b, S10a,b**). Mice were connected to a head-mounted Inscopix nVista 3.0 one-photon microscope, enabling simultaneous recording of neural activity and behavior as they experienced acute inflammatory pain (left hindpaw intraplantar injection of 2% capsaicin, 10 μL) and subsequent morphine-induced analgesia (0.5 mg/kg; **Fig. 2c,d**). Using a Fisher linear decoder with 100-fold cross-validation, we found that spontaneous behaviors could be reliably decoded from ACC population activity across mice and imaging sessions, regardless of injury or opioid treatment (**Fig. 2e, S10g,h**).

**Fig. 2:**
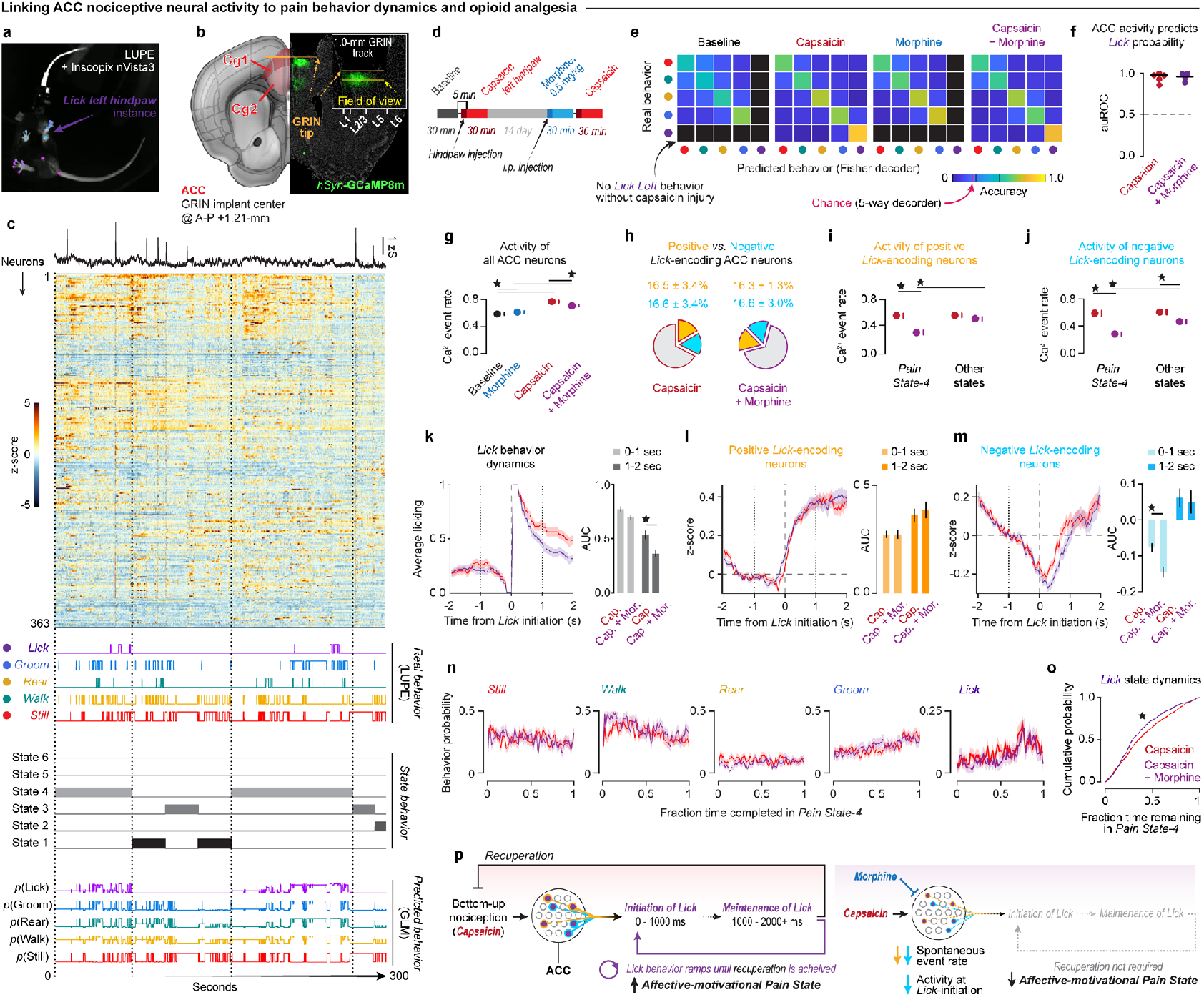
Neural dynamics in ACC track acute pain and analgesia. **(a)** Microendoscope calcium imaging synced with LUPE behavior tracking. **(b)** GRIN lens implant and *hSyn*-GCaMP8m expression in ACC Cg1. Yellow FOV bar = 1.0-mm. **(c)** *From top to bottom panels*: Average and single cell neural activity (z-score) from a representative mouse, aligned with behaviors, states inferred by our behavioral state model, and probability of behaviors given states and behavior history (binomial GLM). **(d)** Imaging protocol during capsaicin injury (i.pl, 2%, left hindpaw) and morphine (i.p., 0.5 mg/kg; n=5 mice). **(e)** Fisher decoder accuracies predicting behaviors from neural activity, averaged over mice. (Permutation test, **Fig. S10g**). **(f)** auROC of GLMs predicting *p*Lick from (e) in each animal (n=5). **(g)** Calcium events per second of all ACC neurons in all sessions. **(h)** Mean ± SEM fraction of positive and negative *p*Lick neurons during capsaicin (red outline) and capsaicin+morphine (purple outline) sessions. **(i,j)** Calcium events per second of positive (i) and negative (j) *p*Lick neurons in capsaicin and capsaicin+morphine sessions. **(l,m)** *Left*: average activity in positive (l) and negative (m) *p*Lick neurons at lick bout onset, pooled across animals. *Right*: AUC of Lick probability from 0-1 seconds post-initiation and 1-2 seconds post-initiation (Unpaired t-test: *p*Negative, 0-1s = 0.0003). **(n)** Behavior probability as a function of fraction time in *Pain State-4* in all capsaicin injured mice administered 0.0 and 0.5 mg/kg morphine (n=19-20/group). **(o)** Cumulative lick probability over fraction of time remaining in *Pain State-4* between groups (Kologomorov-Smirnov test, p = 1.1e-23). **(p)** Summary of results. Morphine inhibits ACC neurons encoding affective-motivational pain behaviors, disrupting the pain-recuperation loop and reducing pain state expression.⋆ = P < 0.05. Bars, lines, or dots are mean; error bars and shaded areas are SEM. See ***Supplementary Table 2*** for statistics.

To further characterize the functional organization of ACC ensembles, we tested whether distinct neuronal populations encode nociceptive signals based on sensory modality or valence. In uninjured animals, we delivered mechanical and thermal stimuli to the left hindpaw (0.16-g filament, pin prick, 30°C water drop, 55°C hot water drop, or 6°C acetone drop) and compared responses to orally consumed stimuli of opposing valence—appetitive 10% sucrose solution and aversive bitter quinine – and a 55°C hot water drop. ACC neurons showed greater overlap in responses to noxious stimuli across modalities (∼11–20% overlap among excited cells, ∼25% among inhibited cells) than to stimuli of opposite valence. For example, overlap among excited cells responding to 55°C heat and sucrose was only 6%, and 13% for heat and quinine; overlap among inhibited cells was ∼11% for both pairings (**Fig. S11a,c, S12e**). Furthermore, neurons activated by 55°C water versus sucrose exhibited opposing activity patterns and performed poorly when used to decode responses to the other stimulus (**Fig. S12a–e**), consistent with valence-specific encoding in ACC neural populations.

### Morphine inhibits pain-tracking neurons to induce analgesia

Having established that licking behavior scales with pain, we hypothesized that neurons encoding lick probability would be targets of morphine-induced analgesia. To estimate lick probability over time within individual animals, we aligned neural activity to behavior and latent states and trained a binomial generalized linear model (GLM) to predict licking based on current state and recent behavioral history (two prior time steps; **Fig. 2c**). These state-based GLMs performed significantly better than chance and produced a pseudo-continuous vector of lick probability over time (**Fig. 2c**). To identify neurons encoding the probabilities of specific behaviors, we trained GLMs to predict behavior probability from the principal components accounting for 80% of neural variance in each animal and session (**Fig. S10c**). These models consistently outperformed GLMs trained on shuffled data, confirming reliable mapping between population activity and behavior probability (**Fig. 2f, S10d, Supp. Table 2 Rows 11-15**). Neurons with the highest absolute weights along the three most significant principal components (p < 0.001, |z-score of neuron coefficient| > 1.5) were classified as behavior probability–encoding neurons (**Fig. S10e,f**).

As expected, ACC neurons exhibited higher spontaneous activity during capsaicin sessions compared to non-capsaicin sessions, and this activity was selectively suppressed by 0.5 mg/kg morphine only in the presence of injury (**Fig. 2g, Supp. Table 2 Row 1**). Neurons encoding lick probability—hereafter referred to as *p*Lick neurons—were further classified as positive or negative depending on whether their activity increased (16.5 ± 3.4% of cells) or decreased (16.6 ± 3.4% of cells) at lick bout onset (**Fig. 2h**). Morphine inhibited both subpopulations, but in distinct ways: activity in positive *p*Lick neurons was suppressed selectively during *Pain State-4*, whereas negative *p*Lick neurons were inhibited more broadly across states (**Fig. 2i,j, Supp. Table 2 Rows 2-3**). These results suggest that morphine targets *p*Lick neurons in a population- and state-dependent manner.

We next asked whether morphine modulates the dynamics of *p*Lick neuron activity around individual lick bouts. Behaviorally, morphine reduced the probability of licking 1–2 seconds after lick onset, suggesting decreased lick maintenance (**Fig. 2k, Supp. Table 2 Rows 4-5**). At the neural level, morphine did not alter the activation of positive *p*Lick neurons at lick onset but enhanced the inhibition of negative *p*Lick neurons 0–1 second after lick offset— preceding the behavioral change (**Fig. 2l,m, Supp. Table 2 Rows 6-9**). Together, these state- and behavior-specific effects increased the selectivity of *p*Lick neurons for licking relative to other behaviors (**Fig. S10l, Supp. Table 2 Rows 16-17**).

Last, to understand how morphine alters lick dynamics over longer timescales, we compared lick dynamics in capsaicin mice over *Pain State-4* bouts between mice that received 0.0 (saline) or 0.5 mg/kg of morphine (**Fig. 2n**). Interestingly, morphine seemed to sharpen the lick probability curve over *Pain State-4*, such that there was a reduced probability of licking earlier in the state and an even greater accumulation of licks at the end (**Fig. 2o, Supp. Table 2 Row 10**).

In summary, these data suggest that morphine alleviates the negative affective component of pain, reducing the motivational drive to engage in recuperative behaviors. This is reflected in the altered licking dynamics during *Pain State-4*—characterized by delayed initiation and reduced maintenance of licking—potentially mediated by suppression of spontaneous activity in positive *p*Lick neurons and enhanced lick behavior-locked inhibition of negative *p*Lick neurons (**Fig. 2p**).

### Morphine restores ACC dynamics related to affective motivational behaviors disrupted by chronic pain

To investigate how behavior and ACC neural dynamics adapt following the onset of chronic pain, we performed longitudinal miniscope calcium imaging in n=18 mice expressing *hSyn*-GCaMP8m + GRIN lens in Cg1 ACC. Mice were recorded in LUPE for 30 minute session at one day before, and at 1, 7, 14, and 21 days after SNI (n = 9) or remained uninjured (n = 9; **Fig. 3a**). SNI produced an immediate and sustained increase in spontaneous lick rate at the injured left hindpaw, pain state occupancy, and AMPS scores relative to uninjured controls (**Fig. 3b–e, Supp. Table 3 Rows 1-3**). As with an acute capsaicin injury (**Fig. 1**), a single 0.5 mg/kg dose of morphine effectively reduced AMPS scores in SNI mice at three-weeks post-injury (**Fig. 3f, Supp. Table 3 Row 4**), indicating that opioids remain effective for treating affective-motivational features of chronic neuropathic pain.

**Fig. 3:**
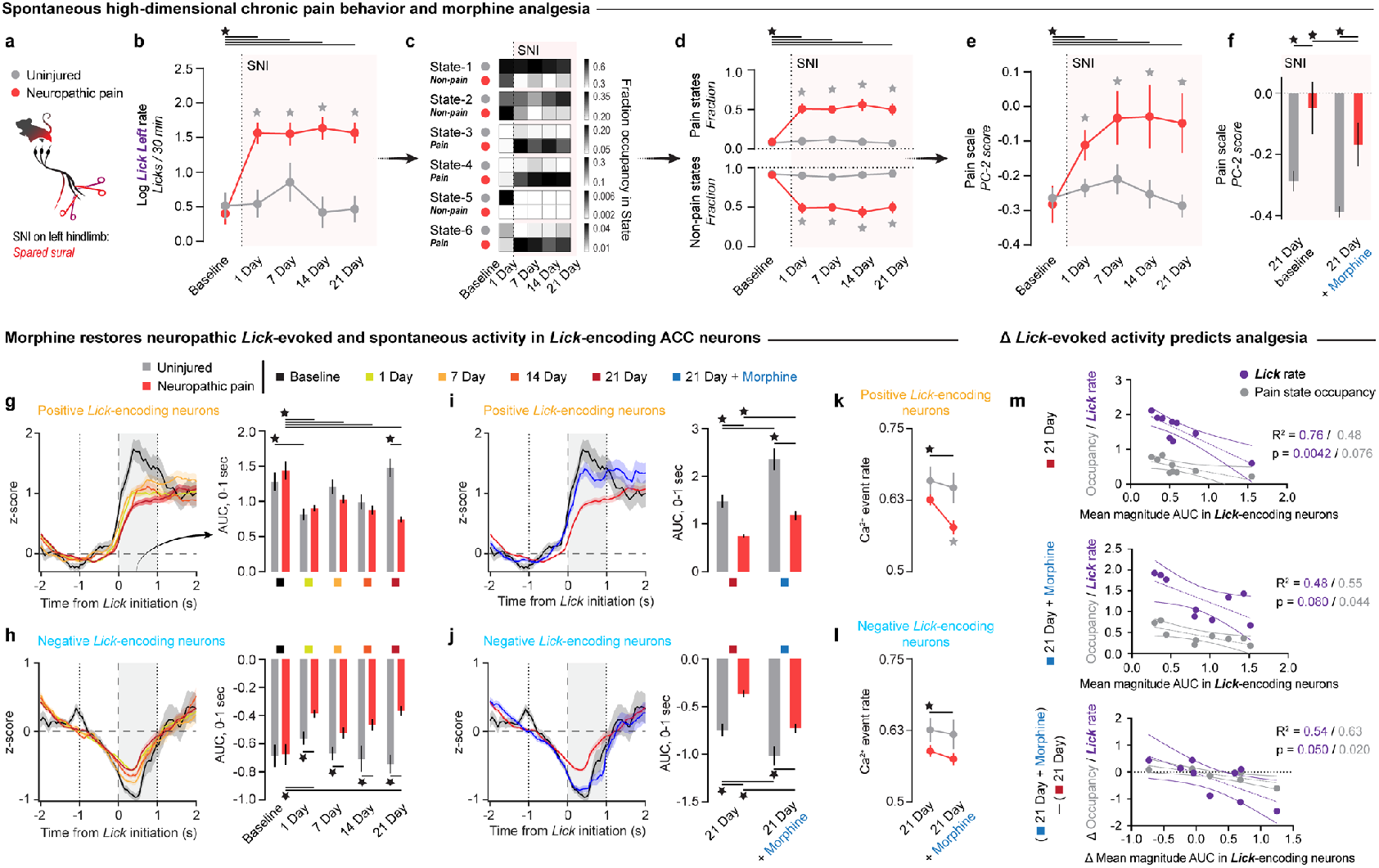
Morphine targets functionally compromised ACC neurons to relieve chronic pain. **(a)** SNI protocol for chronic neuropathic pain. **(b)** Log-transformed rate of spontaneous licking at the injury limb at -1, 1, 7, 14, and 21 days post-SNI (red, n=9) *vs*. uninjured controls (gray, n=9; 2-Way RM ANOVA + Tukey: *p*interaction = 0.0036). **(c)** Heatmap of average state occupancy. **(d)** Occupancy of pain (top) and non-pain (bottom) states in SNI and uninjured control mice (2-way RM ANOVA +Tukey: *p*interaction < 0.0001). **(e)** AMPS score (state PC2) score in SNI and uninjured control mice before and after SNI or anesthesia (2-way RM ANOVA + Tukey: *p*interaction = 0.0089). **(f)** AMPS score (state PC2) score in SNI and uninjured control mice three weeks post-SNI or no-injury, before and after morphine (0.5 mg/kg, i.p; 2-way RM ANOVA +Tukey: *p*injury = 0.0085, *p*treatment = 0.0041). **(g,h)** *Left*: Lick-evoked activity in positive (g) and negative (h) *p*Lick neurons, respectively, before (black) and after SNI (warm color gradient, yellow = 1 day post-SNI, red = 3 weeks post-SNI). *Right*: Area under the curve of lick-evoked activity (0-1s post-onset) in SNI (red) and uninjured (gray) control mice (2-way ANOVA + Tukey: *p*Positive, interaction < 0.0001, *p*Negative, interaction = 0.021). **(i,j)** *Left*: Same as (g,h left) visualizing lick-evoked activity in baseline (black), three weeks post-SNI (red), and three weeks post-SNI + morphine (blue) in SNI mice. *Right*: Same as (g,h) right comparing lick-evoked activity three weeks post-SNI *vs*. uninjured controls (2-way ANOVA + Tukey: *p*Positive, interaction = 0.035, *p*Negative, treatment < 0.0001, *p*Negative, injury < 0.0001). **(k,l)** Spontaneous calcium event rate in positive (k) and negative (l) *p*Lick neurons, before and after morphine (2-Way ANOVA + Tukey: *p*Positive, treatment = 0.0023, *p*Negative, treatment = 0.0124). **(m)** Linear regression predicting log-transformed lick rate (purple) and pain state occupancy (gray) from the average magnitude of lick-evoked activity three weeks post-SNI (*top*), after morphine (*middle*), and change between sessions (*bottom*; Bonferroni-corrected p-values displayed). ⋆ = P < 0.05. Bars, lines, or dots = mean; error bars and shaded areas = SEM. See ***Supplementary Table 3*** for statistics.

Here, *p*Lick neurons were identified using the same approach as in **Fig. 2**. Consistent with prior findings^80^, SNI impaired decoding accuracy of sensory stimuli, behaviors, and latent states relative to baseline and uninjured controls (**Fig. S10e–g, S13a–h, Supp. Table 3 Rows 14-17**). This impairment extended to *p*Lick neurons: SNI immediately and persistently blunted lick-evoked responses in both positive and negative *p*Lick neurons compared to baseline and control mice (**Fig. 3g,h, S14, Supp. Table 3 Rows 5-6**). Notably, morphine reversed these SNI-induced deficits in *p*Lick activity at lick onset, and further enhanced *p*Lick responses in uninjured controls (**Fig. 3i,j, Supp. Table 3 Rows 7-8**). SNI also reduced single-cell selectivity for licking in both positive and negative *p*Lick neurons, which was restored by morphine (**Fig. S13i,j, Supp. Table 3 Rows 18-21**). At the three-week time point, morphine suppressed activity in positive *p*Lick neurons in SNI mice and in negative *p*Lick neurons across both groups, mirroring the state-dependent and -independent effects observed in the acute pain model (**Fig. 3k,l, Supp. Table 3 Rows 22-25**). The magnitude of lick-evoked responses in *p*Lick neurons predicted both lick rate and pain state occupancy within session. Most critically, increases in lick-evoked response magnitude following morphine administration predicted reductions in both licking behavior and time spent in pain states (**Fig. 3m, Supp. Table 3 Rows 11-13**).

Together, these results show that, as in acute pain, morphine produces analgesia in chronic pain by inhibiting spontaneous activity in positive *p*Lick neurons in an injury-dependent manner, and by suppressing both spontaneous and evoked activity in negative *p*Lick neurons. Given that SNI blunted lick-locked activity, morphine’s action ultimately restored the behavior- and state-discriminating properties of these ACC ensembles. Thus, systemic morphine relieves chronic pain affect by rescuing impaired activity dynamics in pain-tracking ACC neurons.

### A cell-type–specific gene therapy targeting opioid-sensitive ACC neurons

Given that morphine relieves chronic pain by modulating defined populations of ACC neurons—including injury-dependent inhibition of positive *p*Lick neurons and widespread suppression of negative *p*Lick neurons—we next sought to mimic these effects through a targeted gene therapy approach. Rather than systemic delivery of opioids, which carries significant risk of addiction and off-target effects, we aimed to develop a circuit- and cell-type– specific strategy for precision pain management^1^. Specifically, we designed a gene therapy to silence MOR–expressing neurons in the ACC using chemogenetic inhibition. To accomplish this, we engineered an AAV-packaged synthetic mouse MOR promoter (*MORp*) to drive expression of Gi-coupled inhibitory hM4 DREADD (Designer Receptors Exclusively Activated by Designer Drugs), which leverage endogenous transcription factors and molecular machinery to expression transgene cargo in MOR^+^ cell-types. The *MORp* sequence was derived from a 1.5 kb region upstream of the *Oprm1* transcription start site, based on conserved regulatory elements in both mouse and human promoter regions previously shown to drive selective expression in MOR^+^ cells^41^. This allowed us to restrict hM4 DREADD expression to MOR^+^ neurons within the ACC, enabling remote and reversible inhibition via the ligand deschloroclozapine (DCZ) ^81^.

We tested two AAV-delivered strategies for targeting these cells: (1) direct expression of hM4Di or a control fluorescent protein (eYFP) under the control of *MORp* (constructs: *MORp*-hM4Di, MORp-eYFP), and (2) an intersectional approach combining *pain*TRAP labeling of noxious stimulus–responsive neurons with *MORp*-dependent expression using a Cre_ON_/Flp_ON_ switch to restrict expression to MOR^+^ pain-activated neurons (**Fig. 4a**). Following viral incubation, FISH quantification confirmed that over 97% of ACC neurons expressing endogenous *Oprm1* mRNA also expressed *MORp*-driven *eYfp* mRNA, indicating high specificity of the synthetic promoter (**Fig. 4b**). All viral constructs yielded robust expression throughout ACC layers (**Fig. 4c, S15**), validating this strategy as a viable tool for selective neuromodulation of opioid-sensitive cortical ensembles.

**Fig. 4.**
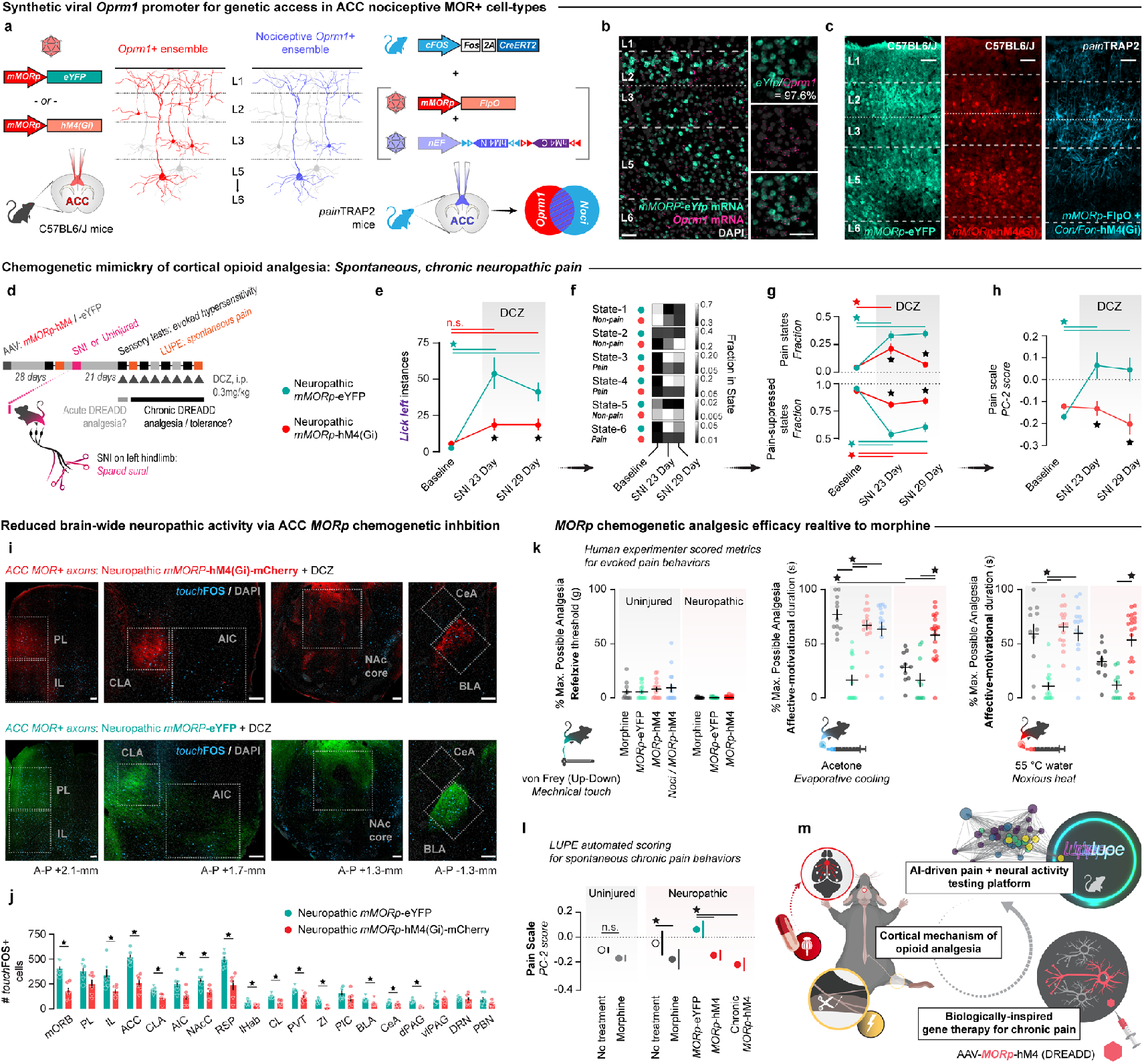
Precision neuromodulation of affective chronic pain via an ACC opioid cell-type specific gene therapy. **(a)** Strategies to deliver inhibitory chemogenetic actuators to ACC MOR+ neurons (Red: *MORp*-hM4) or nociceptive/MOR+ cell-types (Blue: *MORp*-FlpO + Cre_ON_/Flp_ON_-hM4 or *MORp*-DIO-iC++ in *pain*TRAP mice). **(b)** Co-expression of *MORp*-*eYfp* mRNA and endogenous *Oprm1* mRNA. **(c)** *MORp*-driven fluorophore or hM4 transgene expression across ACC layers. **(d)** Timeline of chronic neuropathic pain and DCZ exposure (0.3 mg/kg, i.p., once daily. Gray triangles) with LUPE and sensory testing (N=19 (10 male and 9 female) *MORp*-eYFP mice, N=30 (equal sexes) *MORp*-hM4 mice). **(e-h)** Effect of nerve injury and chemogenetic inhibition on spontaneous licking of the injured limb (e, *p*Interaction = 0.0012), occupancy of behavioral states (f), fraction of time spent in pain versus non-pain states (g, pain states *p*Interaction<0.0001, non-pain states *p*Interaction<0.0001), and AMPS scores (h, *p*Interaction = 0.0003) in *MORp*-hM4 *vs. MORp*-eYFP mice. 2-way RM-ANOVA + Tukey for all panels. **(i,j)** Brain-wide projections of ACC MOR+ axons expressing hM4 or eYFP (i) to assess neuropathic activity (*touch*FOS) within connected brain areas with or without ACC inhibition (j). *t*-Test, 2-tailed per region. **(k)** Analgesic efficacy for morphine (0.5 mg/kg, i.p) *vs. MORp*-eYFP, *MORp*-hM4, or nociceptive-*MORp*-hM4 + DCZ (0.3 mg/kg, i.p.) in uninjured and chronic neuropathic pain conditions. Effects on % Maximum Possible Analgesia on evoked mechanical thresholds (*left, p*Interaction = 0.0061), and evoked affective-motivational behaviors to acetone (*middle, p*Interaction <0.0001), and 55°C hot water (*right, p*Interaction P<0.0001). 1-way ANOVA + Tukey. **(l)** AMPS scores for spontaneous pain behaviors to assess analgesic efficacy of morphine *vs*. acute or chronic *MORp*-hM4 ACC inhibition in uninjured or chronic pain conditions. (Neuropathic left: 1-way ANOVA + Tukey, *p*Treatment =0.0085. Neuropathic right: 2-way ANOVA + Tukey, *p*Interaction =0.0003). **(m)** Overview of advancements: using deep-learning behavior tracking software to classify behavior states in pain, paired with calcium imaging we uncovered new cortical mechanisms of opioid analgesia, which informed our creation of an opioid cell-type gene therapy approach for chronic pain that mimics morphine analgesia with circuit-targeted precision. For detailed statistics, see ***Supplemental Table 4*** rows 1-10. ⋆ = P < 0.05. Errors bars = s.e.m.

### Targeted ACC gene therapy reduces spontaneous chronic pain without tolerance

Building on the successful targeting of ACC MOR^+^ neurons with AAV-*MORp*, we next asked whether chemogenetic inhibition of this population could provide sustained relief from chronic neuropathic pain without inducing tolerance. In mice three weeks after SNI, we began daily administration of the DREADD agonist DCZ (0.3 mg/kg, i.p.) for one week to assess: (1) whether DCZ activation of hM4Di-expressing MOR^+^ neurons reduced spontaneous and evoked pain behaviors, and (2) whether repeated chemogenetic inhibition would produce analgesic tolerance (**Fig. 4d**). Chemogenetic inhibition of ACC MOR^+^ neurons significantly reduced multiple measures of evoked affective-motivational pain behavior to 6°C acetone and 55°C water, with no evidence of tolerance. Spontaneous pain-related licking, occupancy of high-pain latent states, and AMPS scores were all reduced following a single dose of DCZ (SNI Day 23) and remained suppressed after one week of daily dosing (SNI Day 29; **Fig. 4e–h, S17, Supp Table 4 Rows 1-4**).

To further validate the analgesic potential of our cell-type–specific opioid gene therapy, we used a modified real-time place preference (RTPP) assay to assess pain relief–driven negative reinforcement (**Fig. S18, Supp. Table 5 Rows 3-4**). In painTRAP mice, we injected a Cre-dependent, *MORp*-driven inhibitory chloride channel (iC++) into the bilateral ACC to target MOR^+^ neurons activated by noxious stimuli. Fiber optic cannulas were implanted above the injection sites, and mice were assigned to uninjured or neuropathic pain (SNI) groups. At three weeks post-SNI, all mice underwent a 9-day RTPP protocol. A Day 1 pre-test assessed baseline chamber preference. Over the next seven days, daily 20-minute closed-loop sessions allowed mice to self-administer optogenetic inhibition via blue light (5 mW) triggered by entry into one chamber. On the final day, mice freely explored both chambers to assess conditioned place preference. Optogenetic inhibition of ACC MOR^+^ neurons had no effect on chamber preference in uninjured mice during conditioning or recall. In contrast, SNI mice showed progressively increased preference for the LED-paired chamber and robust conditioned recall—spending significantly more time in the chamber previously paired with inhibition. These findings indicate that silencing nociceptive MOR^+^ neurons in the ACC relieves ongoing spontaneous pain and is reinforcing only in the context of injury. Importantly, the lack of preference in uninjured animals suggests reduced risk of addiction-like liability. Together with our chemogenetic results, these data show that selective, chronic inhibition of ACC MOR^+^ neurons provide stable, state-dependent analgesia without tolerance or reinforcement in uninjured animals.

Given the anatomical connectivity of the ACC with key pain-processing regions—including the prelimbic and orbitofrontal cortices^82,83^, nucleus accumbens core^32,84^, insula and calustrum^85,86^, medial thalamus^87,88^, basolateral amygdala^42,66,89^, periaqueductal gray ^90^, and other regions—we hypothesized that inhibition of ACC MOR^+^ neurons may influence activity across broader nociceptive circuits. To test this, we collected brain tissue from the same chronic pain cohort after DCZ treatment and exposure to a standardized light touch stimulus (0.16g filament to the injured hindpaw). Using FOS expression as a proxy for neuropathic neural activation, we quantified the number of light touch-responsive neurons (*touch*Fos^+^ cells) across brain-wide projection targets of ACC MOR^+^ axons (**Fig. 4i**). We observed significant reductions in *touch*Fos^+^ cell counts in 14 of 19 brain regions downstream of ACC MOR^+^ projections, indicating that inhibition of this cortical ensemble can suppress widespread nociceptive responses throughout the brain (**Fig. 4j, Supp Table 4 Row 5**). These findings highlight the capacity of ACC MOR^+^ neurons to coordinate distributed pain networks and establish this targeted gene therapy as a circuit-level intervention for chronic pain relief without the limitations of opioid tolerance.

### Chemogenetic gene therapy-mediated analgesia mimics morphine to reduce acute and chronic pain

To evaluate the translational potential of our gene therapy, we compared its analgesic efficacy to morphine across a range of pain models and noxious stimuli. We quantified analgesia as *percent maximum analgesi*a(**Fig. 4k**, raw data in **Fig. S16**, Supp Table 4 Rows 6-8, 11-19) in mice expressing *MORp*-hM4Di, intersected *pain*TRAP/*MORp*-hM4Di, or *MORp*-YFP controls, and compared these to mice receiving systemic morphine (0.5 mg/kg, i.p.) in both uninjured and neuropathic pain conditions. Consistent with prior reports that low-dose morphine and ACC manipulation do not alter reflexive thresholds ^22,25,26^, none of the interventions significantly altered responses to mechanical von Frey filaments (**Fig. 4k, *left***). However, both morphine and DCZ (in hM4Di- or *pain*TRAP/*MORp*-hM4Di–expressing mice) produced significant analgesia to noxious acetone and 55°C water stimuli in uninjured animals compared to *MORp*-YFP controls (**Fig. 4k, *middle and right***). Importantly, in mice with SNI-induced neuropathic pain, DCZ administration in *MORp*-hM4Di mice significantly reduced responses to noxious heat and cold, whereas morphine failed (**Fig. 4k**). In addition, DCZ—whether delivered acutely or chronically over a week—was equally effective as morphine in reducing AMPS scores during chronic neuropathic pain (**Fig. 4l**), confirming its efficacy in modulating spontaneous, affective-motivational components of chronic pain. These findings demonstrate that our gene therapy not only mimics morphine’s analgesic effects in both acute and chronic settings but, in some cases, may provide superior efficacy against the affective-motivational components of chronic pain that morphine fails to address.

## Discussion

In summary (**Fig. 4m**), we developed LUPE, an automated behavioral analysis platform that quantifies naturalistic, spontaneous pain behaviors and latent state transitions in freely moving mice (**Fig. 1**). Using LUPE paired with single-cell calcium imaging, we identified specific ACC neural ensembles that track pain and are selectively modulated by morphine in both acute and chronic pain models (**Figs. 2–3**). These insights guided the design of a targeted, cell-type–specific gene therapy that mimics the analgesic effects of morphine by selectively inhibiting opioid-sensitive neurons in the ACC, providing durable and injured state-dependent pain relief without tolerance or reinforcement (**Fig. 4**). Importantly, ACC *MORp*-driven chemogenetic inhibition preserved sensory reflexes—allowing for necessary detection and localization of noxious stimuli—while reducing the affective-motivational aspects of chronic pain. We observed no reinforcement in uninjured animals with repeated *MORp*-iC++ optogenetic self-administration, suggesting a lower risk of addiction with this cortical opioidergic circuit-specific approach ^41^. Notably, we also found no evidence of analgesic tolerance with repeated *MORp*-chemogenetic or *MORp*-optogenetic activations. Prolonged *MORp*-hM4 signaling reversed affective hypersensitivity in neuropathic mice and reduced neuropathic activity across brain regions receiving direct innervation by ACC MOR+ circuits, indicating that disrupting persistent ACC-driven nociception can alleviate the chronic emotional burden of pain ^83,91^. Together, these findings establish a biologically inspired framework for precision neuromodulation of affective pain, advancing a new direction in circuit-targeted therapeutics.

Opioids have been used for millennia in medicinal cases to alleviate the unpleasantness of pain and make it “less bothersome” ^1,92^. Today, many prescription analgesics for moderate-to-severe pain are opioid derivatives, including morphine, codeine, oxycodone, hydrocodone, fentanyl, tramadol, methadone, hydromorphone, buprenorphine, and tapentadol ^4,93,94^. Despite their efficacy, these compounds carry fatal and addictive side effects, largely due to the widespread activation of opioid receptors in critical areas processing reward and respiration. This highlights the urgent need for novel, non-opioid therapies to address the Opioid Epidemic ^95,96^. While there have been significant reductions in opioid prescribing over the past decade, few safer alternatives for pain management have emerged, leaving millions of chronic pain patients with limited treatment options. Over recent decades, analgesic drug development aimed at molecular targets in peripheral sensory afferent neurons or spinal neural circuits has seen both clinical trial failures ^5^ and recent successes ^97^. These efforts are directed toward advancing non-opioid pain management. Different pain-relieving drugs do not equally interfere with the same underlying mechanisms or parts of the nervous system that contribute to pain ^98^—for example, non-steroidal anti-inflammatory drugs (NSAIDs) reduce cyclooxygenase production of prostaglandins, Nav1.7 and Nav1.8 sodium channel blockers reduce peripheral nociceptor transmissions to the spinal cord, or as we show here, morphine modulates cortical affective processes to reduce pain unpleasantness ^4,23,64,92^. Thus, identifying neural mechanisms that mediate opioid effects on unpleasantness are a critical step for advancing therapeutic discovery efforts aimed at developing more targeted, effective, and safer treatments^1^.

Perceived pain unpleasantness strongly correlates with fMRI activity in the human ACC ^8,16^. Patients with intractable chronic pain treated with surgical cingulotomy lesions do not report changes in pain perception, intensity discrimination, or reactions to momentary harmful stimuli ^17,18,99,100^. Rather, their attitude toward pain is modified, dissociating the negative valence from the experience of pain. In these cases, pain becomes a sensation rather than a threat ^19^. However, cingulotomy can result in difficulties sustaining attention and focus, as well as impairments in executive functions like response intention, generation, and persistence ^101,102^. The primary overall effect of cingulotomy is the attenuation of patients’ continual emotional and behavioral responses to their ever-present pain, leading to reduced attention switching to and rumination on pain. Like cingulotomy, exogenous opioids separate the sensation of pain from its negative valence and attention-drawing aspects ^23,49^. Reports from patients treated with ACC lesions and opioids suggest that relief from the affective symptoms of chronic pain is therapeutically meaningful. Acute pain engages MORs in the dorsal anterior cingulate and lateral prefrontal cortex, and the affective perception of pain is correlated with MOR availability in the ACC and other brain regions ^23,24,62,103,104^. Together, this evidence supports the role of MOR signaling within the ACC in altering the human perception of pain ^12^. The similarity between the affective mechanisms of morphine and cingulotomy in humans mirrors our preclinical results with the genetic tuning of MOR expression in the ACC and our AAV-*MORp*-driven therapy. We conclude that opioids, as well as our opioid mimicry therapy, inhibit cortical nociceptive functions to reduce the integration of negative valence information within the ACC, resulting in reduced aversive arousal and attentional processes that bias the selection of nocifensive behaviors.

Theories of ACC function center around highly specialized cognitive-affective functions— task engagement, motivation, error detection, attention allocation, value appraisal, and action selection. For example, value appraisal of sensory information may lead to the engagement of appropriate motor actions ^12,105^, which are critical in measured performance in cognitive tasks involving decision-making based on sensory-guided information ^106^. During a painful experience, the “task” can be defined as sustained motivational recuperative behaviors, and the “decision” as the selection to engage in protective and/or escape behaviors ^42^. We find that ACC nociception preferentially engages animals in affective-motivational pain attending and escape behaviors. This aligns with the ACC’s established role in amplifying task-relevant signals ^107^, especially under conflicting conditions ^108^ —such as balancing pain, ongoing behaviors, and analgesic effects. In acute pain, this circuitry may facilitate adaptive responses by enhancing nociceptive signals that promote protective behaviors like licking or rearing. In chronic pain, where pain persists beyond tissue injury, this same circuitry may become maladaptive, sustaining heightened emotional and behavioral responses to ongoing pain. Specifically, in line with ethological theory, the detailed behavioral measurements and dynamic transitions quantified from our LUPE experiments indicate that these volitional behaviors are composed of observable, stereotyped behavioral motifs that construct larger behavior repertoires, relying on affective and motivation-based signaling in the ACC during pain experiences ^109^.

Developing precise, accurate, and reliable preclinical methods to capture behavioral biomarkers that reflect the complexity of pain is essential for identifying treatments that engage cognitive-emotional pain circuits. Traditional rodent measures—such as reflexive withdrawal thresholds—provide valuable information about stimulus detection but fail to capture the affective-evaluative dimensions of pain that shape motivational behavior. In this context, deep learning and machine vision have become powerful tools for automated, high-resolution behavioral analysis, enabling unbiased quantification of pain-related dynamics and responses to analgesics or neural interventions ^110–112^. LUPE advances this field by combining expert-guided semi-supervised classification with unsupervised clustering to robustly identify a core set of interpretable, ethologically grounded behaviors. Unlike fully unsupervised methods such as MoSeq^113^—which rely on fine-scale pose dynamics to define micro-behaviors that may not map cleanly onto cognitive-affective states^111^—LUPE explicitly captures high-level, volitional behaviors like paw licking that are historically linked to pain. By modeling transitions between these behaviors as Markov processes and clustering them using a k-means framework, LUPE identifies latent behavioral states that encode injury and analgesia, including a reproducible pain-associated state (*Pain State-4*). From these latent dynamics, we derive the mouse *Affective-Motivational Pain Scale* (AMPS)—a continuous, data-driven metric that tracks affective-motivational aspects of pain and their relief with opioids. Together, these features make LUPE a significant leap forward in behavioral phenotyping for pain neuroscience. It not only enables standardized and high-throughput behavioral analysis but also provides a new lens for decoding internal pain states—advancing our ability to evaluate therapeutics that target the affective burden of pain.

In total, our study demonstrates the therapeutic potential of targeting defined cortical ensembles for precision pain relief. By leveraging a synthetic MOR promoter to drive chemogenetic inhibition in opioid-sensitive ACC neurons, we successfully mimicked the analgesic effects of morphine while avoiding its sensory, reinforcing, and tolerance-related side effects. This cell-type– specific strategy offers a new direction for pain management— one that could bypass the systemic risks of traditional opioids by modulating unique dimensions of pain perception at its cortical origin. For translational neuromodulation, our gene therapy could be adapted for non-invasive delivery using focused ultrasound blood-brain barrier opening^114^, to access and control multiple pain-encoding opioidergic neurons simultaneously in cortical and subcortical circuits. Thus, by integrating behavioral state modeling, neural ensemble identification, and circuit-targeted intervention, our work provides a framework for biologically informed, precision-based pain therapeutics. Selectively targeting the ACC opioidergic circuits via *MORp*-based gene therapies hold the promise of offering safer and more effective alternatives to conventional pain treatments, ultimately advancing precision medicine in pain management.

## Acknowledgements

This work was supported by: National Institute of Health (NIH) awards NIGMS DP2GM140923 (G.C.), NIDA R00DA043609 (G.C.), NIDA R01DA054374 (G.C., K.B.), NINDS R01NS130044 (G.C., K.B.), NIDA R01DA056599 (G.C., K.B.), NIDA R21DA055846 (B.C.R.), NIDA F31DA062445 (C.S.O), NIDA F32DA053099 (N.M.M), NIDA F32DA055458 (B.A.K.), NIDA F31DA057795 (L.M.W.), NINDS F31NS125927 (J.A.W.), NIDA T32DA028874 (G.S.), NINDS RF1NS126073 (M.R.B.), the Whitehall Foundation (G.C.), the Howard Hughes Medical Institute (K.D.). We thank the following for shared reagents and technical assistance: Julie Blendy, Angelina Heyeler, Lauren Marconi, Joseph Stzyuncki, Morgan Kindel, and the Penn University Laboratory Animal Resources (ULAR) veterinarians and husbandry staff. We thank Stanford University “Cracking the Neural Code” Program and Gene Vector and Virus Core for custom viruses.

## Competing interests

G.C, K.D., C.R. and G.J.S. are inventors on a provisional patent application through the University of Pennsylvania and Stanford University regarding the custom sequences used to develop, and the applications of synthetic opioid promoters (patent application number: 63/383,462 462 ‘Human and Murine *Oprm1* Promotes and Uses Thereof’).

## Code, data and materials availability

All code for LUPE analyses of behavior and paired calcium imaging can be found at: https://github.com/justin05423/LUPE-2.0-AnalysisPackage. All new mu-opioid receptor promoter viruses (AAV-MORp) will be available for academic use from the Stanford Gene Vector and Virus Core (https://neuroscience.stanford.edu/research/neuroscience-community-labs/gene-vector-and-virus-core) and/or by contacting the lead authors. Upon manuscript peer-review publication, all data (e.g. 10X snRNAseq, LUPE video files, raw TIFF images from histology, etc.) will be unembargoed at Zenodo and GEO.

## Author contributions

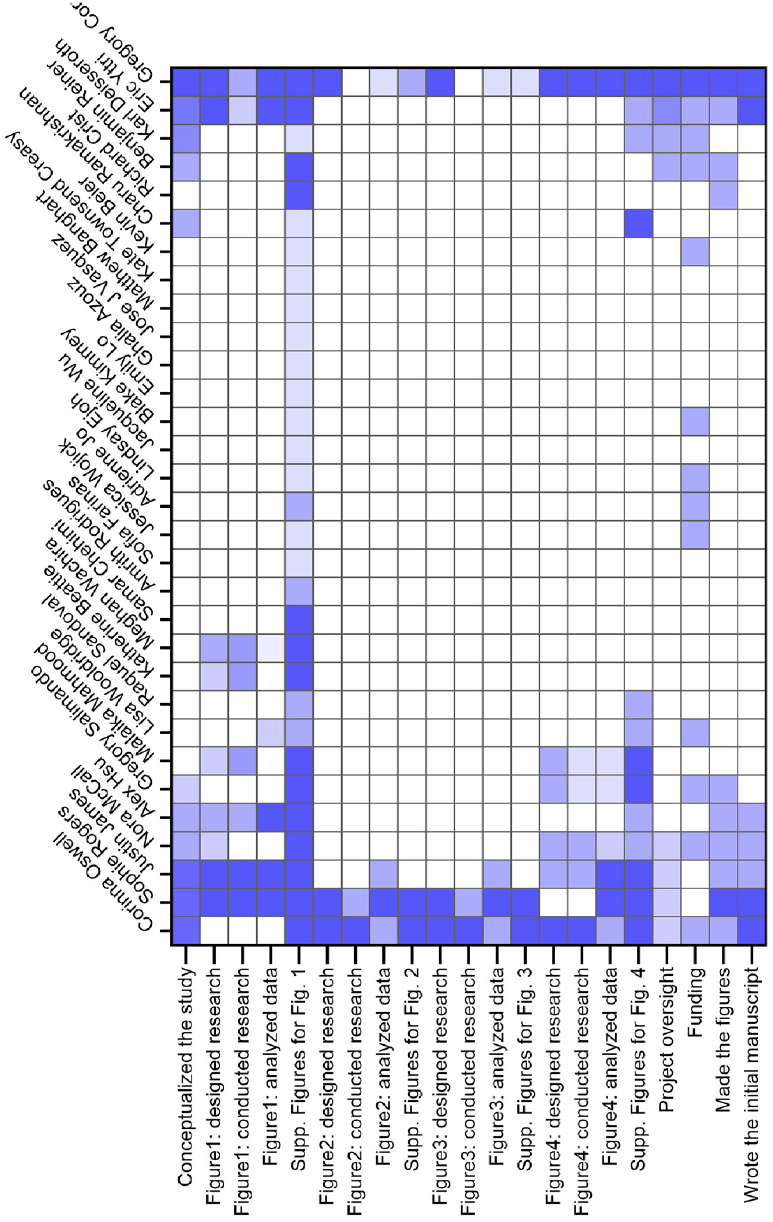

## Supplementary Materials

### Methods

#### Sample sizes

Sample size was not statistically determined prior to the studies. Rather, the group sizes were based off published literature for the type of manipulation (i.e. chemogenetics, site-specific genetic knockout, pharmacology) and measured outcome (e.g. pain behaviors) published in the field and/or by the authors involved (examples include PMID 37704594, 30655440, 34646022, 25948274). Sample sizes for all studies are included in the figure legends.

#### Data Exclusions

For all imaging and behavioral studies, virus injected animals with either little or no evidence of viral transduction and/or incorrect viral targeting were excluded from any final analyses. No other mice or data points were excluded across analyses.

#### Replication

For many of the behavior studies, multiple cohorts were used due to the large number of animals in the final group sizes. All behavior results were consistent and replicated across cohorts. Individual data points or lines are included and indicate consistent trends across many mice in each behavior study.

#### Blinding

Mice were randomly assigned into control or experimental groups to the best of the experimenter’s abilities, with counterbalancing for age and sex as needed. In most of the included studies, the experimental and control groups only differ in the type of virus intracranially infused. The surgical protocol for all mice was identical in the amount, wait time, and location of the intracranial injection. Each surgery day was randomly assigned as a control/experimental surgery date and the according mice from the pre-determined groups underwent surgery that day. GRIN lens and fiber placements and viral spread maps are included in the supplement to demonstrate the similarity of the injection protocol and outcome. Once experimental/control groups were formed to comprise the studies cohort of mice, the cohort underwent all behavioral testing concurrently and experimenters were blinded. After analyses were complete, the experimenters were unblinded.

#### Animals

All experimental procedures were approved by the Institutional Animal Care and Use Committee of the University of Pennsylvania and performed in accordance with the US National Institutes of Health (NIH) guidelines. Male and female mice aged 2-5 months were housed 2-5 per cage and maintained on a 12-hour reverse light-dark cycle in a temperature and humidity-controlled environment. All experiments were performed during the dark cycle. Mice had ad-libitum food and water access throughout experiments. For behavioral, anatomical, and transcriptomic experiments, we utilized *Fos*-FOS-2A-iCreER^T2^ or “TRAP2” mice (Fos^tm2.1(icre/ERT2)Luo)Luo^/Jackson Laboratory, stock #030323) ^115^ bred to homozygosity, C57BL/6J wild type mice, or Oprm1^Cre/Cre^ mice, and Oprm1^fl/fl^ mice. Additional anatomical experiments utilized TRAP2 mice crossed with *Ai9* (B6.Cg-Gt(ROSA)26Sor^tm9(CAG-tdTomato)Hze^/J) reporter mice that express a tdTomato fluorophore in a Cre-dependent manner (“TRAP2:tdTomato”) purchased from Jackson Laboratory, stock #007909 bred to heterozygosity or homozygosity for both genes.

#### Mouse μ-Opioid Receptor promoter (MORp)

*MORp is* a 1.5 Kb segment selected and amplified from mouse genomic DNA using cgcacgcgtgagaacatatggttggacaaaattc and ggcaccggtggaagggagggagcatgggctgtgag as the 5’ and 3’ end primers respectively. All *MORp* plasmids were constructed on an AAV backbone by inserting either the *MORp* promoter ahead of the gene of interest (i.e., iC++-eYFP) using M1uI and AgeI restriction sites. Every plasmid was sequence verified. Next, all AAVs were produced at the Stanford Neuroscience Gene Vector and Virus Core. Genomic titer was determined by quantitative PCR of the WPRE element. All viruses were tested in cultured neurons for fluorescence expression prior to use *in vivo*.

#### Viral Vectors

All viral vectors were either purchased from Addgene.org, or custom designed and packaged by the authors as indicated. All AAVs were aliquoted and stored at -80°C until use. The following AAVs were used:

- AAV1-*MORp*-DIO-iC++ (titer: 1.35e12 vg/mL)
- AAV1-*MORp*-hM4(Gi)-mCherry (titer: 1.7e12 vg/mL)
- AAV1-*MORp*-eYFP (titer: 1.0e12 vg/mL),
- AAV1-*MORp*-Flpo (titer: 9.2e11 vg/mL)
- AAV5-*nEF*-Con/Fon-hM4Di-mCherry (7.4e12 vg/mL)
- AAVDJ-*hSyn*-DIO-mCh-2A-MOR (titer: 1.13e12 vg/mL)
- AAV9-*hSyn*-HI.eGFP-Cre (Addgene 105540-AAV9; titer: 5.44e11 vg/mL)
- AAV9-*hSyn*-GFP (Addgene 50465-AAV9; titer: 1.9e11 vg/mL)
- AAV5-*hSyn*-DIO-EGFP (Addgene 50457-AAV5; titer: 1.3e12 vg/mL)
- AAV9-*hSyn*-jGCaMP8m-WPRE (Addgene 162375-AAV9; titer: 1.2e12 vg/mL)

#### Stereotaxic surgery

Adult mice (∼8 weeks of age) were anesthetized with isoflurane gas in oxygen (initial dose = 5%, maintenance dose = 1.5%), and fitted into WPI or Kopf stereotaxic frames for all surgical procedures. 10 µL Nanofil Hamilton syringes (WPI) with 33 G beveled needles were used to intracranially infuse AAVs into the ACC. The following coordinates were used, based on the Paxinos mouse brain atlas, to target these regions of interest: ACC (from Bregma, AP: -1.50 mm, ML: ± 0.3 mm, DV: −1.5 mm). Mice were given a 3– 8-week recovery period to allow ample time for viral diffusion and transduction to occur. For all surgical procedures in mice, meloxicam (5 mg/kg) was administered subcutaneously at the start of the surgery, and a single 0.25 mL injection of sterile saline was provided upon completion. All mice were monitored and given meloxicam for up to three days following surgical procedures.

#### Chronic neuropathic pain model

As described previously ^116^, to induce a chronic pain state, we used a modified version of the Spared Nerve Injury (SNI) model of neuropathic pain. This model entails surgical section of two of the sciatic nerve branches (common peroneal and tibial branches) while sparing the third (sural branch). Following SNI, the receptive field of the lateral aspect of the hindpaw skin (innervated by the sural nerve) displays hypersensitivity to tactile and cool stimuli, eliciting pathological reflexive and affective-motivational behaviors (allodynia). To perform this peripheral nerve injury procedure, anesthesia was induced and maintained throughout surgery with isoflurane (4% induction, 1.5% maintenance in oxygen). The left and/or hind leg was shaved and wiped clean with alcohol and betadine. We made a 1-cm incision in the skin of the mid-dorsal thigh, approximately where the sciatic nerve trifurcates. The biceps femoris and semimembranosus muscles were gently separated from one another with blunt scissors, thereby creating a <1-cm opening between the muscle groups to expose the common peroneal, tibial, and sural branches of the sciatic nerve. Next, ∼2 mm of both the common peroneal and tibial nerves were transected and removed, without suturing and with care not to distend the sural nerve. The leg muscles are left unsutured and the skin was closed with tissue adhesive (3M Vetbond), followed by a Betadine application. During recovery from surgery, mice were placed under a heat lamp until awake and achieved normal balanced movement. Mice were then returned to their home cage and closely monitored over the following three days for well-being.

#### Targeting Recombination in Active Populations (TRAP) protocols

##### *pain*TRAP

*pain*TRAP induction was performed similarly to previously described ^116^. We habituated mice to a testing room for two to three consecutive days. During these habituation days, no nociceptive stimuli were delivered, and no baseline thresholds were measured (i.e. mice were naïve to pain experience before the TRAP procedure). We placed individual mice within red plastic cylinders (∼9-cm D), with a red lid, on a raised perforated, flat metal platform (61-cm x 26-cm). The experimenters sat in the testing room for the thirty minutes of habituation; this was done to mitigate potential alterations to the animal’s stress and endogenous antinociception levels. To execute the TRAP procedure, we placed mice in their habituated cylinder for 30 min, and then a 55°C water droplet was applied to the central-lateral plantar pad of the left hindpaw) once every 30 s over 10 min. Following the water stimulations, the mice remained in the cylinder for an additional 60 min before injection of 4-hydroxytamoxifen (4-OHT, 40 mg/kg in vehicle; subcutaneous). After the injection, the mice remained in the cylinder for an additional 4 hours to match the temporal profile for c-FOS expression, at which time the mice were returned to the home cage.

##### *Home-cage*TRAP

*Home-cage*TRAP induction was performed without habituation. At least 2 hours into the dark cycle, mice were gently removed from their home cages. Mice were then injected with 4-OHT (40 mg/kg in vehicle; subcutaneous) and returned to their home cages.

#### Immunohistochemistry

Animals were anesthetized using FatalPlus (Vortech) and transcardially perfused with 0.1 M phosphate buffered saline (PBS), followed by 10% normal buffered formalin solution (NBF, Sigma, HT501128). Brains were quickly removed and post-fixed in 10% NBF for 24 hours at 4 °C, and then cryo-protected in a 30% sucrose solution made in 0.1 M PBS until sinking to the bottom of their storage tube (∼48 h). Brains were then frozen in Tissue Tek O.C.T. compound (Thermo Scientific), coronally sectioned on a cryostat (CM3050S, Leica Biosystems) at 30 μm or 50 μm and the sections stored in 0.1 M PBS. Floating sections were permeabilized in a solution of 0.1 M PBS containing 0.3% Triton X-100 (PBS-T) for 30 min at room temperature and then blocked in a solution of 0.3% PBS-T and 5% normal donkey serum (NDS) for 2 hours before being incubated with primary antibodies (1^°^Abs included: chicken anti-GFP [1:1000, Abcam, ab13970], guinea pig anti-FOS [1:1000, Synaptic Systems, 226308], rabbit anti-FOS [1:1000, Synaptic System, 226008], rabbit anti-DsRed [1:1000, Takara Bio, 632496]; prepared in a 0.3% PBS-T, 5% NDS solution for ∼16 h at room temperature. Following washing three times for 10 min in PBS-T, secondary antibodies (2^°^Abs included: Alexa-Fluor 647 donkey anti-rabbit [1:500, Thermo Scientific, A31573], Alexa-Fluor 488 donkey anti-chicken [1:500, Jackson Immuno, 703-545-155], Alexa-Fluor 555 donkey anti-rabbit [1:500, Thermo Scientific, A31572] Alexa-Fluor 647 donkey anti-guinea pig [1:500, Jackson Immuno, 706-605-148], prepared in a 0.3% PBS-T, 5% NDS solution were applied for ∼2h at room temperature, after which the sections were washed again three times for 5 mins in PBS-T, then again three times for 10 min in PBS-T, and then counterstained in a solution of 0.1 M PBS containing DAPI (1:10,000, Sigma, D9542). Fully stained sections were mounted onto Superfrost Plus microscope slides (Fisher Scientific) and allowed to dry and adhere to the slides before being mounted with Fluoromount-G Mounting Medium (Invitrogen, 00-4958-02) and cover slipped.

#### Fluorescence *in situ* hybridization

Animals were anesthetized using isoflurane gas in oxygen, and the brains were quickly removed and fresh frozen in O.C.T. using Super Friendly Freeze-It Spray (Thermo Fisher Scientific). Brains were stored at −80° C until cut on a cryostat to produce 16 μm coronal sections of the ACC. Sections were adhered to Superfrost Plus microscope slides, and immediately refrozen before being stored at −80° C. Following the manufacturer’s protocol for fresh frozen tissue for the V2 RNAscope manual assay (Advanced Cell Diagnostics), slides were fixed for 15 min in ice-cold 10% NBF and then dehydrated in a sequence of ethanol serial dilutions (50%, 70%, and 100%). Slides were briefly air-dried, and then a hydrophobic barrier was drawn around the tissue sections using a Pap Pen (Vector Labs). Slides were then incubated with hydrogen peroxide solution for 10 min, washed in distilled water, and then treated with the Protease IV solution for 30 min at room temperature in a humidified chamber. Following protease treatment, C1 and C2 cDNA probe mixtures specific for mouse tissue were prepared at a dilution of 50:1, respectively, using the following probes from Advanced Cell Diagnostics: *Oprm1* (C1, 315841), *Slc17a7* (C3, 416631), *Fos* (C4, 316921). Sections were incubated with cDNA probes (2 h) and then underwent a series of signal amplification steps using FL v2 Amp 1 (30 min), FL v2 Amp 2 (30 min) and FL v2 Amp 3 (15 min). 2 min of washing in 1x RNAscope wash buffer was performed between each step, and all incubation steps with probes and amplification reagents were performed using a HybEZ oven (ACD Bio) at 40° C. Sections then underwent fluorophore staining via treatment with a serious of TSA Plus HRP solutions and Opal 520, 570, and 620 fluorescent dyes (1:5000, Akoya Biosystems, FP1487001KT, FP1495001KT). All HRP solutions (C1-C2) were applied for 15 min and Opal dyes for 30 min at 40° C, with an additional HRP blocker solution added between each iteration of this process (15 min at 40° C) and rinsing of sections between all steps with the wash buffer. Lastly, sections were stained for DAPI using the reagent provided by the Fluorescent Multiplex Kit. Following DAPI staining, sections were mounted, and cover slipped using Fluoromount-G mounting medium and left to dry overnight in a dark, cool place. Sections from all mice were collected in pairs, using one section for incubation with the cDNA probes and another for incubation with a probe for bacterial mRNA (dapB, ACD Bio, 310043) to serve as a negative control.

#### Imaging and Quantification

All tissue was imaged on a Keyence BZ-X all-in-one fluorescent microscope at 48-bit resolution using the following objectives: PlanApo-λ x4, PlanApo-λ x20 and PlanApo-λ x40. All image processing prior to quantification was performed with the Keyence BZ-X analyzer software (version 1.4.0.1). Quantification of neurons expressing fluorophores was performed via manual counting of TIFF images in Photoshop (Adobe, 2021) using the Counter function or using HALO software (Indica Labs), which is a validated tool for automatic quantification of fluorescently-labeled neurons in brain tissue ^117–119^. Counts were made using 20X magnified z-stack images of a designated regions of interest (ROI). For axon density quantification, immunohistochemistry was performed to amplify the signal and visualize ACC axons throughout the brain in 50 μm tissue free floating slices as described above. Areas with dense axon innervation were identified using 4X imaging. Areas implicated in emotion and nociception were selected for additional 20X imaging with Z-stacks. These regions of interest (ROIs) were initially visualized at 20X to determine the region with the highest fluorescence. The exposures for FITC and CY3 were adjusted to avoid overexposed pixels for the brightest area. This exposure was kept consistent for all slices for an individual mouse. For an individual ROI, one slice per mouse was included.

We used HALO software for all quantifications. One representative 16 μm slice containing ACC (selected from 1.1-1.3mm anterior of bregma) was quantified per mouse, using HALO Image Analysis software (Indica Labs). The borders for left and right ACC, Cg1, Cg2, L1, L2/3, L5, L6a, and L6b were hand-drawn as individual annotation layers, using the Allen Brain Reference Atlas as a guide. Slices were visually inspected for damage, dust or other debris, and bound probe, and these areas were manually excluded from their respective annotation layers. Colocalization of nuclei (DAPI) with *Oprm1, Fos*, and *Vglut* mRNA puncta was automatically quantified using the FISH module (version 3.2.3) and traditional nuclear segmentation. Setting parameters were optimized by comparing performance across 6 slices, randomly selected across experimental groups, and confirming proper detection by visual inspection. Identical parameters were applied across all slices in the data set.

#### Drugs and Delivery

For chemogenetic studies, deschloroclozapine (DCZ dihydrochloride, water soluble; HelloBio, HB9126) was delivered i.p. at a dose of 0.3 mg/kg body weight. For Oprm1 knockout, re-expression, and miniscope testing, morphine sulfate (Hikma) was delivered acutely i.p. at a dose of 0.5 mg/kg body weight

#### Human-scored Behavioral tests

All experiments took place during the dark phase of the cycle (0930 hour to 1830 hour). Group and singly housed mice were allowed a 1–2-week acclimation period to housing conditions in the vivarium prior to starting any behavior testing. Additionally, three to five days before the start of testing, mice were handled daily to help reduce experimenter-induced stress. On test days, mice were brought into procedure rooms ∼1 hour before the start of any experiment to allow for acclimatization to the environment. Mice were provided food and water *ad libitum* during this period. For multi-day testing conducted in the same procedure rooms, animals were transferred into individual “home away from home” secondary cages ∼1 hour prior to the start of testing and were only returned to their home cages at the end of the test day. All testing and acclimatization were conducted under red light conditions (< 10 lux), with exposure to bright light kept to a minimum to not disrupt the animals’ reverse light cycle schedule. Equipment used during testing was cleaned with a 70% ethanol solution before starting, and in between, each behavioral trial to mask odors and other scents.

##### Sensory testing for pain affective-motivational and nociceptive reflex behavioral assays

To evaluate responses to acute stimuli, animals were placed in transparent red cylinders placed on top of a metal hexagonal-mesh floored platform. Stimuli were applied to the underside of the left plantar hind paw. This process was repeated for a total of 10 applications, with each droplet applied at a 1 min interval. Animals were continuously recorded by a web camera positioned to face the front of the cylinder in which the animal was housed, and the time spent attending to the affected paw was quantified for up to 30 sec after the stimulation.

To evaluate mechanical reflexive sensitivity, we used a logarithmically increasing set of 8 von Frey filaments (Stoelting), ranging in gram force from 0.07- to 6.0-g. These filaments were applied perpendicular to the plantar hindpaw with sufficient force to cause a slight bending of the filament. A positive response was characterized as a rapid withdrawal of the paw away from the stimulus within 4 seconds. Using the Up-Down statistical method, the 50% withdrawal mechanical threshold scores were calculated for each mouse and then averaged across the experimental groups ^116^. To evaluate affective-motivational responses evoked by thermal stimulation ^116^, we applied either a single, unilateral 55°C drop of water or acetone (evaporative cooling) to the left hindpaw, and the duration of attending behavior was collected for up to 30 s after the stimulation. Responses to the noxious stimulus was also tested following acute i.p. administration of morphine (0.5 mg/kg body weight) or DCZ (0.3 mg/kg body weight). After injection, animals were placed back in their home away from home cages for 30 mins to allow for complete absorption of the drug. Hot water hind paw stimulation testing then proceeded as described above in the naïve condition.

Additionally, we used an inescapable hotplate set to 50° C. The computer-controlled hotplate (6.5 in x 6.5 in floor, Bioseb) was surrounded by a 15 in high clear plastic chamber and two web cameras were positioned at the front or side of the chamber to continuously record animals to use for post hoc behavioral analysis. For the tests conducted for chemogenetic or pharmacology studies, mice were administered morphine or DCZ 30 min prior to the start of behavior testing to allow for complete absorption of the drug and previous sensory testing ^120^. Mice were gently placed in the center of the hotplate floor and removed after 60 seconds.

##### Maximum Possible Analgesia effect (%MPA) Calculation

The Maximum Possible Analgesia (%MPA) metric quantifies how much a pain-related behavior is reduced following drug administration, relative to both the animal’s baseline response and the maximum behavioral response for that assay. This normalization enables meaningful comparisons across animals with different baseline sensitivity levels.

It is calculated as:

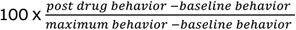

This normalization allows for comparisons across animals with different baseline response levels. For example: In the von Frey Up-Down test, the *maximum behavior* for withdrawal threshold is 6.0 grams (the highest filament force that would indicate the maximal amount of analgesia post-drug), and the minimum threshold is 0.007 grams. While for affective-motivational responses to thermal stimuli, behavioral responses such as attending or escape are measured over a 60-second window. Here, the minimum behavior response time is 0.0 seconds, and the maximum behavior is capped at 30 seconds (the trial duration); in this case, maximum analgesia would be to not respond at all the noxious stimulus, and thus the maximum behavior in the %MPA formula is 0.0 seconds. This approach ensures consistent scaling of behavioral change across experiments and conditions. This formula provides a normalized score ranging from 0% (no analgesia) to 100% (complete analgesia). The definition of “maximum behavior” depends on the behavioral test and reflects the highest measurable response in the absence of any analgesia, while the “baseline behavior” is typically the pre-drug measurement for that animal. To demonstrate, here are examples of use:

Mechanical Sensitivity (von Frey Up-Down test): In this assay, paw withdrawal threshold is measured in grams of filament force. The maximum behavior is defined as 6.0 grams (the highest force filament, indicating no withdrawal and thus complete analgesia), and the minimum threshold is 0.007 grams (the lowest measurable force, often reflecting high sensitivity). A mouse with a baseline threshold of 0.4 grams that increases to 3.0 grams after drug administration would have:

- Baseline withdrawal threshold = 0.4 grams
- Post-drug threshold = 3.0 grams
- Maximum threshold (no pain) = 6.0 grams
- Formula:
  - %MPA = 100 x (3.0 - 0.4) / (6.0 - 0.4)
  - %MPA = 100 × 2.6 / 5.6
  - %MPA = 46.4%

Affective-Motivational Responses (e.g., to thermal stimuli): In this test, behavioral responses such as licking or escape are measured over a 30-second trial window. The maximum behavior is defined as 30 seconds (a full response indicating high pain sensitivity), and the minimum behavior is 0 seconds (no response, indicating full analgesia). For these tests, because lower values reflect reduced pain, the formula is adapted by reversing the direction:

- Baseline response time = 20.0 seconds
- Post-drug response time = 5.0 seconds
- Minimum response = 0.0 seconds (no response = complete analgesia)
- Formula:
  - %MPA = 100 x (20.0 - 5.0) / (20.0 - 0.0)
  - %MPA = 100 × 15.0 / 20.0
  - %MPA = 75.0%

This approach ensures consistent, interpretable scaling of drug-induced behavioral changes across assays and experimental conditions.

#### LUPE – Light aUtomated Pain Evaluator acquisition and analysis software

##### Video Acquisition

Behavioral videos were recorded using a Basler ace UacA2040-120um camera at a fixed resolution of 768 (width) by 770 (height) pixels. Imaging parameters were standardized, with gain set to 10.0 dB and gamma at 2.0. Exposure mode was timed with an exposure duration of 1550 ms per frame, triggered at the start of each frame. Videos were captured at a consistent frame rate of 60 frames per second (fps) at maximum quality, with a recording buffer size of 128 frames. Frames were stored every 16 ms to ensure high temporal resolution of behavioral sequences.

##### Pose Estimation via DeepLabCut

Assigning 2D markerless pose estimation of mice within LUPE was achieved through the DeepLabCut (DLC) program (version 2.3.5-8). DLC was favored for this purpose because of its capability to track body-points at high confidence when animals perform diverse behaviors as well as its ability to accurately report if a body part is visible in a given frame. Its extensive toolkit, documentation, and forums allow flexible user input and manipulation when creating models.

Body-points considered for assigning pose in the LUPE-DLC Model were based on the clarity and frequency of appearance, involvement in behavior sequences, and prospective analyses performed. As such, 20 body-points were included in the LUPE pose estimation network built in DLC: snout, upper mouth, middle fore-paw digit and palm of left and right forepaws, all digits and palm of left and right hindpaws, genital region, and tail base.

Model was trained iteratively 17 times for up to 350,000 iterations per training when loss and learning rate plateaued. Network architecture and augmentation method chosen was resnet_50 and imgaug, respectively. Percent of dataset trained on was 95% with the remaining 5% remaining used as test dataset for evaluation. For the final training iteration, the mean average Euclidean error between the manual labels and the ones predicted by model was 2.2 pixels (0.073 cm) for training dataset error and 2.33 pixels (0.077 cm) for test dataset error.

Frames for labeling were extracted manually focused on specific behavior sequences and individual frames not accurately or confidently labeled by the model. After each training, frames for data input were added as needed for accuracy and confidence to label videos trained on and novel videos analyzed through LUPE not trained in the model. The total number of frames labeled was 14554, with 10,825 (74.38%) and 3,729 (25.62%) frames coming from male and female video data files, respectively. There were 169 total number of unique, mouse video files that frames were extracted from (133, 78.69% male; 36, 21.31% female). The behavioral assays chosen for recording and model input capture different experimental paradigms and chemically evoked manipulations. From this, the model is able to assign pose data points with high accuracy and confidence for both male and female mouse video data from a variety of behavioral data.

The male video data included: subcutaneous saline injection response (210, 1.94%), subcutaneous morphine 10 mg/kg response (1105, 10.21%), left hindpaw intraplantar capsaicin response (666, 6.15%), left hindpaw intraplantar 1% formalin (960, 8.87%), left hindpaw intraplantar 5% formalin (1271, 11.74%), habituation to LUPE chamber (720, 6.65%), formalin left hindpaw intraplantar injection (829, 7.66%), formalin right hindpaw intraplantar injection (719, 6.64%), formalin cheek injection (472, 4.36%), SNI left hindpaw injury day-0 (170, 1.57%), SNI left hindpaw injury day-3 (508, 4.69%), SNI left hindpaw injury day-7 (380, 3.51%), SNI left hindpaw injury day-21 (255, 2.36%), SNI right hindpaw injury day-0 (652, 6.02%), SNI right hindpaw injury day-3 (508, 4.69%), SNI right hindpaw injury day-7 (510, 4.71%), SNI right hindpaw injury day-21 (254, 2.35%), and naloxone precipitated morphine withdrawal (636, 5.88%).

The female video data included: subcutaneous morphine 10 mg/kg response (133, 3.57%), habituation to LUPE chamber (320, 8.59%), formalin left hindpaw intraplantar injection (148, 3.97), formalin right hindpaw intraplantar injection (150, 4.02%), SNI left hindpaw injury day-0 (1050, 28.17%), SNI left hindpaw injury day-3 (450, 12.07 %), SNI left hindpaw injury day-7 (350, 9.39%), SNI right hindpaw injury day-0 (826, 22.16%), SNI right hindpaw injury day-3 (150, 4.02%), and SNI right hindpaw injury day-7 (150, 4.02%).

##### Behavior Classification via A-SOiD/B-SOiD

We trained a Random Forest Classifier to predict 5 different behaviors (still, walking, rearing, grooming, licking hindpaw) given the pose estimation of the previously described 20 body parts. This supervised classifier is refined using an active learning approach over 27 iterations, with a total of 51,377 frames (still: 11,599, walking: 12,809, rearing: 7,270, grooming: 12,719, and licking hindpaw: 6,971). Upon reaching an average f1 score amongst the 5 classes was 93.5%, we predicted all existing pose files and segmented licking hindpaw into licking left hindpaw and licking right hindpaw as they were clearly dissociable. After splitting the laterality of licking hindpaw, we retrained the Random Forest Classifier to expand its classification from 5 classes to 6 classes. The final average f1 score amongst 6 classes was 94.3%.

##### LUPE Analyses

Once the model is trained, we predicted all the behavioral data in this paper with the same Random Forest Classifier model. Due to the nature of intermittent pose estimation noise, we decided to smooth the output behavior to only consider continuous bouts that are >= 200ms.

To analyze behavior ratio over time (seen in Figure 1), we calculated the counts for each behavior for each minute then normalized by the total number of frames. This quantification allows us to track when a particular behavior occurs during each session. To explore the variability across animals, we plotted the mean +-standard error of the mean.

To analyze the distance traveled, one body-point that is high in confidence for pose detection and is always present in behavioral sequences was chosen, being the tail base, to calculate the Euclidean distance between consecutive frames of the tail base position. This was calculated by subtracting the x- and y-coordinates of the tail base between consecutive frames and then calculating the Euclidean norm of the resulting vector. Distance calculated in pixels was then converted to a specified, being centimeters, using a conversion factor of 0.0330828 cm/pixels. This conversion factor is unique to the aspect ratio of our frames and resolution of the video data. To explore the variability across animals, we plotted the mean +-standard error of the mean.

Heatmaps of the distance traveled were generated by constructing a 2D histogram of the tail base x- and y-coordinates. The code functions by binning the pose data points into a specified number of bins (being 50 in this case) along each x- and y-coordinate range. The ‘counts’ collected represent the frequency of occurrences in the 2D histogram that fall in each bin representing a range of x- and y-coordinates.

#### Identification of behavior states and the AMPS, Affective-Motivational Pain Scale

##### Behavioral state identification

60Hz LUPE behavior scores from all male and female animals in all pain models used in the paper were downsampled to 20Hz by taking the mode of every 3 frames. Transition matrices were generated between each behavior in which values were expressed as % of all frames at behavior bx at time t would go on to behavior by at time t+1. Transition matrices were taken over 30 second windows sliding by 10 second increments within animals, so as not to miss transitions. Window size was chosen in line with empirical findings that showed spontaneous bouts of intense subjective pain under chronic pain conditions last 22.5 ± 22.1s (mean, S.D.) (Baliki et al., 2006). Transition matrices were transformed into single rows such that each transition matrix became a single vector of probabilities 36 possible transitions: p(still_t+1_ | still_t_), p(walk_t+1_ | still_t_), p(rear_t+1_ | still_t_)…p(right lick_t+1_ | right lick_t_). These probability vectors were then stacked to create a matrix of 215,760 observations (3,596 observations for 60 animals) by 36 transitions. These observations were clustered using 100-fold cross-validated K-Means, where the silhouette and elbow methods robustly converged at 6 clusters over 100 iterations. Each of 6 centroids thus defined a single behavioral state that could be expressed as reconstructed transition matrices.

##### Behavioral state classification

To classify each time-point as one of six behavioral states, the same process as described above was repeated to generate smoothed transition matrices over time for each animal. At each timepoint the Euclidean distance was calculated between its given transition matrix and each model centroid. The state at that timepoint was chosen to minimize the distance from the true transition matrix and the model centroid. Model fit for each animal in each session is thus expressed as the mean distance from the nearest centroid to its real transition matrix over the session. As distance approaches 0, the model approaches perfect fit. Transition matrices randomly shuffled over probabilities show that, at chance, model fit converges between 2-3 A.U.s, whereas true model fit ranges between 0-1.5 A.U.s, indicating genuine discovery of behavioral structure at a timescale of seconds-to-minutes.

##### Behavior state model validation

To ensure that states were not trivially dependent on the occurrence or absence of a single behavior, states were classified after systematically removing each behavior from the dataset. Model fit and the % of observations matching the original classifications were compared to those of the shuffled dataset.

##### Pain scale

To distill behavioral states into a single index of pain, PCA was performed on the state distributions of each animal in the uninjured, capsaicin, and formalin experiments that received 0mg/kg morphine. Each animal was described by the fraction time spent in each state, yielding a six-dimensional dataset. The scores of each animal along the first two PCs were considered. To yield scores for animals in every other experiment and condition, state distributions were projected into this PC state by subtracting the mean of the original dataset and matrix multiplying by the coefficients defining the PC space. We pre-determined that the PC that scales oppositely with pain condition and analgesia would be designated the AMPS “pain scale.” PC2 met this requirement.

##### Within-state behavior dynamics

To assess temporal dynamics of behaviors over states, the binarized behavior classification vector (behavior of interest = 1, all others = 0) over the course of a given bout of a state were resampled to be 100 steps long. These vectors were pooled across bouts and animals in a given condition and averaged to yield behavior probability as a fraction of time completed in state. The fraction of time remaining in state with respect to a behavior occurrence was calculated by subtracting the absolute time of the behavior from the absolute end time of that state bout and dividing by the duration.

For simulated behavior dynamics, 100 Markov simulations of behaviors based on the State 4 centroid transition matrix were produced over the course of a state for each possible initial condition (Stillness, Walking, Rearing, Grooming, Licking) and allowed to proceed for the empirically determined average number of time steps State 4 lasts (600) before undergoing the same procedure.

#### *In vivo* calcium imaging

##### Miniscope surgery

For Miniscope studies, all mice underwent an initial intracranial injection using previously described methods, followed two weeks later by a GRIN lens implant surgery. During the intracranial injection surgery, 800nL of AAV9-hSyn-jGCaMP8m at a titer of 1.2e12 (Addgene virus #162375) was infused into the right ACC (AP +1.5, ML 0.3, DV -1.5 mm). Two weeks later, GRIN lens implantation surgeries occurred and followed the same protocol until the craniotomy step. A 1mm craniotomy was made by slowly widening the craniotomy with the drill. The dura was peeled back using microscissors, sharp forceps, and curved forceps. The craniotomy was regularly flushed with saline, and gel foam was applied to absorb blood. An Inscopix Pro-View Integrated GRIN lens and baseplate system was attached to the Miniscope and stereotax. Using the Inscopix stereotax attachment, the lens was slowly lowered into a position over the injection site. The final DV coordinate was determined by assessing the view through the Miniscope stream. If tissue architecture could be observed in full focus with light fluctuations suggesting the presence of GCaMP-expressing cells, the lens was implanted at that coordinate (−0.6 to -0.3mm DV). The GRIN lens/baseplate system was secured to the skull with Metabond, followed by dental cement. After surgery, mice were singly housed and injected with Meloxicam for three consecutive days during recovery.

##### Miniscope data collection – Acute capsaicin

Miniscope neural activity and associated behavior data was collected over two days - baseline/capsaicin (test day 1) and morphine/capsaicin (test day 2) - with two weeks in between test days. On each day, the Inscopix nVista3.0 Miniscope was first affixed on the mouse and the ideal focus was determined based on the field of view. Imaging parameters (power 0.7 mW/mm2, gain 2) were held consistent across all mice and test days. Mice were injected with saline (test day 1) or 0.5 mg/kg morphine (test day 2) and placed in LUPE. Five minutes later, the miniscope and LUPE recordings were started and continued for 20 minutes uninterrupted. The recording was then stopped for 5 minutes to reduce photobleaching risk. Next, mice were injected with 2ug capsaicin (HelloBio HB1179) in the left hindpaw (both test days) using a Hamilton syringe affixed with a 30G needle and placed in LUPE. Both miniscope and LUPE recording was restarted immediately after and continued for 30 minutes.

##### Miniscope data collection – Chronic neuropathic pain

Behavior and neural activity of mice were recorded 8 times before, during, and after the onset of SNI. Mice were tested at baseline (1 day before SNI), and then 1 hour, 1 day, 3 days, 1 week, 2 weeks, and 3 weeks post-SNI. One day after the 3 week testing session mice underwent another test day where they were injected with 0.5mg/kg morphine 30-min prior to recording. Each testing day consisted of a 30-minute LUPE recording. Ideal imaging parameters were determined on each day and neural activity was aligned to LUPE behavior tracking via a TTL pulse at the start of the recording session.

##### Miniscope data collection – Pain and valence panels

Within 1 week after completion of the chronic neuropathic pain LUPE testing, mice underwent exposure to a panel of acute stimuli while neural activity was recorded. Animals were placed in transparent containers placed on top of a metal hexagonal-mesh floored platform. Stimuli were applied to the underside of the left plantar hind paw. Animals were continuously recorded by two web cameras.

The stimuli were light touch (0.16g Von Frey filament), 30°C water, pin prick (25G syringe needle), acetone, and hot water (55°C). Water and acetone stimuli were delivered using a needle-less syringe and a droplet of the liquid was applied. Five presentations of each stimulus were administered to the left hindpaw with 90 seconds in between each presentation.

After the pain panel, mice underwent 3 days of 20-minute training for the valence panel, where they learned to lick a 10% sucrose (Sigma-Aldrich S7903, diluted in water) droplet from a needleless syringe poked through the mesh of the metal rack. On the day of the panel, neural activity was recorded while they licked sucrose (approx. 7 presentations), with at least 45 seconds in between each presentation, then the liquid in the syringe was switched to 0.06mM quinine (Sigma-Aldrich Q1125, diluted in water). Quinine was presented until mice licked at least 5 times. Finally, 7 presentations of 55°C water were applied to the left hindpaw. Both sucrose and quinine concentrations were adapted from Corder et. al., 2019.

##### Calcium imaging pre-processing

Videos were downloaded from the Inscopix Data Acquisition Box and uploaded to the Inscopix Data Processing Software (IDPS). Videos were spatially downsampled by a factor of 4 and spatial bandpass filtered between 0.005 and 0.500. Videos were then motion corrected with respect to their mean frame. Cells were identified and extracted using CNFM-E (default parameters in the Inscopix implementation of CNMF-E, except the minimum in-line pixel correlation = 0.7 and minimum signal to noise ratio = 7.0) and second-order deconvolved using SCS.

##### Spontaneous activity in neurons

Deconvolved activity in each neuron was z-scored. Peaks were identified using the findpeaks function in matlab with the argument “MinPeakProminence” set to 1.

##### Neural encoding of behavior probabilities

Categorical frame-by-frame vectors of behavior values (still, walking, rearing, grooming, licking left hindpaw, and licking right hindpaw) were downsampled from 60 to 20Hz to match sampling rates of neural recordings by taking the floor of the mode for within a sliding window of 3 frames. The probability of engaging in a behavior as generated by this K-Means-GLM model thus takes into account recent behavior history and slowly evolving state of the animal. These higher-order measurements of behavioral state have been linked to neural activity previously (Bolkan et al., 2022, Piet et al., 2024, Tervo et al. 2021). To identify neurons encoding this higher-order index of behavioral state, a binomial GLM was trained to predict binarized probabilities of a given behavior (thresholded at p(behavior)=0.5) from principal components of ACC activity explaining 80% of the variance in individual animals. Neurons with a coefficient > 1.5 z-score in the three most highly weighted PCs were classified as p(behavior)-encoding neurons. Neurons that on average increased or decreased their activity in the 500ms following onset of behavior were classified as behavior+ or behavior-neurons, respectively.

##### Behavior-evoked activity

To assess the modulation of activity in these neurons by behavior, peri-behavioral time histograms of neural activity were generated from two seconds before to two seconds after the start of each behavior bout. To quantify, peri-behavioral time histograms were z-scored to the one second before behavior onset and areas-under-the-curve were taken for each of the two seconds after the bout start. If a neuron was suppressed after behavior onset, it was designated a behavior-off neuron and vice versa for a behavior-on neuron. Behavioral tuning curves were generated by taking the z-score of the activity of each population of encoding neurons during each scored behavior. Selectivity of these neurons for a given behavior was calculated by taking the d’ between their encoded behavior and each other behavior:

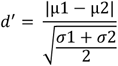

##### Behavior, sensory, and state decoding

For behavior and sensory decoding, a Fisher linear decoder was trained to predict each behavior or sensory stimulus in a given session from the activity of all neurons. For state decoding, an SVM decoder was trained to predict whether animals were in a Pain or Non-Pain State using the activity of lick probability-encoding neurons. Each decoder for each animal and session underwent 100-fold cross-validation, training on a random 80% of the data each time and testing on 20%. Data was randomly subsampled such that there were the same number of samples for each class to eliminate training bias. Decoders were also trained on randomly shuffled data as a control. Confusion matrices were generated, averaged over cross-validations and normalized by the true frequency of each behavior in the test set. Significance was determined using the permutation test.

##### Identifying stimulus-active neurons

For pain and valence panels, activity of each neuron from 0-3s after stimulus was compared to activity -3 - -1s before stimulus with a permutation test (FDR threshold p<0.01).

#### Single nuclei RNA sequencing

##### Nuclei preparation

A single punch of the right side of the BLA measuring 2 mm in width and 1 mm in depth, was used to prepare the nuclei suspensions. Nuclei isolation was performed utilizing the Minute™ single nucleus isolation kit designed for neuronal tissue/cells (Cat# BN-020, Invent Biotechnologies). Briefly, the tissue was homogenized using a pestle in a 1.5 mL LoBind Eppendorf tube. Subsequently, the cells were resuspended in 700 µl of cold lysis buffer and RNAse inhibitor and incubated on ice for 5 minutes. The homogenate was then transferred to a filter within a collection tube and incubated at -20°C for 8 minutes. Following this, the tubes were centrifuged at 13,000 x g for 30 seconds, the filter was discarded, and the samples were centrifuged at 600 x g for 5 minutes. The resulting pellet underwent one wash with 200 µL of PBS + 5% BSA and then resuspended in 60 µL of PBS + 1% BSA. The concentration of nuclei in the final suspension was assessed by staining with Trypan Blue and counted using a hemacytometer. The suspension was diluted to an optimal concentration of 500-1000 nuclei/µL.

##### Single-Nuclei Gene Expression Assay

Nuclei suspensions were used as input for the 10x Genomics 3’ gene expression assay (v3.1), following manufacturer protocols. A total of 20,000 nuclei were loaded into the 10x Genomics microfluidics Chromium controller, with the aim of recovering approximately 10,000-12,000 nuclei per sample. Subsequently, sequencing libraries were constructed, and unique dual indexed libraries were pooled together at equimolar concentrations of 1.75 nM and sequenced on the Illumina NovaSeq 6000, using 28 cycles for Read 1, 10 cycles for the i7 index, 10 cycles for the i5 index, and 90 cycles for Read 2.

##### Data analysis

###### Preprocessing of snRNAseq data

Paired end sequencing reads were processed using 10x Genomics Cellranger v5.0.1. Reads were aligned to the mm10 genome optimized for single cell sequencing through a hybrid intronic read recovery approach ^121^. In short, reads with valid barcodes were trimmed by TSO sequence, and aligned using STAR v2.7.1 with MAPQ adjustment. Intronic reads were removed and high-confidence mapped reads were filtered for multimapping and UMI correction. Empty GEMs were also removed as part of the pipeline. DESeq2 was used to compare expression at the 3-day, 3-week, and 3-month timepoints to control animals for each cluster. Pseudobulked expression differences were assessed by Wald test and a false discovery rate (FDR) of 0.05 was used to correct for multiple testing.

###### Clustering and comparison

Count matrices for each individual sample were converted to Seurat objects using Seurat 4.3, and nuclei were filtered with thresholds of > 200 minimum features and < 5% mitochondrial reads. Initial dimensionality reduction and clustering was performed to enable removal of cell free mRNA using SoupX ^122^. SCTransform was used to normalize and scale expression data and all samples were combined using the Seurat integration method. Putative doublets identified by DoubletFinder, as well as residual clusters with mixed cell type markers or high mean UMI, were removed. The cleaned dataset was clustered using the first 20 PCs at a resolution of 0.3. Cluster identity was determined by expression of known marker genes.

###### Modular activity scoring

Modular activity scores were calculated for all clusters using AddModuleScore with the list of the 25 putative immediate early genes (*Arc, Bdnf, Cdkn1a, Dnajb5, Egr1, Egr2, Egr4, Fos, Fosb, Fosl2, Homer1, Junb, Nefm, Npas4, Nr4a1, Nr4a2, Nr4a3, Nrn1, Ntrk2, Rheb, Sgsm1, Syt4, Vgf*) against a control feature score of 5 ^123^.

###### Gene ontology analysis

Gene ontology (GO) analysis was performed on differentially expressed genes (DEGs) identified between uninjured and spared nerve injury (SNI) conditions. To investigate the functional enrichment of genes upregulated in the SNI condition, we used SynGO (https://www.syngoportal.org/), a synapse-specific, evidence-based GO annotation platform that provides curated information about synaptic genes and their roles in biological processes, molecular function, and cellular localization. Gene symbols corresponding to upregulated DEGs were submitted to the SynGO web-based analysis tool. Enrichment analysis was performed against the background of all protein-coding human genes mapped to orthologous mouse genes, using SynGO’s default statistical settings. We focused on enrichment within high-confidence, expert-curated GO categories based on experimental evidence. Overrepresentation testing was conducted with false discovery rate (FDR) correction, and enriched GO terms were considered significant at FDR < 0.05. The analysis enabled functional interpretation of synaptic gene expression changes in chronic pain conditions, with particular attention to processes related to synaptic signaling, neurotransmitter transport, and pre- or post-synaptic structural components.

#### Quantitative PCR

Cellular RNA was extracted with RNAzol (Sigma-Aldrich, R4533) according to manufacturer’s protocol, and cDNA was synthesized from 1 ug RNA (Applied Biosystems, #4374966). cDNA was diluted 1:10 and assessed for mRNA transcript levels by qPCR with SYBR Green Mix (Applied Biosystems, #A25741) on a QuantStudio7 Flex Real-Time PCR System (Thermo Fisher). Oligonucleotide primer sequences for target and reference genes are as follows: mouse_GAPDH (forward: AACGACCCCTTCATT-GACCT, reverse: TGGAAGATGGTGATGGGCTT), mouse_L30 (forward: ATGGTGGCCGCAAAGAAGACGAA, reverse: CCTCAAAGCTGGACAGTTGTTGGCA), mouse_*OPRM1* (forward: CTGCAAGAGTTGCATGGACAG, reverse: TCAGATGACATTCACCTGCCAA). The fold change in the target mRNA abundance was normalized by the reference gene GAPDH, was calculated using the 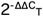 method^124^.

#### Closed-loop, Optogenetic Real-Time Place Preference (oRTPP) & Conditioned Place preference (oCPP)

First, *pain*TRAP2 mice injected with AAV1-*MORp*-DIO-iC++ and bilateral fiber-optic implants (200-µm diameter, 0.66-NA, 700-µm pitch between each fiber center; Thorlabs), with SNI or no-injury, were habituated for 30-min a day for 5-days in a holding cage. For basal place-preference measurements (*Pre-test session*), mice with attached patch cables, were placed in a two-chamber acrylic box (60 × 25 × 30 cm3), with each side of the chamber measuring 30 × 25 cm2), at room temperature (∼23 °C), under ∼10 lux red-light. Each chamber had different contextual pattern cues on the wall: one side had a black-and-white-striped pattern and the other a dotted pattern. Mouse movements were recorded in real-time with an overhead top-view Basler camera connected to Ethovision tracking software (Noldus), connected via a mini-I/O box to a 450-nm LED and pulse-wave generator (Prizmatix), for 30-min. Chamber preference times were quantified by Ethovision for the amount of time spent in each chamber. Using a biased-design, based on the quantified basal preference, mice received

LED stimulation in the non-preferred chamber, as our hypothesis was that activation of iC++ would reduce spontaneous pain and drive increased dwell-time in the LED-paired chamber. The following day, to assess oRTPP, mice were placed in the center of the apparatus for a 30-min session. During this session, Ethovision body-contour tracking of the mouse center-point activated the blue LED when the center-point was detected in the originally non-preferred chamber, which in turn delivered 10-mW continuous light through the bifurcating fiber optic implants for the entire duration that the mouse center-point was detected in the chamber; the LED would turn off when the mouse center-point was detected in the other chamber (i.e. Closed-loop protocol). This procedure was repeated daily for seven consecutive sessions to oRTPP-induced learning rates over time. After the Pre-test, and seven closed-loop optogenetic sessions, we performed one Post-test session to assess if a conditioned place preference had developed. Here, mice remained connected to the patch cables but no light was delivered, and the time spent in each chamber was calculated. Increased time in the formally LED-paired chamber indicates a learned preference for iC++-mediated inhibition of ACC nociceptive MOR+ cell-types.

## Supplemental Figures

**Figure S1.**
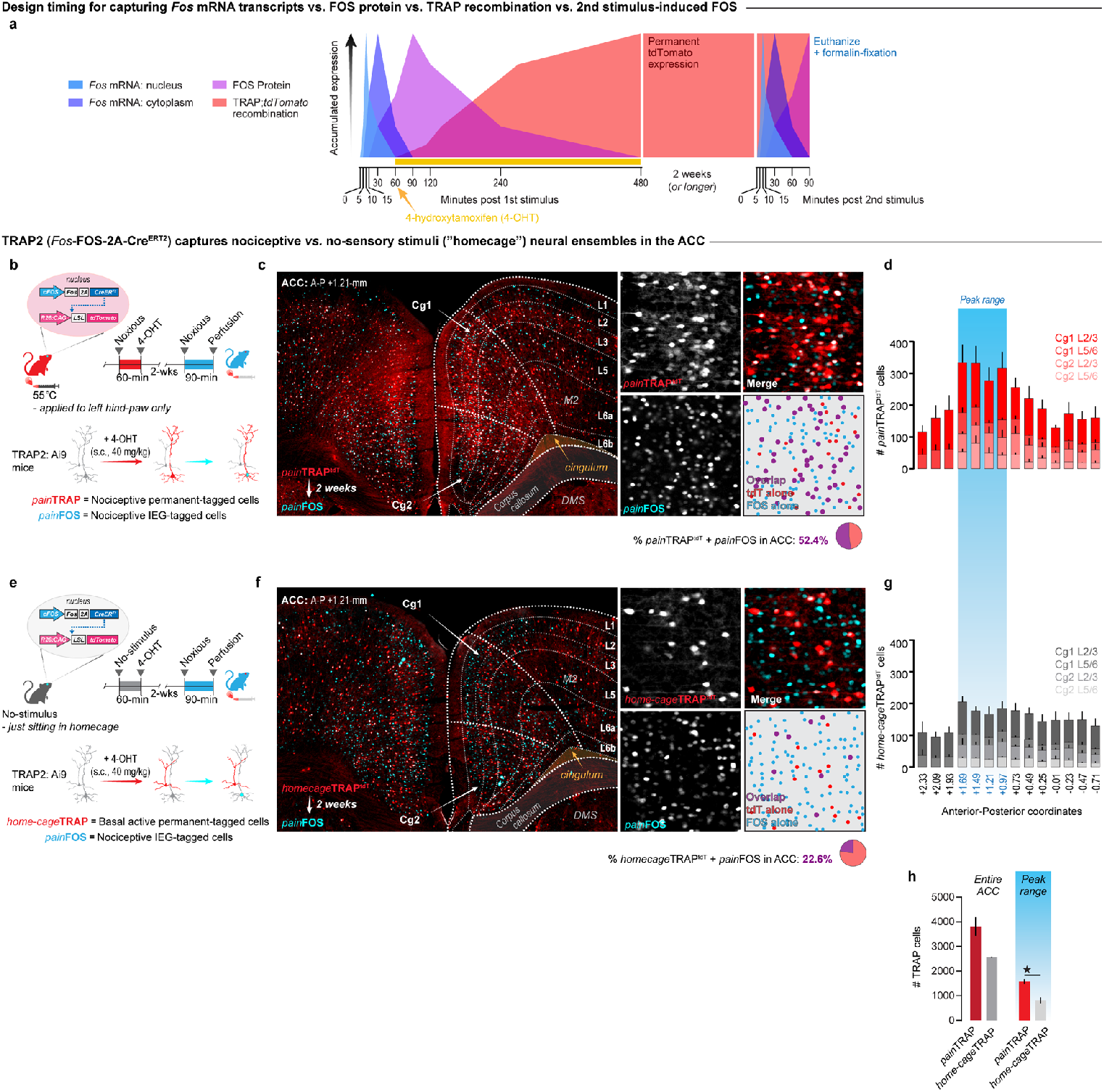
Genetic capture and tagging of noxious stimulus-responsive neurons in the ACC. **(a**) Timing strategy of 4-hydroxytamoxifen-mediated recombination in TRAP2 mice for activity-dependent genetic capture and permanent tagging of stimulus-active neurons, paired with stimulus-driven FOS expression and timed tissue collection for visualization of multiple stimulus-activity pairings. **(b)** Utilizing TRAP2 mice for permanent fluorescent tagging of nociceptive cell-types (*pain*TRAP), and noxious re-engagement visualized with the immediate early gene, FOS (*pain*FOS). **(c)** ACC *pain*TRAP tagging and *pain*FOS co-expression. **(d)** Quantification of *pain*TRAP cells across all layers of the dorsal Cg1 and ventral Cg2, displayed across the entire anterior-posterior extent of the ACC. **(e)** Utilizing TRAP2 mice to permanently tag active neurons in a no-stimulus control condition (*home-cage*TRAP) paired with timed tissue collection of neurons active to a noxious stimulus via expression of FOS (*pain*FOS). **(f)** Expression and overlap of *homecage*TRAP tagged cells and *pain*FOS expression. **(g)** Quantification of *home-cage*TRAP cells across layers and anterior-posterior coordinates of ACC. (Two-way ANOVA) **(h)** Comparison of TRAP cells in *pain*TRAP and *home-cage*TRAP conditions across the entire ACC and within the peak range that displayed the highest values (Unpaired t-test, p=0.0402). For detailed statistics, see **Supplemental Table 5**. ⋆ = P < 0.05. Errors bars = s.e.m.

**Figure S2.**
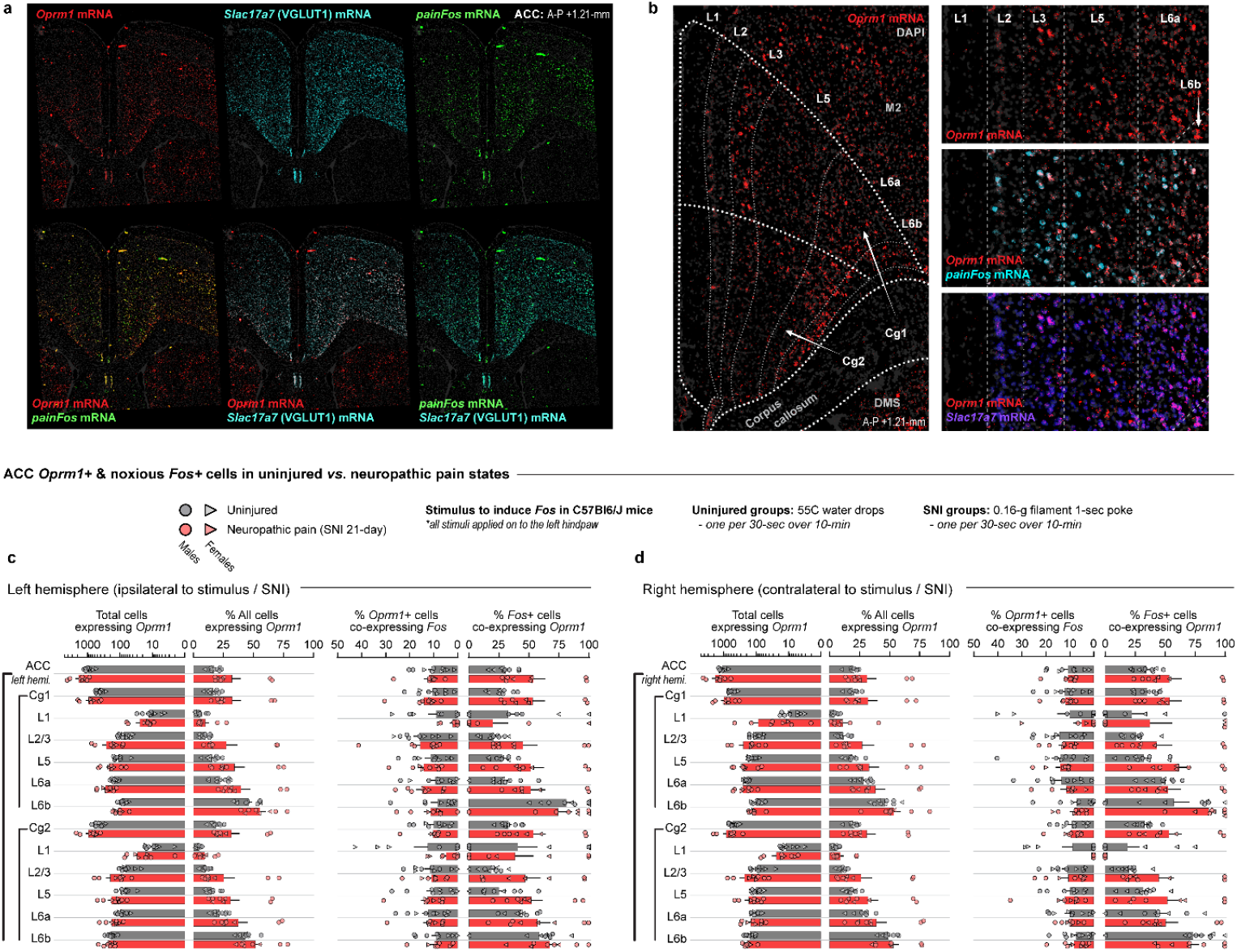
FISH quantification of nociceptive and opioidergic ACC neurons in chronic neuropathic pain. **(a)** Representative 20X stitched images for *Oprm1*, noxious-evoked *Fos* (pain*Fos*) and *Slc17a7* mRNA using Fluorescent *in situ* hybridization (FISH). **(b)** Expression of *Oprm1*, and its overlap with pain*Fos* and *Slc17a7* mRNA across the layers of ACC in uninjured male and female mice. **(c)** *Oprm1* mRNA quantification across the left hemisphere (ipsilateral to injury) of ACC, broken down by subregion and layers, in male and female mice with or without neuropathic pain. **(d)** Same as (C) but for the right hemisphere, contralateral to the injury. (N = 8 neuropathic (5 male, 3 female) and 9 uninured (4 male, 5 female)).

**Figure S3.**
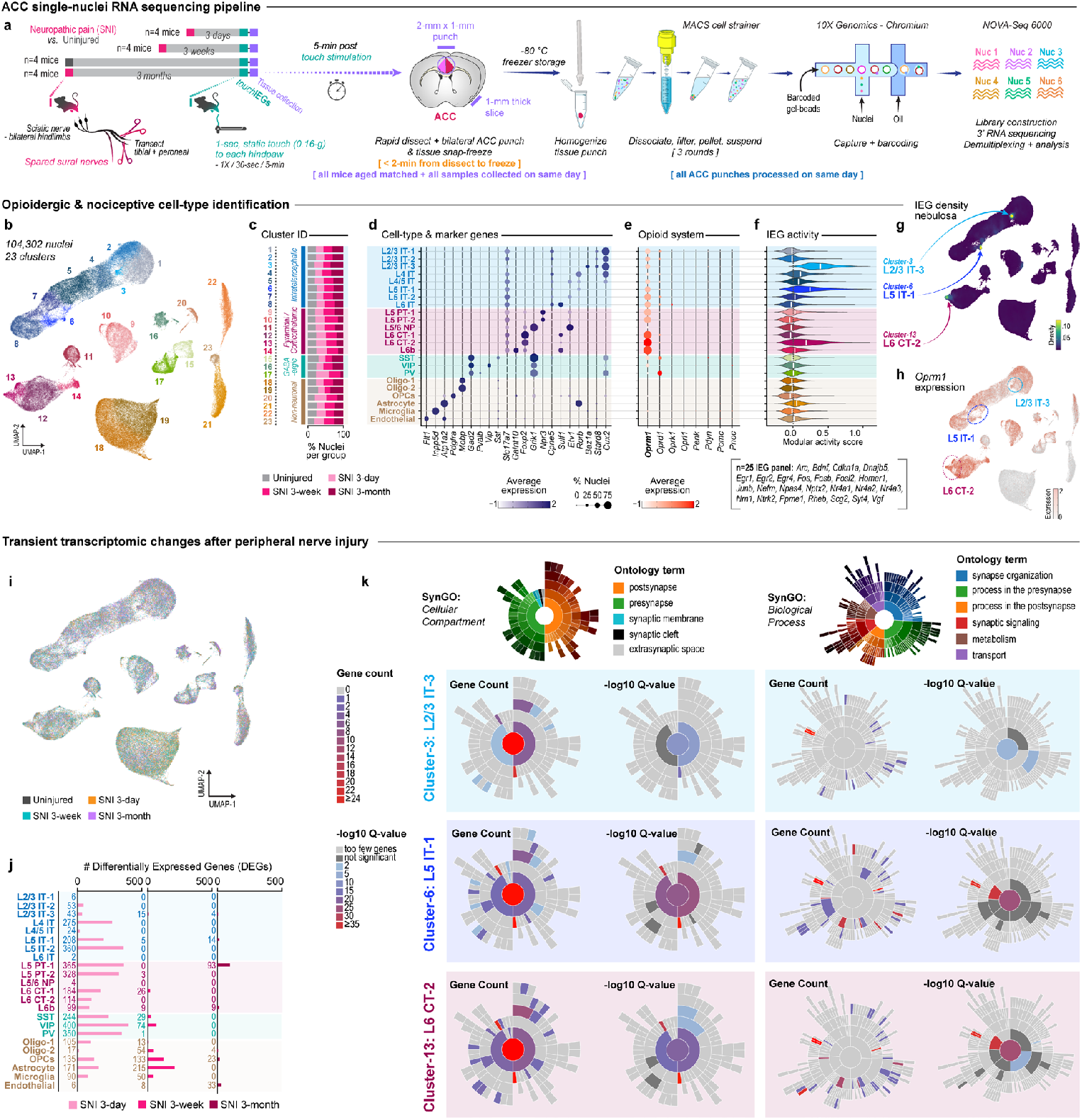
Transcriptomic classification of nociceptive and opioidergic cell-types in the cingulate. **(a)** Design of single-nuclei RNA sequencing experiments from ACC punches of mice with bilateral sciatic nerve transections collected after different timepoints of chronic pain development and light-touch-evoked immediate early genes (IEGs); n=4 male mice per condition. **(b)** Dimensionality reduction plot of all nuclei from all groups in 23 cell-type clusters. **(c)** Broad classification of all clusters into neuronal and non-neuronal classes with the percentage of nuclei per condition in each cluster. **(d)** Specific cell-type identities and dot-plot for selected marker genes. **(e)** Dot plot for all opioid receptors and peptides. **(f)** Nociceptive activity score per cluster based on a panel of n=25 IEGs. **(g)** Nebulosa density plot of the IEG panel overlaying the UMAP of cell-type clusters. **(h)** Feature plot for *Oprm1* expression. **(i)** Intermixed nuclei with UMAP space from Uninjured mice and the three post-Spared Nerve Injury (SNI) timepoints: 3-days, 3-weeks and 3-months. **(j)** The number of DEGs per cell-type for each injury condition compared to the Uninjured condition. **(k)** Synaptic Gene Ontology (SynGO) analysis for differential expressed genes (DEGs) in ACC nociceptive cell-types (L2/3 IT-3 (light blue), L5 IT-1 (dark blue), L6 CT-2 (maroon) 3-days after SNI compared to Uninjured. *Left*: DEG gene count and logQ10 enrichment related to Cellular Components, such as presynapse. *Right*: DEG gene counts and logQ10 enrichment related to Biological Processes, such as vesical recycling.

**Figure S4.**
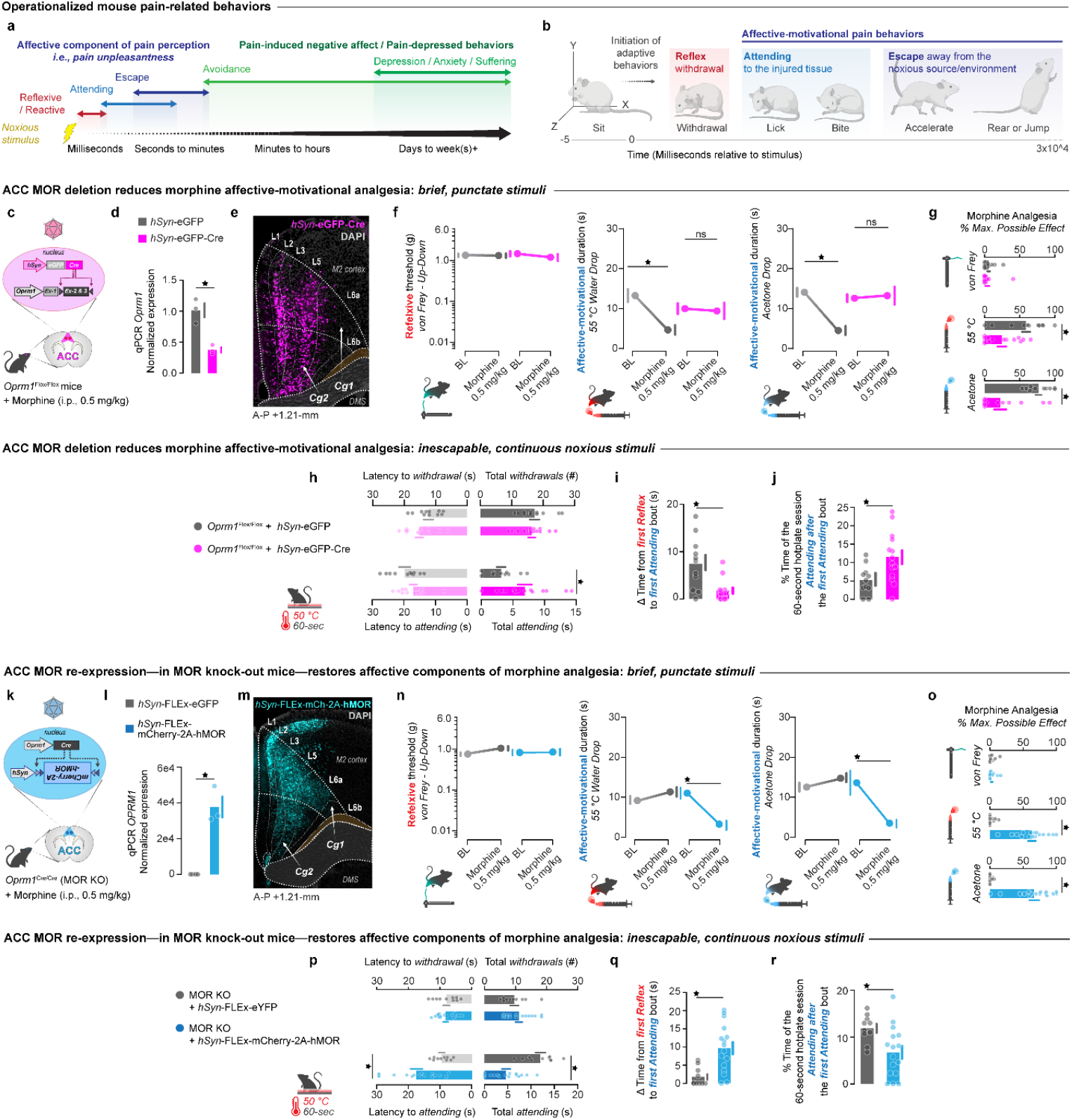
Morphine analgesia is mediated by action on ACC MOR+ cell types. **(a)** Schematic of the timing of common behavioral responses to noxious stimuli over seconds to weeks. **(b)** Classification and timing of reflexive and affective-motivational pain behaviors. **(c)** Design to virally-delete MORs from ACC neurons in *Oprm1*^Flox/Fox^ mice (N=11 *hSyn*-eGFP mice, N=14 *hSyn*-eGFP-Cre mice; 7 males and 7 females). **(d)** ACC *Oprm1* qPCR (N=3 males per group, two-tailed Student’s t-test, p=0.002). **(e)** Expression of *hsyn*-eGFP-Cre injected into ACC of *Oprm1*^Flox/Fox^ mice. **(f)** Reflexive withdrawal threshold and time spent engaging in affective-motivational pain behaviors before and after administration of morphine (0.5 mg/kg, i.p.) in mice with virally deleted ACC MORs (*hsyn*-eGFP-cre) or controls (*hsyn*-eGFP). 2-Way ANOVA + Tukey, Von Frey, interaction p=0.2; 55°C, interaction p=0.007; Acetone, interaction p=0.003. **(g)** Morphine analgesia for mechanical and thermal noxious stimuli. t-Test, 2-tailed, unpaired: von Frey p=0.66; 55° p=0.015; acetone p=0.001. **(h)** Behavior metrics for thermal thresholds and affective-motivational engagement on an inescapable hotplate after morphine. t-Test, 2-tailed, unpaired: Latency withdrawal, p=0.089; total withdrawal, p=0.617; latency to attend, p=0.131; total attending, p=0.0123. (**i)** Time between the first reflexive withdrawal and first attending bout on the inescapable hotplate after morphine. (t-Test, 2-tailed, unpaired p=0.0015) (**j)** Proportion of time spent engaging in attending behaviors after the first bout of such attending behaviors on the inescapable hotplate after morphine. (t-Test, 2-tailed, unpaired, p=0.012) **(k)** Design to virally re-express MORs in an embryonic MOR knockout mouse (N=10 null expression mice, *hsyn*-FLEx-eGFP; N=16 re-expression mice, *hSyn*-FLEx-mCherry-2A-hMOR; 8 males and 8 females). **(l)** ACC *OPRM1* qPCR (N=3 males per group). (t-Test, 2-tailed, unpaired, p=0.0006) **(m)** Expression of *hSyn*-FLEx-mCherry-2A-hMOR injected into ACC of MOR KO mice. **(n)** Reflexive withdrawal threshold and time spent engaging in affective-motivational pain behaviors before and after administration of morphine (0.5 mg/kg, i.p.) in mice with a global knock-out of MORs or with re-expression of ACC MORs. 2-Way ANOVA + Tukey, Von Frey, interaction p=0.66; 55°C, interaction p=0.005; Acetone, interaction p=0.008 **(o)** Morphine analgesia for mechanical and thermal noxious stimuli. t-Test, 2-tailed, unpaired: von Frey p=0.84; 55° p=0.003; acetone p=0.001 **(p)** Behavior metrics for thermal thresholds and affective-motivational engagement on an inescapable hotplate after morphine. t-Test, 2-tailed, unpaired: Latency withdrawal, p=0.12; total withdrawal, p=0.22; latency to attend, p=0.013; total attending, p=0.019. **(q)** Time between the first withdrawal and first attending bout. T-Test, 2-tailed, unpaired p= 0.003. **(r)** Proportion of time of attending behaviors after the first attending bout. T-Test, 2-tailed, unpaired p= 0.021. ⋆ = P < 0.05. Errors bars = s.e.m.

**Figure S5.**
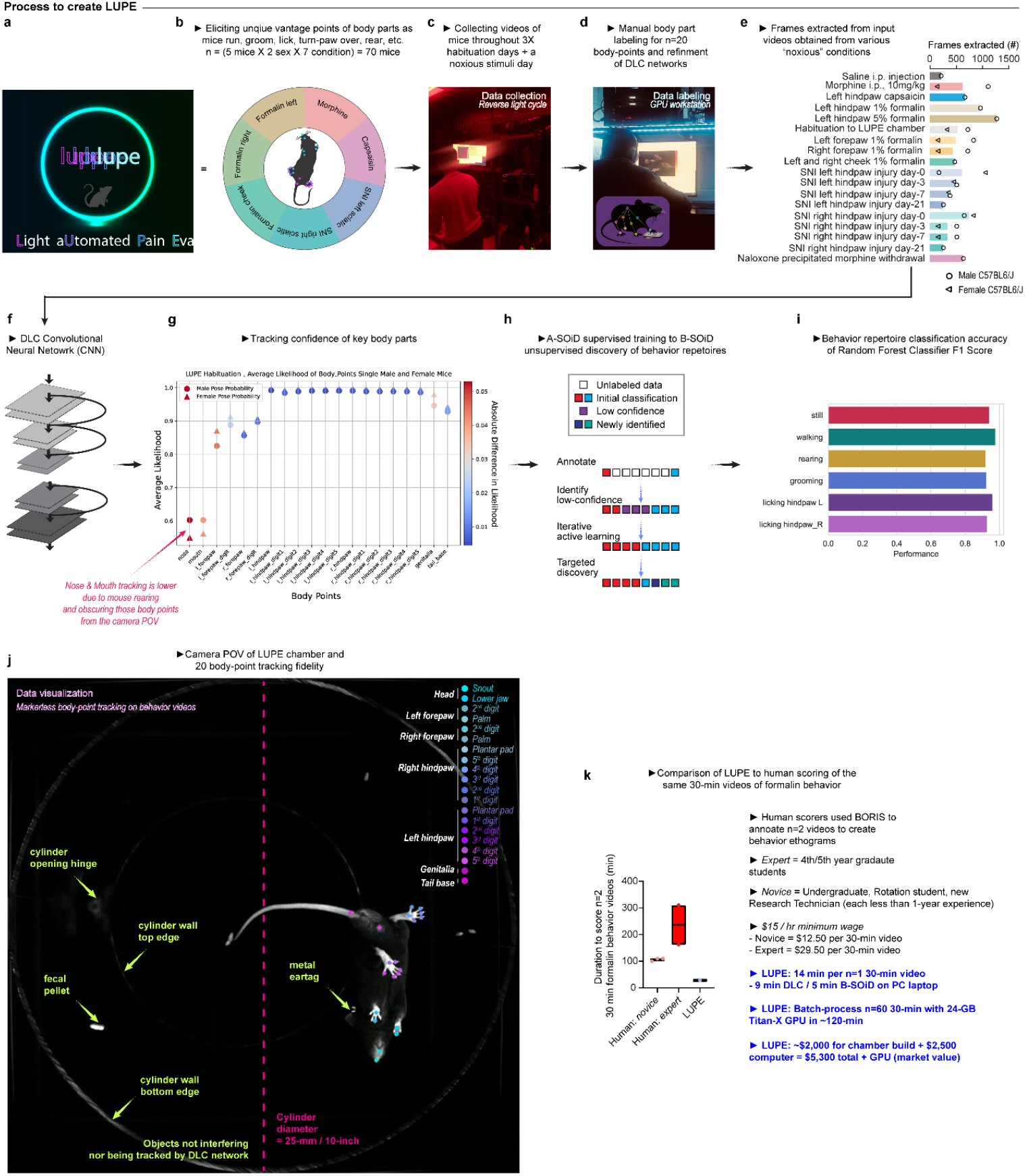
Process to create the LUPE analysis package. **(a)** LUPE (**<u>L</u>**ight a**<u>U</u>**tomated **<u>P</u>**ain **<u>E</u>**valuator), a novel method for studying pain in mice in a controlled environment that allows for streamlined ethological behavior classification using open-source, machine learning algorithms for pose estimation and behavior classification. **(b)** Model training data includes video data from different pain and analgesia behavioral assays, capturing a range of behavior sequences with diverse pose dynamics. **(c)** Experimental assays were collected under reverse light cycles. Video data was collected for three days when mice habituated to behavior room and LUPE chamber and on the fourth day when pain or analgesia assay was performed. **(d)** Input data for the LUPE-DLC network included manually extracted frames that were labeled with 20 body-points for each training iteration using a graphics processing unit (GPU). **(e)** Sum of frames labeled per experimental conditions per sex were manually selected based on sequences of behaviors where pose was not labeled confidently or correctly by the model. **(f)** This procedure of training evaluation, manual frame extraction and labeling, and retraining was iterative, taking advantage of DLC’s convolutional neural network algorithm until pose-estimation performance was qualitatively and **(g)** quantitatively confident and accurate. The quantitative evaluation of the LUPE-DLC model was based on the computed mean average Euclidean error (MAE) between the manual labels and the ones predicted by DLC after each training iteration. The labeled test (DLC produced, unseen images) and training images were evaluated qualitatively to confirm model predictions aligned with required accuracy for pose assignment. **(h)** After developing the LUPE-DLC network, A-SOiD’s active-learning algorithm translated the 2-dimensional pose of the 20 body-points into meaningful behavioral classification. **(i)** The A-SOiD active-learning training procedure was iterated 27 times until the model successfully classified six behavior motifs of interest at high degree of confidence assessed by the random forest classifier F1 score. **(j)** Example image of the bottom-up camera perspective of the LUPE chamber details the maintained pose-estimation of the 20 body-points of interest while ignoring noise of the background in the image. **(k)** To compare the savings of time and money by utilizing LUPE networks, time to complete behavior classification for two 30-minute behavior assays was compared between LUPE, novice (n=3), and expert (n=2) human scorers. LUPE outcompeted human-scorers by requiring approximately 14 minutes to complete behavior classification of one 30-minute behavior recording.

**Figure S6.**
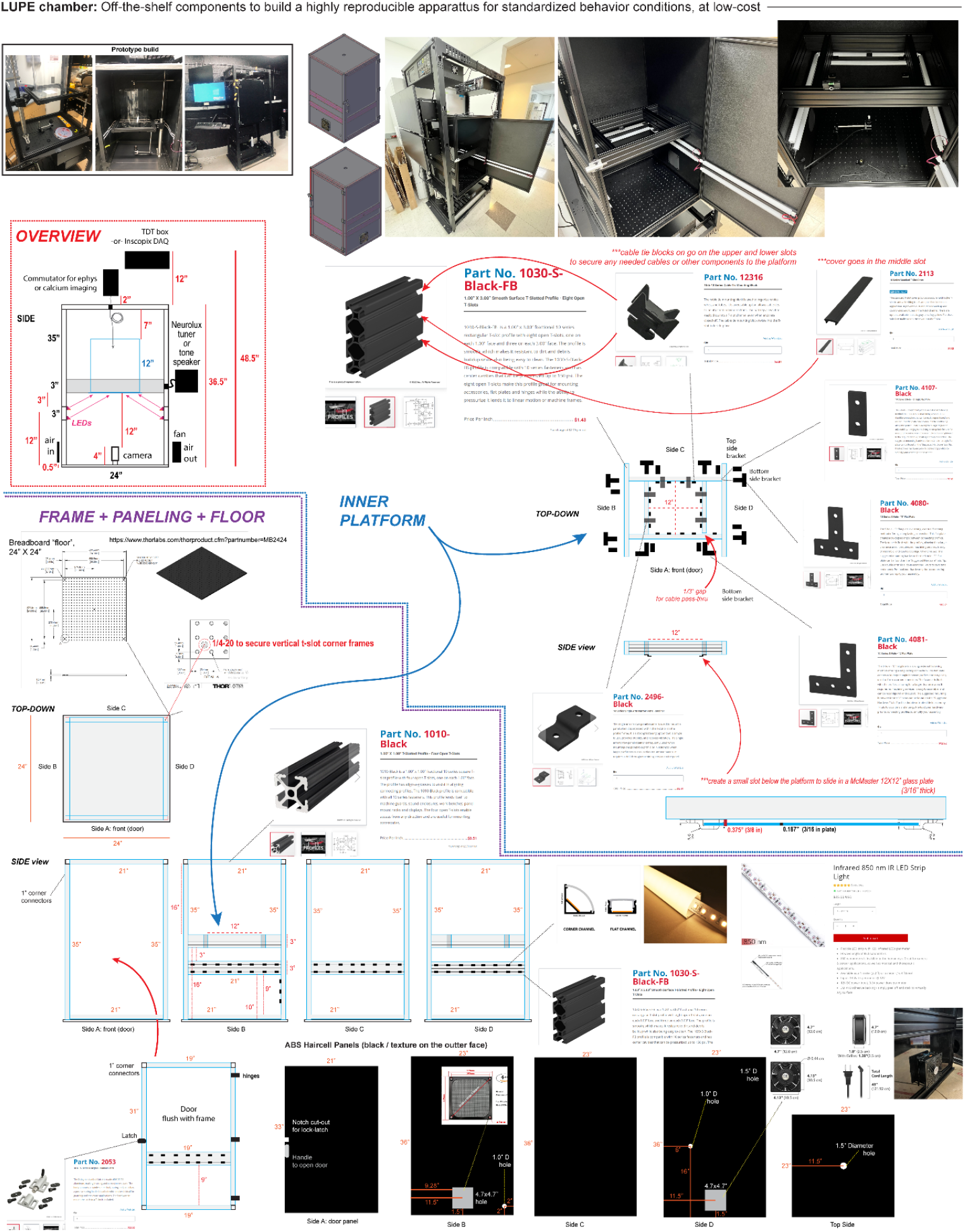
Process to create the LUPE chamber. The standardized chamber consists of commercially available components, such as plastic sheeting, 80/20 t-slot frames and brackets, and infra-red 850-nm LED strips. The exact build specifications, specifically the distance from the camera lens to the glass floor plate and location of the LEDs, provide a consistent video recording environment that can be easily replicated with and across labs. All dimensions and part numbers are listed in the figure layout. The approximate total time to build the LUPE chamber is 2.5 hours.

**Figure S7.**
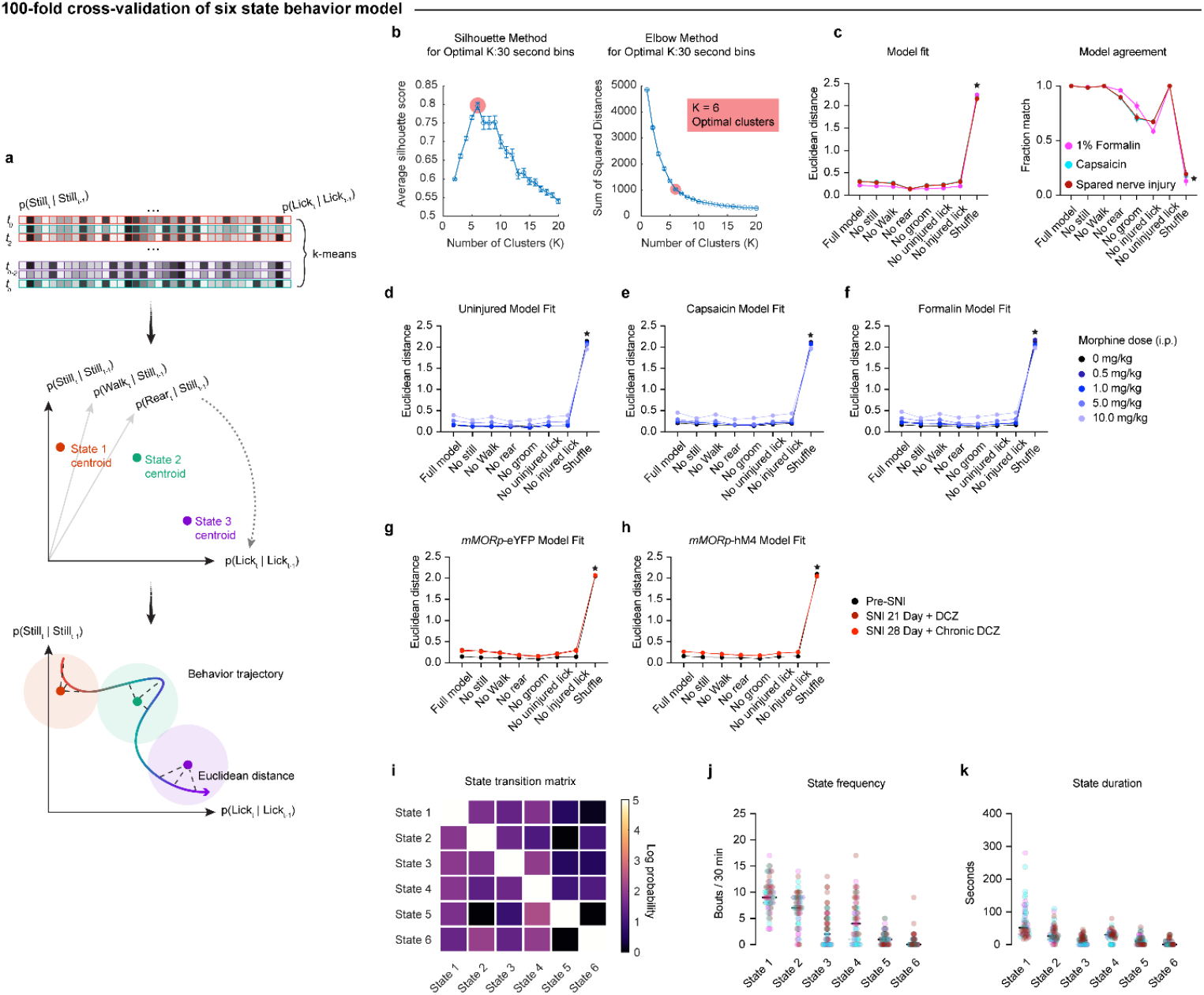
Behavior state model fits animals across models and treatments. **(a)** Cartoon of unfolded transition matrices between six spontaneous behaviors (top), which are k-means clustered in a 36-dimensional space. Model centroids are described by 36 dimensional points (unfolded transition matrices) in that state space (middle). Behavioral trajectories can be plotted in this state space, and portions of this trajectory are classified as belonging to a state by calculating the nearest centroid (Euclidean distance, bottom). **(b)** 100-fold cross-validated models with k=6 clusters maximize silhouette score and represent the “elbow” of the sum of squared distances. **(c)** Fit across full model and control models (left; permutation tests: p<0.0001 for each condition compared to n=100 shuffles) and fraction of matching classification across control models (right; permutation tests: p<0.0001 for each condition compared to n=100 shuffles) in the pain model animals used as training data. **(d-f)** Fit across full model and control models in uninjured (e), capsaicin (f), and formalin (g) mice across morphine doses (Permutation tests: p<0.0001 for each condition and treatment compared to n=100 shuffles). **(g,h)** Fit across full model and control models in mice expressing either MORp-YFP (left) or MORp-hM4Di (right) before and after spared nerve injury (SNI) with DCZ administration (Permutation tests: p<0.0001 for each condition compared to n=100 shuffles). **(i)** Log-transformed transition matrices between states. **(j,k)** Bouts (j) or mean duration (k) per session of each state in capsaicin (cyan), formalin (magenta), and SNI (dark red) animals (One-way RM ANOVA, Tukey correction, p_state, bout_ < 0.0001, p_state, duration_ < 0.0001). Stars indicate p<0.05. Bars, lines, or dots are mean; error bars and shaded areas are SEM. See **Supplementary Table 1** for statistics.

**Figure S8.**
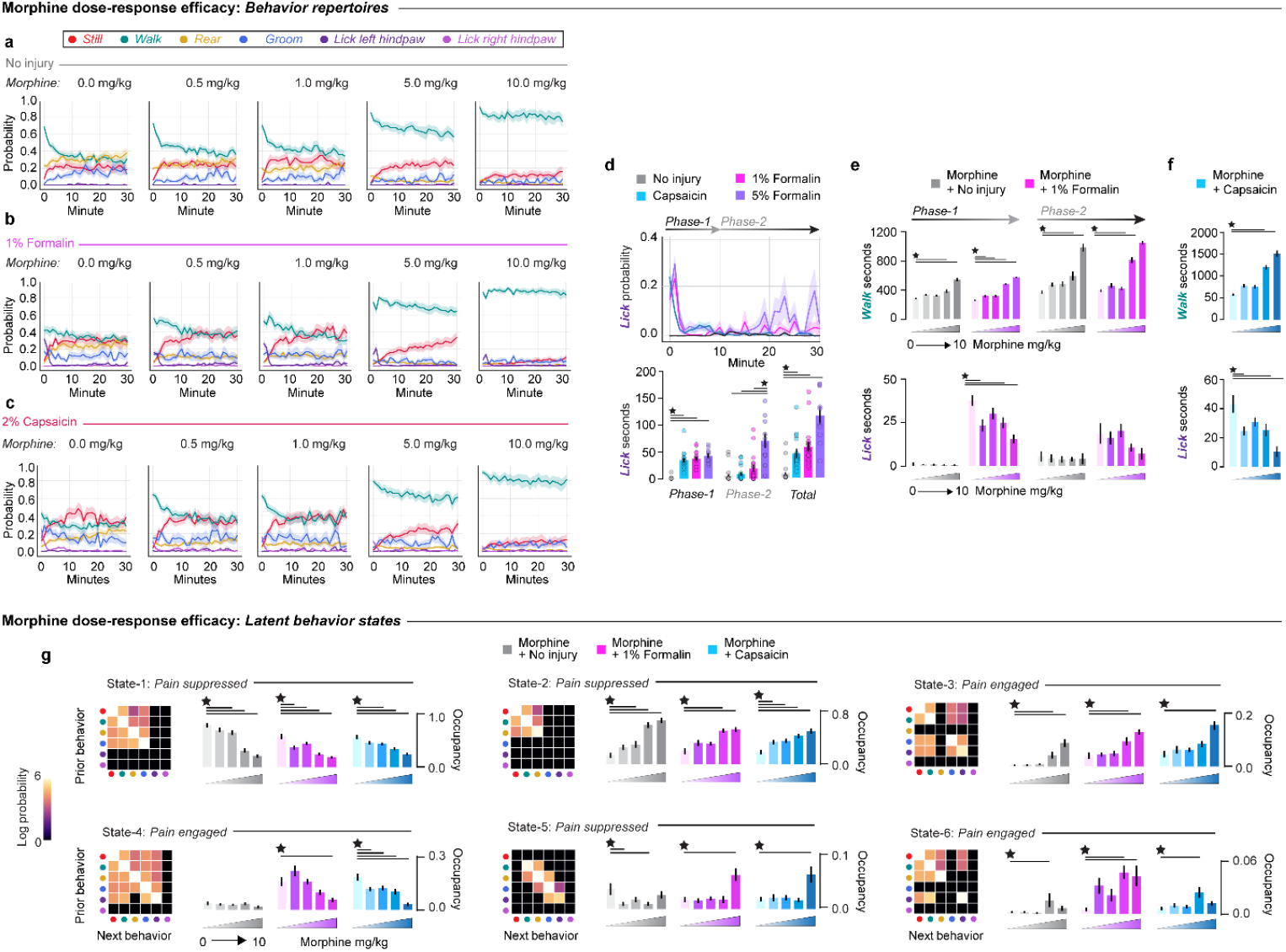
Behaviors and states are dose-dependently modulated by morphine. **(a-c)** Average behavior probabilities over 30 minutes in uninjured (a), 1% formalin-(b), and 2% capsaicin-injured (c) mice (n=20/group). **(d)** Top: Average probability of licking injured hindpaw in uninjured, 1% formalin-, 5% formalin, and 2% capsaicin injured mice over 30 minutes. Bottom: Comparison of total seconds licking between groups in the first 10 minutes (left) and second 20 minutes (right; One-way ANOVA, Tukey correction). **(e)** Dose response of morphine on average probability of walking (top) and licking (bottom) in 1% formalin-injured mice during Phase 1 (left) and Phase 2 (right; One-way ANOVA, Tukey correction). **(f)** Dose response of morphine on average probability of walking (top) and licking (bottom) in capsaicin-injured mice (One-way ANOVA, Tukey correction). **(g)** Dose response of morphine on fraction occupancy in all behavioral states in uninjured, 1% formalin, and capsaicin (One-way ANOVA, Tukey correction: p_all conditions, states 1-3_ < 0.0001, p_capsaicin and formalin, state 4_ < 0.0001, p_uninjured, State 5_ = 0.019, p_formalin, State 5_ < 0.0001, p_capsaicin, State 5_ = 0.0001, p_uninjured, State 6_ = 0.046, p_formalin, State 6_ = 0.0018, p_capsaicin, State 6_ = 0.001). Stars indicate p<0.05. Bars, lines, or dots are mean; error bars and shaded areas are SEM. See **Supplementary Table 1** for statistics.

**Figure S9.**
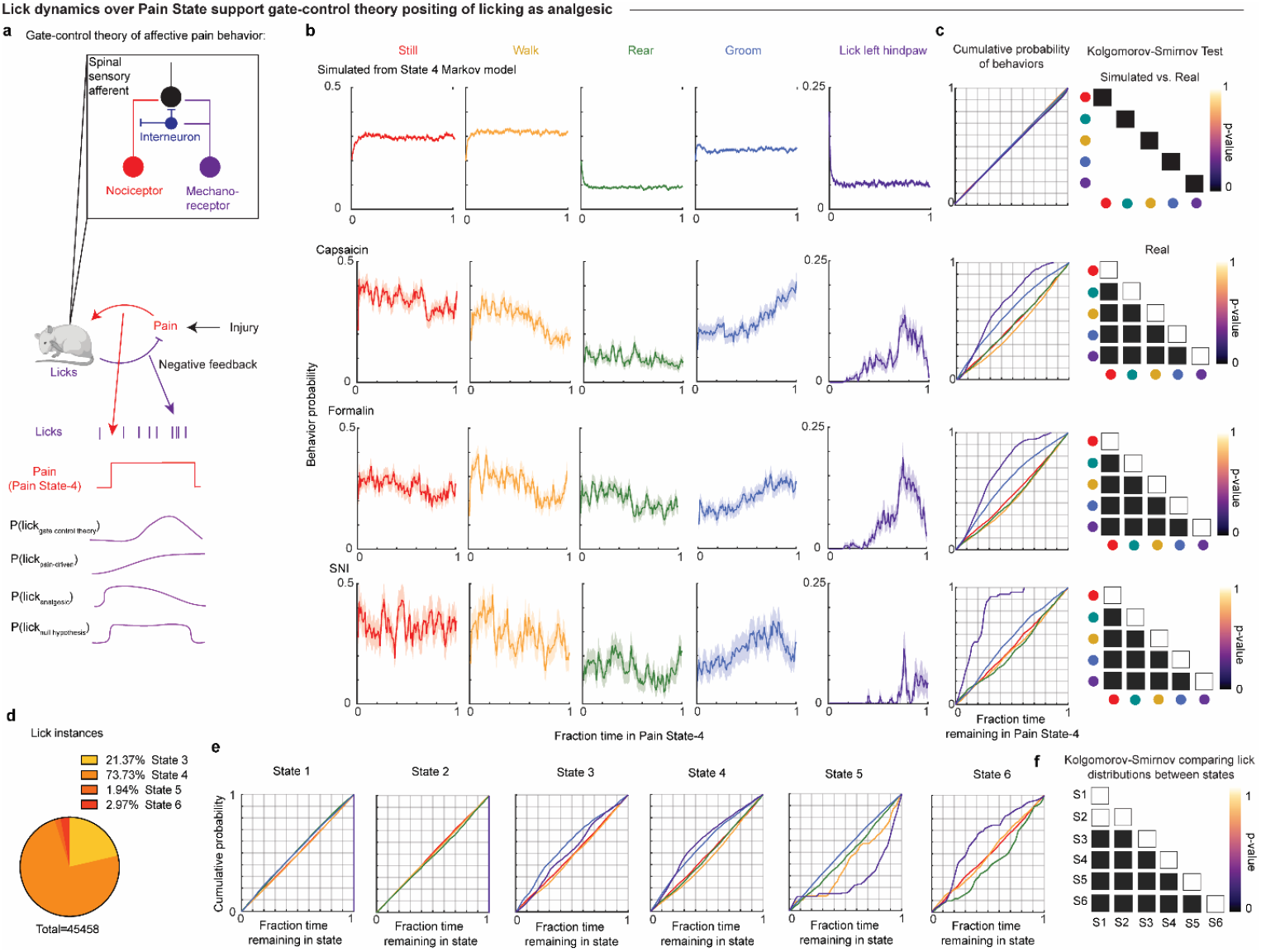
Licking is an affective-motivational pain behavior as predicted by Gate Control Theory. **(a)** Schematic of behavioral negative feedback loop of pain and recuperation and its physiological basis as posited by gate control theory. Alternative hypotheses as to how licking could be organized during pain if each of four hypotheses (pain-driven and analgesic; pain-driven but not analgesic; analgesic but not pain-driven; and neither pain-driven nor analgesic) were true. **(b)** Top row: Markov simulated probabilities of each behavior (left to right: still, walk, rear, groom, lick) averaged over initial conditions (n=100 simulations per initial condition). Bottom rows: real probabilities of each behavior in each injury condition as a function of fraction time in *Pain State-4* (n=19-20 animals per injury group, bouts pooled over animals). **(c)** Left: Cumulative distributions of each behavior as fraction time remaining in *Pain State-4*. Right: K-S test p-values comparing cumulative distributions to each other. **(d)** Distribution of licks in each state pooled across all injured animals. **(e)** Cumulative distribution of each behavior as a fraction time remaining in each state. **(f)** K-S test p-values comparing the distributions of licks between each state.

**Figure S10.**
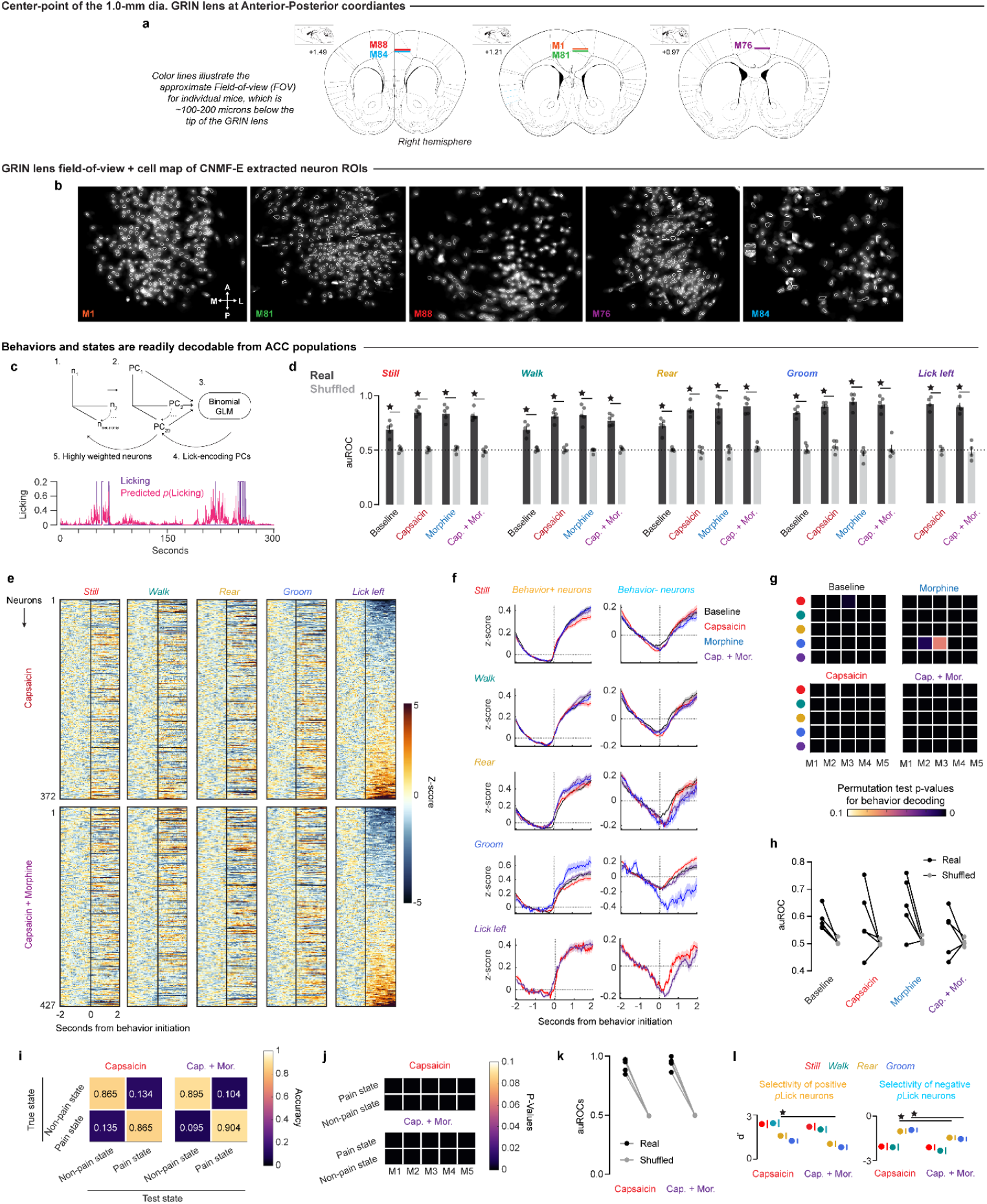
Characterizing behavior probability-encoding ACC neurons. **(a)** GRIN lens implant locations in each mouse. **(b)** Maps of registered ROIs in each mouse. **(c)** Diagram of procedure to identify behavior probability-encoding neurons from GLM over representative trace of a lick probability-encoding PC aligned to licks. **(d)** auROC of GLMs predicting each behavior across sessions (Paired t-tests, FDR<0.01: p_all comparisons_ < 0.0001). **(e)** Average PSTHs of lick-encoding neurons over onset of each behavior, sorted by Lick-evoked activity and z-scored to 1-second prior to behavior initiation. **(f)** Average activity across pooled behavior probability-encoding neurons around their preferred behavior. From left to right: neurons activated vs. inhibited by that behavior. From top to bottom: behavior-evoked activity in still, walk, rear, groom, and lick-probability encoding neurons. **(g)** Permutation test p-values within each animal testing if Fisher decoding accuracy for each behavior is significantly higher than chance in each session (n=1000 shuffles). **(h)** auROC of Fisher decoder in each animal and session in real and shuffled data. **(i)** Average Fisher decoding accuracy of pain vs. non-pain states in capsaicin (left) and capsaicin + morphine (right). **(j)** Permutation test p-values within each animal testing if Fisher decoding accuracy for states is significantly higher than chance in each session (n=1000 shuffles). **(k)** auROC of Fisher decoder in each animal and session in real and shuffled data. Stars indicate p<0.05. Bars are mean; dots are individual animals; error bars and shaded areas are SEM. See **Supplementary Table 2** for statistics.

**Figure S11.**
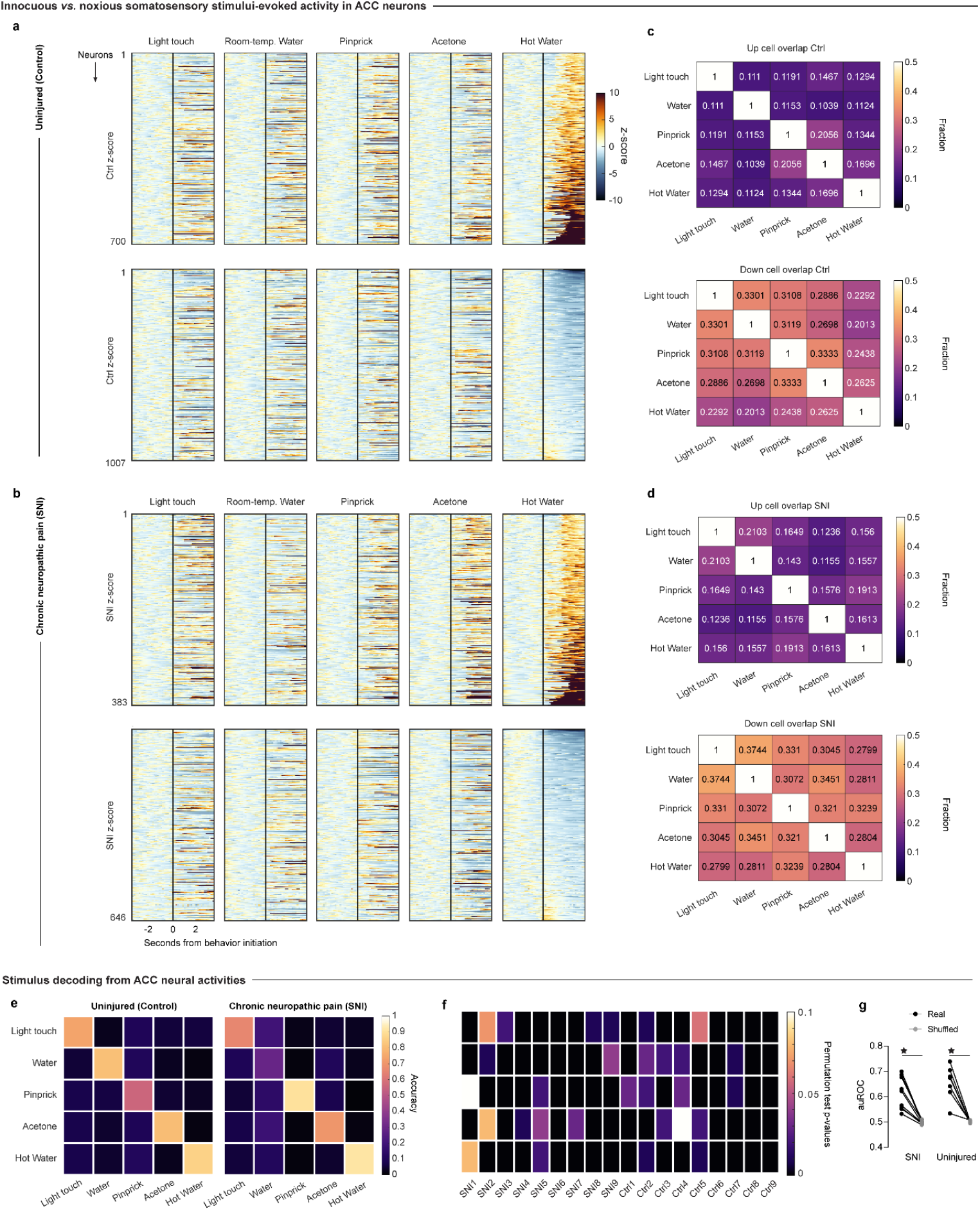
ACC neurons discriminate sensory stimuli of varying modalities, valence. **(a)** Top: Activity of hot water-activated neurons during all sensory stimuli in sorted to hot water-responses in uninjured mice (z-scored to 1s before onset; n=9 mice). Bottom: Same as top for hot water-inhibited neurons (n=9 mice). **(b)** Same as (a) in SNI mice (top: n=9 mice; bottom: n=9 mice). **(c,d)** Average fraction overlap of significantly stimulus-activated (top) or -suppressed (bottom) in uninjured (c) and SNI **(d)** mice. **(e)** Average Fisher decoding accuracy of sensory stimuli in uninjured (left) and SNI (right) mice. **(f)** Permutation test p-values within each animal testing if Fisher decoding accuracy for stimuli is significantly higher than chance in each session (n=1000 shuffles). **(g)** auROC of Fisher decoder in each animal in real and shuffled data (Paired t-tests, p_SNI_ = 0.001, p_uninjured_ < 0.0001). Stars indicate p<0.05. Bars are mean; dots are individual animals; error bars and shaded areas are SEM.

**Figure S12.**
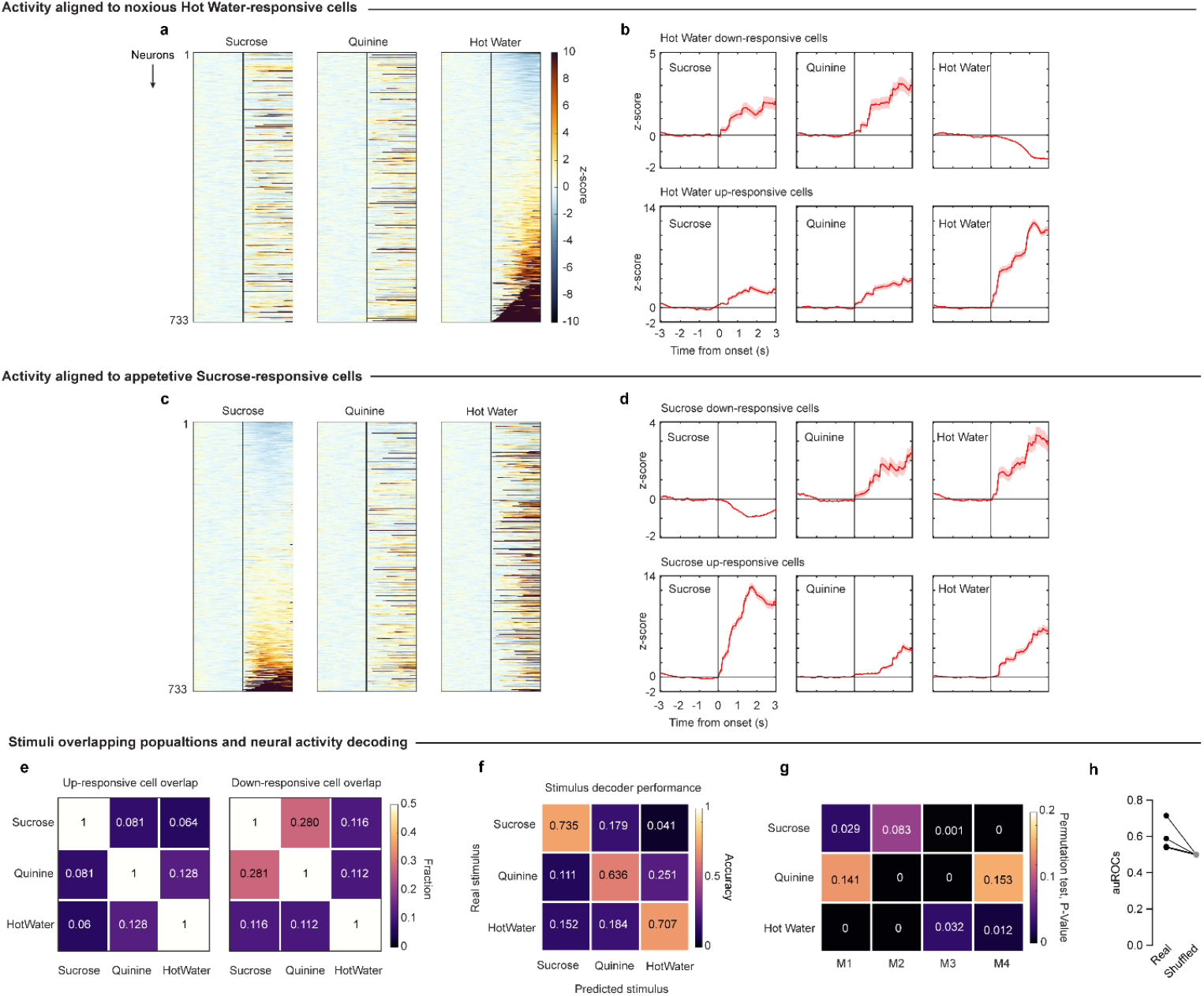
ACC neurons are selective for valence, nociception. **(a,c)** Activity of all recorded neurons across stimuli, sorted to hot water- (a) or sucrose- (c) evoked activity (n=4 mice). **(b,d)** Average activity of significantly hot water- (b) or sucrose- (d) suppressed (top) and -activated (bottom) neurons during each stimulus. **(e)** Average fraction overlap of stimulus-activated (left) and -suppressed (right) neurons. **(f)** Average Fisher decoding accuracy of sensory stimuli. **(g)** Permutation test p-values within each animal testing if Fisher decoding accuracy for stimuli is significantly higher than chance in each session (n=1000 shuffles). **(h)** auROC of Fisher decoder in each animal in real and shuffled data. Stars indicate p<0.05. Lines are mean; shaded areas are SEM.

**Figure S13.**
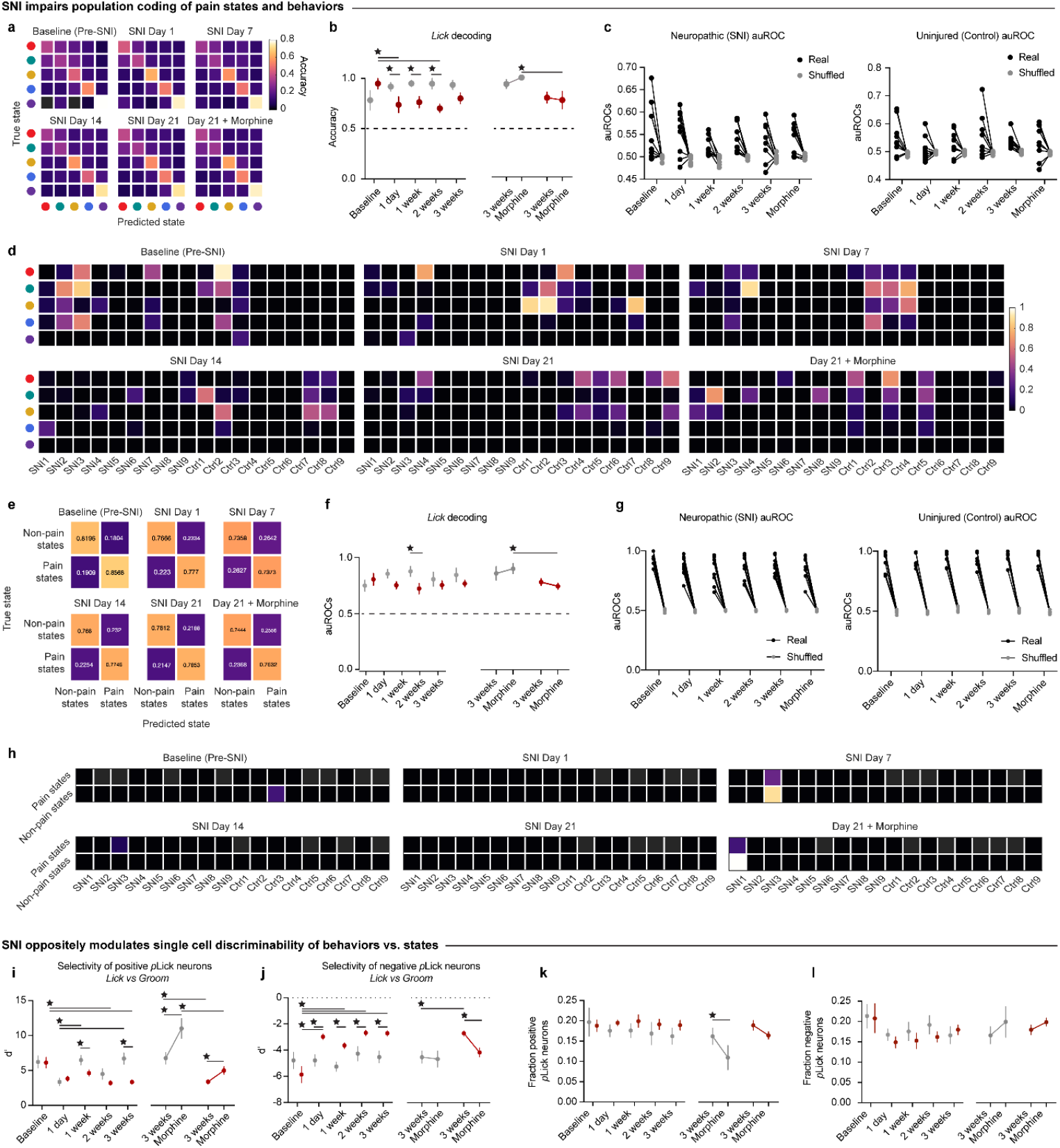
Chronic pain impairs encoding of behaviors and states in ACC. **(a,e)** Average Fisher decoding accuracy of behaviors (a) and pain states (e) in SNI mice in each session. **(b,f)** Left: Average Fisher decoding accuracy of behaviors (b) and pain states (f) in SNI (n=9) and uninjured mice (n=9) before and after SNI or anesthesia (Mixed effects model, Tukey correction: p_lick, interaction_ = 0.0089, p_state, interaction_ = 0.067). Right: Same as left at three weeks post-SNI or anesthesia before and after morphine (Mixed effects model, Tukey correction: p_lick, interaction_ = 0.041, p_state, injury_ = 0.025). **(c,g)** auROC of behavior (c) or state (g) Fisher decoder in each SNI (left) or uninjured (right) animal and session in real and shuffled data (Mixed effects model, Tukey correction: p_sni and control, lick and state, real vs shuffle_ < 0.0001). **(d,h)** Permutation test p-values within each animal testing if Fisher decoding accuracy for behavior (d) or state (h) is significantly higher than chance in each session (n=1000 shuffles). **(i)** Left: Single cell discriminability for lick in positive *p*Lick neurons before and after SNI or anesthesia (Two-Way ANOVA, Tukey correction: p_interaction_ = 0.0044). Right: Same as left at three weeks post-SNI or anesthesia before and after morphine (Two-Way ANOVA, Tukey correction: p_interaction_ = 0.048). **(j)** Same as (g) for negative *p*Lick neurons (Two-Way ANOVA, Tukey correction: p_left, interaction_ = 0.0031, p_right, injury_ = 0.0016, p_right, treatment_ = 0.029). **(k)** Left: Fraction of positive *p*Lick neurons before and after SNI or anesthesia (Mixed effects model, Tukey correction). Right: Same as left at three weeks post-SNI or anesthesia before and after morphine (Mixed effects model, Tukey correction: p_interaction_ = 0.0024). **(l)** Same as (k) for negative *p*Lick neurons (Mixed effects model, Tukey correction: p_right, treatment_ = 0.046). Stars indicate p<0.05. Bars, lines, or dots are mean; error bars and shaded areas are SEM. See **Supplementary Table 3** for statistics.

**Figure S14.**
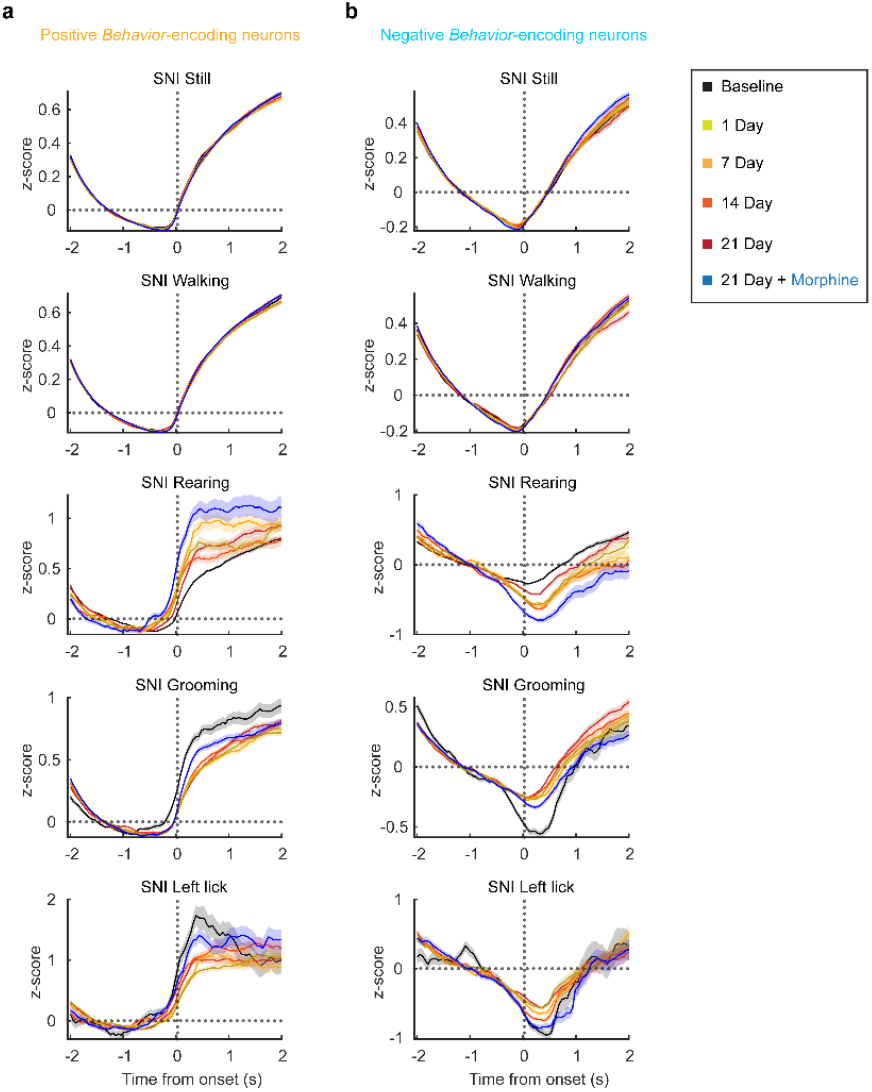
Chronic pain and morphine alter evoked activity in ACC behavior probability-encoding neurons. **(a,b)** Average behavior-evoked activity in neurons encoding that behavior which increase (left) or decrease (right) activity upon bout onset (z-score from -2 to -1 seconds before onset). From top to bottom: Still, walk, rear, groom, and lick-encoding neurons at still, walk, rear, groom, and lick onsets, respectively.

**Figure S15.**
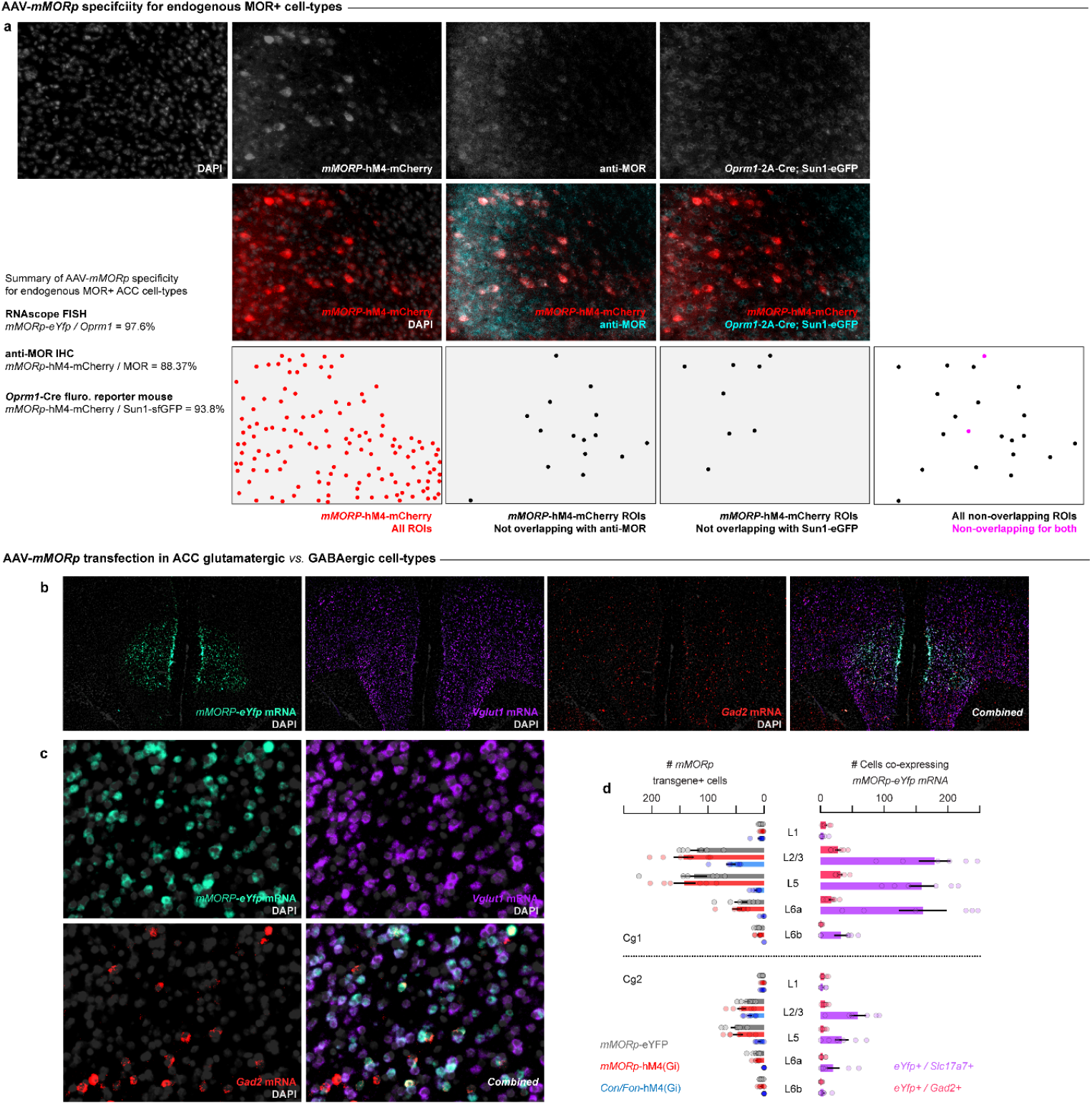
AAV-m*MORp* transfection in ACC *Oprm1*+ cell-types. **(a)** AAV-mMORp-hM4-mCherry DREADD expression in Cg1 ACC of *Oprm1*-2A-Cre; Sun1sf-GFP mice co-stained for anti-MOR immunoreactivity. **(b)** RNAscope FISH for eYfp (AAV-mMORp-eYFP), Slc17a7 (Vesicular glutamate transporter 1, VGLUT1) and Gad2 (Glutamic acid decarboxylase 2, GAD2). **(c)** 20X images of panel B. **(d)** HALO quantification of *eYfp* mRNA overlaps with *Slac17a7*+ and *Gad2*+ cells. N=6 separate hemisphere injection sites from n=3 mice.

**Figure S16.**
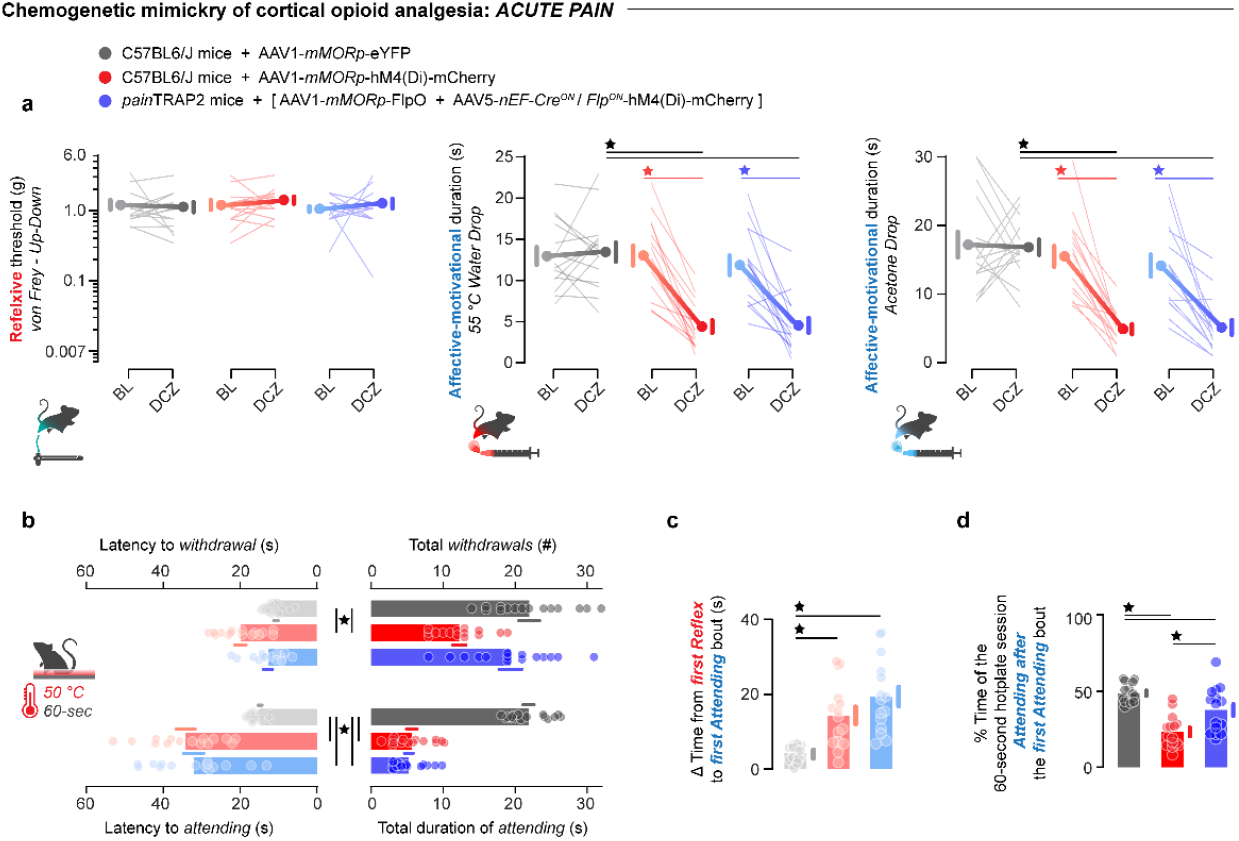
Opioid analgesia mimicry via chemogenetic inhibition of nociceptive MOR+ ACC neurons in acute pain. Chemogenetic inhibition of ACC MOR+ cells (Red: *mMORp*-hM4, N=15), ACC nociceptive MOR+ cells (Blue: *mMORp*-FlpO + *Cre*^*ON*^*/Flp*^*ON*^-hM4 in *pain*TRAP mice, N=15), or control/non-inhibited (Gray: *mMORp*-eYFP, N=15). **(a)** Reflexive withdrawal thresholds and affective-motivational response duration to noxious hot (55°C hot water) and cold (acetone) stimuli at baseline and after administration of DCZ. (Two-way ANOVA + Tukey: von Frey P_interaction_ = 0.6451, acetone P_interaction_ = 0.003, hot water P_interaction_ <0.0001) **(b)** Latency to withdraw (P <0.0001), total withdrawals (P<0.0001), latency to attend (P<0.0001), and total duration of attending (P<0.0001) induced in 60 seconds on a 50°C inescapable hot plate after DCZ. (One-way ANOVA + Tukey for all panels) **(c)** Duration of time between the first reflexive withdrawal and the first bout of attending behaviors on the inescapable hot plate after DCZ. (One-way ANOVA + Tukey, P<0.0001) **(d)** The proportion of the trial engaging in attending behaviors after the first attending bout on the inescapable hot plate after DCZ. (One-way ANOVA + Tukey, P<0.0001). For detailed statistics, see **Supplemental Table 4**. ⋆ = P < 0.05. Errors bars = s.e.m.

**Figure S17.**
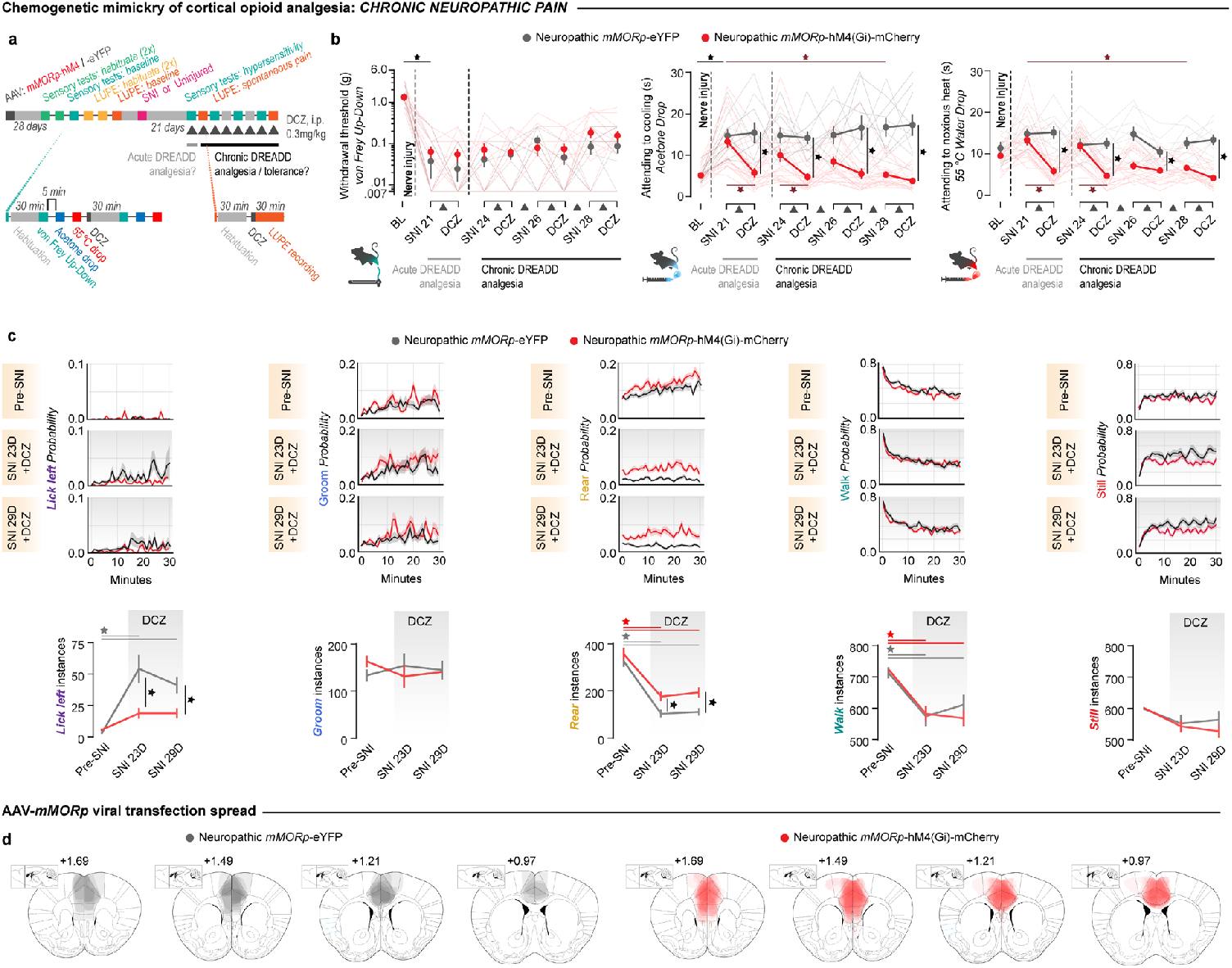
Opioid analgesia mimicry via chemogenetic inhibition of MOR+ ACC neurons in chronic pain. **(a)** Design for chemogenetic inhibition of ACC MOR+ cell-types in neuropathic pain mice (N=19 *mMORp*-eYFP mice, N=30 *mMORp*-hM4; equal sexes). **(b)** Effects of acute and chronic DCZ to engage *mMORp*-hM4 signaling on mechanical hypersensitivity (*left*, p_interaction_ =0.9873), cold allodynia (*middle*, P_interaction_ <0.0001), and heat hyperalgesia (*right*, p_interaction_ <0.0001). (Two-way ANOVA + Tukey for all panels) **(c)** Effects of mMORp-hM4 inhibition on LUPE-scored neuropathic pain spontaneous behavior repertoires pre and 23/29 days post-SNI. (Two-way ANOVA + Tukey for all panels: lick p_interaction_=0.0012, groom p_interaction_=0.3668, rear p_interaction_=0.4428, walk p_interaction_=0.5024, still p_interaction_=0.6690). **(d)** Viral spread maps for AAV-*mMORp*-eYFP vs. AAV-*mMORp*-hM4-mCherry. For detailed statistics, see **Supplemental Table 4**. ⋆ = P < 0.05. Errors bars = s.e.m.

**Figure S18.**
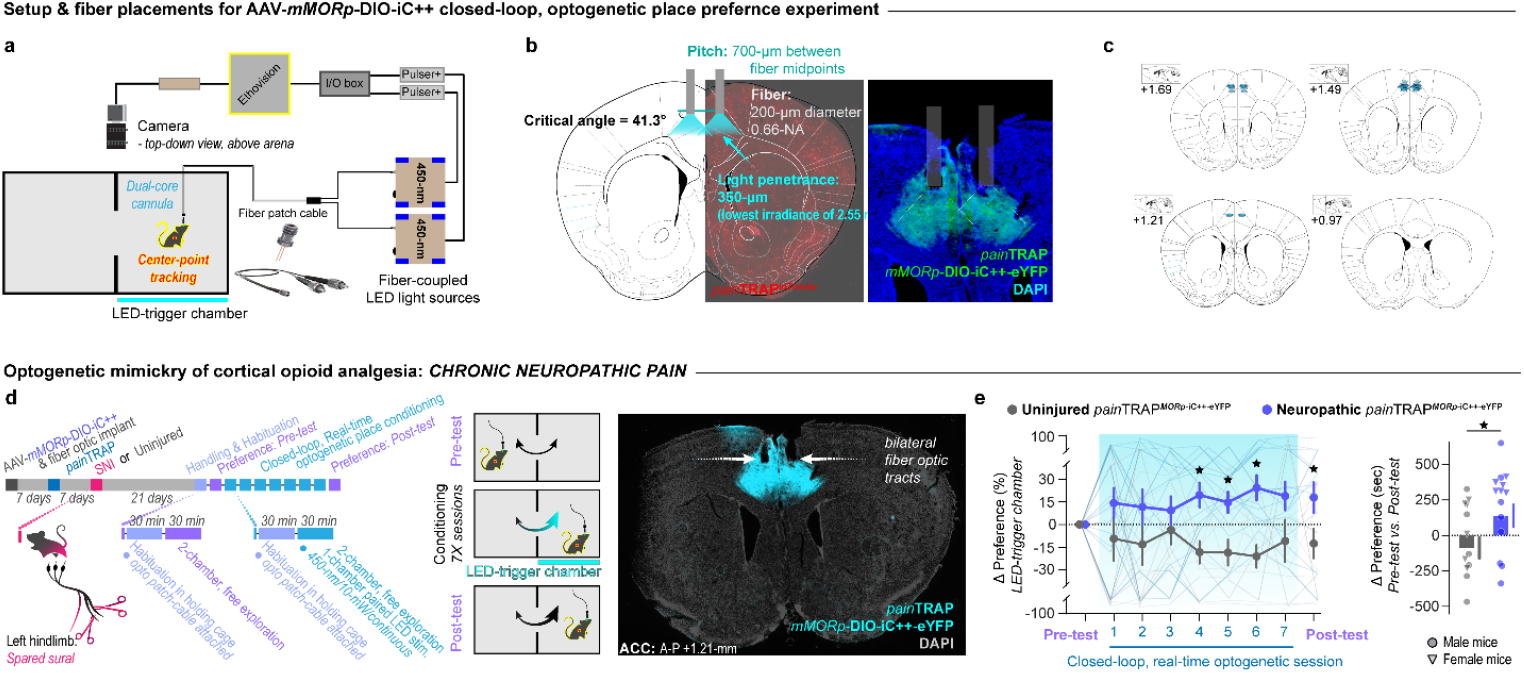
Optogenetic inhibition of ACC MOR+ cells induces a conditioned place preference in injured mice. **(a)** Real-time place preference setup with Ethovision triggering an external LED system to administer blue light to the ACC of *pain*TRAP2 mice transfected with the inhibitory chloride channel opsin, AAV-mMORp-DIO-iC++. **(b)** Bilateral fiber optic design for LED light penetrance to the ACC (left), with a representative image of the cannula tracks over the dorsal Cg1 ACC expressing iC++ in *pain*TRAP neurons. **(c)** Map of optic fiber placements across mice. **(d)** Design for optogenetic inhibition of ACC chronic nociceptive-MOR+ cell-types in a real-time place preference assay. **(e)** Cross-day changes in real-time, within-session self-administered inhibition via volitional selection to remain in an LED-triggered chamber, and subsequent conditioned place preference assessment with no LED exposure (N=12 Uninjured mice, N=14 SNI mice). (Left panel, Two-way ANOVA with Tukey correction: p_virus_=0.0066. Right panel, unpaired t-test: p=0.0466). For detailed statistics, see **Supplemental Table 5**. ⋆ = P < 0.05. Errors bars = s.e.m.

## Notes

https://github.com/justin05423/LUPE-2.0-AnalysisPackage/

### Summary of Updates

-Structural refinement across the Introduction to Discussion to emphasize the conceptual significance of cortical gene therapy for chronic pain. -Condensed the Main Figures from 5 to 4, and the Supplementary Figures from 31 to 18. -Focused data presentation by removing or relocating peripheral elements and replication data (e.g., evoked pain measures, opioid cell-type FISH and single nuclei RNAseq) to the Supplement. -Enhanced visualization, with simplified figures, standardized metrics (e.g., LUPE-based deep-learning modeling for spontaneous pain behaviors), and new schematics to clearly convey each figure message. -Expanded methods and figure legends to ensure reproducibility. -New experiments, including: o Calcium imaging with synchronized LUPE tracking across chronic pain development (Figure 3). o A Mouse Pain Scale to quantify analgesia and test our gene therapy (All figures). o Imaging of single-neuron responses to pain and appetitive/aversive stimuli (Supp. Figs. 11 - 12).

## References

1. Kimmey, B. A., McCall, N. M., Wooldridge, L. M., Satterthwaite, T. D. & Corder, G. Engaging endogenous opioid circuits in pain affective processes. J Neurosci Res 1–33 (2020) doi:10.1002/jnr.24762.

2. Wiech, K. Deconstructing the sensation of pain: The influence of cognitive processes on pain perception. Science (1979) 354, 584–587 (2016).

3. Tan, L. L. & Kuner, R. Neocortical circuits in pain and pain relief. Nature Reviews Neuroscience vol. 22 458–471 Preprint at 10.1038/s41583-021-00468-2 (2021).

4. Corder, G., Castro, D. C., Bruchas, M. R. & Scherrer, G. Endogenous and Exogenous Opioids in Pain. Annu Rev Neurosci 453–73 (2018) doi:10.1146/annurev-neuro-080317-061522.

5. Woolf, C. J. Capturing Novel Non-opioid Pain Targets. Biol Psychiatry 87, 74–81 (2020).

6. Grosser, T., Woolf, C. J. & FitzGerald, G. A. Time for nonaddictive relief of pain. Science (1979) 355, 1026–1027 (2017).

7. Bliss, T. V. P., Collingridge, G. L., Kaang, B. K. & Zhuo, M. Synaptic plasticity in the anterior cingulate cortex in acute and chronic pain. Nature Reviews Neuroscience vol. 17 485–496 Preprint at 10.1038/nrn.2016.68 (2016).

8. Rainville, P. Pain Affect Encoded in Human Anterior Cingulate But Not Somatosensory Cortex. Science (1979) 277, 968–971 (1997).

9. Apkarian, A. V., Bushnell, M. C., Treede, R. D. & Zubieta, J. K. Human brain mechanisms of pain perception and regulation in health and disease. European Journal of Pain 9, 463–484 (2005).

10. Cauda, F. et al. Shared ‘core’ areas between the pain and other task-related networks. PLoS One 7, (2012).

11. Barthas, F. et al. The anterior cingulate cortex is a critical hub for pain-induced depression. Biol Psychiatry 77, 236–245 (2015).

12. Shackman, A. J. et al. The integration of negative affect, pain and cognitive control in the cingulate cortex. Nat Rev Neurosci 12, 154–167 (2011).

13. Simons, L. E., Elman, I. & Borsook, D. Psychological processing in chronic pain: A neural systems approach. Neurosci Biobehav Rev 39, 61–78 (2014).

14. Seymour, B. Pain : A Precision Signal for Reinforcement Learning and Control. Neuron 101, 1029–1041 (2019).

15. Economides, M., Guitart-Masip, M., Kurth-Nelson, Z. & Dolan, R. J. Anterior Cingulate Cortex Instigates Adaptive Switches in Choice by Integrating Immediate and Delayed Components of Value in Ventromedial Prefrontal Cortex. Journal of Neuroscience 34, 3340–3349 (2014).

16. Price, D. D. Psychological and Neural Mechanisms of the Affective Dimension of Pain. Science (1979) 288, 1769–1772 (2000).

17. Foltz, E. L. & Lowell, E. W. Pain ‘Relief’ by Cingulotomy. J Neurosurg 19, 89–100 (1961).

18. Foltz, E. L. & White, L. E. Pain “Relief” by Frontal Cingulumotomy. J Neurosurg 19, 89–100 (1962).

19. Grahek, N. Feeling Pain and Being in Pain. (MIT Press, 2007).

20. Hurt, R. W. & Ballantine, H. T. Stereotactic anterior cingulate lesions for persistent pain: a report on 68 cases. Clin Neurosurg 21, 334–351 (1974).

21. Johansen, J. P., Fields, H. L. & Manning, B. H. The affective component of pain in rodents: Direct evidence for a contribution of the anterior cingulate cortex. Proc Natl Acad Sci U S A 98, 8077–8082 (2001).

22. LaGraize, S. C., Labuda, C. J., Rutledge, M. A., Jackson, R. L. & Fuchs, P. N. Differential effect of anterior cingulate cortex lesion on mechanical hypersensitivity and escape/avoidance behavior in an animal model of neuropathic pain. Exp Neurol 188, 139–148 (2004).

23. Zubieta, J. K. et al. Regional Mu opioid receptor regulation of sensory and affective dimensions of pain. Science (1979) 293, 311–315 (2001).

24. Zubieta, J.-K. et al. Regulation of Human Affective Responses by Anterior Cingulate and Limbic ?-Opioid Neurotransmission.

25. Navratilova, X. E. et al. Endogenous Opioid Activity in the Anterior Cingulate Cortex Is Required for Relief of Pain. Journal of Neuroscience 35, 7264–7271 (2015).

26. LaGraize, S. C., Borzan, J., Peng, Y. B. & Fuchs, P. N. Selective regulation of pain affect following activation of the opioid anterior cingulate cortex system. Exp Neurol 197, 22–30 (2006).

27. Hu, T. T. et al. Activation of the Intrinsic Pain Inhibitory Circuit from the Midcingulate Cg2 to Zona Incerta Alleviates Neuropathic Pain. J Neurosci 39, 9130–9144 (2019).

28. Huang, J. et al. A neuronal circuit for activating descending modulation of neuropathic pain. Nat Neurosci 22, 1659–1668 (2019).

29. Gadotti, V. M., Zhang, Z., Huang, J. & Zamponi, G. W. Analgesic effects of optogenetic inhibition of basolateral amygdala inputs into the prefrontal cortex in nerve injured female mice. Mol Brain 12, 10–13 (2019).

30. Gu, L. et al. Pain inhibition by optogenetic activation of specific anterior cingulate cortical neurons. PLoS One 10, 1–17 (2015).

31. Sellmeijer, J. et al. Hyperactivity of Anterior Cingulate Cortex Areas 24a/24b Drives Chronic Pain-Induced Anxiodepressive-like Consequences. The Journal of Neuroscience 38, 3102–3115 (2018).

32. Zhou, H. et al. Inhibition of the prefrontal projection to the nucleus accumbens enhances pain sensitivity and affect. Front Cell Neurosci 12, 1–13 (2018).

33. Meda, K. S. et al. Microcircuit Mechanisms through which Mediodorsal Thalamic Input to Anterior Cingulate Cortex Exacerbates Pain-Related Aversion. Neuron 102, 944-959.e3 (2019).

34. Qi, X. et al. A nociceptive neuronal ensemble in the dorsomedial prefrontal cortex underlies pain chronicity. Cell Rep 41, (2022).

35. Zhang, Q. et al. Chronic pain induces generalized enhancement of aversion. (2017) doi:10.7554/eLife.25302.001.

36. Motzkin, J. C., Kanungo, I., D’Esposito, M. & Shirvalkar, P. Network targets for therapeutic brain stimulation: towards personalized therapy for pain. Frontiers in Pain Research 4, (2023).

37. Riis, T. S., Feldman, D. A., Losser, A. J., Okifuji, A. & Kubanek, J. Noninvasive targeted modulation of pain circuits with focused ultrasonic waves. Pain (2024) doi:10.1097/j.pain.0000000000003322.

38. Chuong, A. S. et al. Noninvasive optical inhibition with a red-shifted microbial rhodopsin. Nat Neurosci 17, 1123–1129 (2014).

39. Ovsepian, S. V. & Waxman, S. G. Gene therapy for chronic pain: emerging opportunities in target-rich peripheral nociceptors. Nat Rev Neurosci 24, 252–265 (2023).

40. Mata, M., Hao, S. & Fink, D. J. Applications of Gene Therapy to the Treatment of Chronic Pain. 42–48 (2008).

41. Salimando, G. J. et al. Human OPRM1 and murine Oprm1 promoter driven viral constructs for genetic access to μ-opioidergic cell types. Nat Commun 14, (2023).

42. Corder, G. et al. An amygdalar neural ensemble that encodes the unpleasantness of pain. Science (1979) 363, 276–281 (2019).

43. Tan, L. L. et al. A pathway from midcingulate cortex to posterior insula gates nociceptive hypersensitivity. Nat Neurosci 20, 1591–1601 (2017).

44. Ntamati, N. R., Acuña, M. A. & Nevian, T. Pain-induced adaptations in the claustro-cingulate pathway. Cell Rep 42, (2023).

45. Mansour, A. et al. Mu, delta, and kappa opioid receptor mRNA expression in the rat CNS: An in situ hybridization study. Journal of Comparative Neurology 350, 412–438 (1994).

46. Baumgärtner, U. et al. High opiate receptor binding potential in the human lateral pain system. Neuroimage 30, 692–699 (2006).

47. Vogt, B., Wiley, R. & Jensen, E. Localization of Mu and delta opioid receptors to anterior cingulate afferents and projection neurons and input/output model of Mu regulation. Exp Neurol 83–92 (1995).

48. Mansour, A., Fox, C. A., Thompson, R. C., Akil, H. & Watson, S. J. mu-Opioid receptor mRNA expression in the rat CNS: comparison to mu-receptor binding. Brain Res 643, 245–65 (1994).

49. Kupers, R. C., Konings ‘, H., Adriaensen, H. & Gybels, J. M. Clinical Section Morphine Differentially Affects the Sensory and Affective Pain Ratings in Neurogenic and Idiopathic Forms of Pain. Pain vol. 47 http://journals.lww.com/pain (1991).

50. Zamfir, M. et al. Distinct and sex-specific expression of mu opioid receptors in anterior cingulate and somatosensory S1 cortical areas. Pain 164, 703–716 (2023).

51. Shields, S. D., Eckert, W. A. & Basbaum, A. I. Spared Nerve Injury Model of Neuropathic Pain in the Mouse: A Behavioral and Anatomic Analysis. Journal of Pain 4, 465–470 (2003).

52. Finnerup, N. B., Kuner, R. & Jensen, T. S. Neuropathic pain: Frommechanisms to treatment. Physiol Rev 101, 259–301 (2021).

53. Mansour, A., Akil, H. & Watson, S. Mu, delta, and kappa opioid receptor mRNA expression in the rat CNS: an in situ hybridization study. J Comp Neurol 412–38 (1994).

54. Cummings, K. A., Bayshtok, S., Dong, T. N., Kenny, P. J. & Clem, R. L. Control of fear by discrete prefrontal GABAergic populations encoding valencespecific information. Neuron 110, 3036-3052.e5 (2022).

55. Cole, R. H., Moussawi, K. & Joffe, M. E. Opioid modulation of prefrontal cortex cells and circuits. Neuropharmacology vol. 248 Preprint at 10.1016/j.neuropharm.2024.109891 (2024).

56. Koopmans, F. et al. SynGO: An Evidence-Based, Expert-Curated Knowledge Base for the Synapse. Neuron 103, 217-234.e4 (2019).

57. Alvarado, S. et al. Peripheral Nerve Injury is Accompanied by Chronic Transcriptome-Wide Changes in the Mouse Prefrontal Cortex. Mol Pain 9, 1744-8069-9–21 (2013).

58. Qiu, X. T. et al. Transcriptomic and proteomic profiling of the anterior cingulate cortex in neuropathic pain model rats. Front Mol Neurosci 16, (2023).

59. Topham, L. et al. The transition from acute to chronic pain: dynamic epigenetic reprogramming of the mouse prefrontal cortex up to 1 year after nerve injury. Pain 161, 2394–2409 (2020).

60. Renthal, W. et al. Transcriptional Reprogramming of Distinct Peripheral Sensory Neuron Subtypes after Axonal Injury. Neuron 108, 128-144.e9 (2020).

61. Üçeyler, N. et al. Cortical Binding Potential of Opioid Receptors in Patients With Fibromyalgia Syndrome and Reduced Systemic Interleukin-4 Levels – A Pilot Study. Front Neurosci 14, (2020).

62. Harris, R. E. et al. Decreased central μ-opioid receptor availability in fibromyalgia. Journal of Neuroscience 27, 10000–10006 (2007).

63. Thompson, S. J. et al. Chronic neuropathic pain reduces opioid receptor availability with associated anhedonia in rat. Pain 159, 1856–1866 (2018).

64. Corder, G. et al. Loss of μ-opioid receptor signaling in nociceptors, and not spinal microglia, abrogates morphine tolerance without disrupting analgesic efficacy. Nat Med 23, 164–173 (2017).

65. Cobos, E. J. et al. Inflammation-induced decrease in voluntary wheel running in mice: A nonreflexive test for evaluating inflammatory pain and analgesia. Pain 153, 876–884 (2012).

66. Wojick, J. A. et al. A nociceptive amygdala-striatal pathway for chronic pain aversion. bioRxiv (2024) doi:10.1101/2024.02.12.579947.

67. Liu, S. et al. Neural basis of opioid-induced respiratory depression and its rescue. doi:10.1073/pnas.2022134118/-/DCSupplemental.

68. Jenny He, X. et al. Convergent, functionally independent signaling by mu and delta opioid receptors in hippocampal parvalbumin interneurons. doi:10.7554/eLife.

69. Bove, G. Mechanical sensory threshold testing using nylon monofilaments: The pain field’s ‘Tin Standard’. Pain vol. 124 13–17 Preprint at 10.1016/j.pain.2006.06.020 (2006).

70. King, T. & Porreca, F. Preclinical Assessment of Pain: Improving Models in Discovery Research. http://www.springer.com/series/7854.

71. Vierck, C. J., Hansson, P. T. & Yezierski, R. P. Clinical and pre-clinical pain assessment: Are we measuring the same thing? Pain vol. 135 7–10 Preprint at 10.1016/j.pain.2007.12.008 (2008).

72. King, T. et al. Unmasking the tonic-aversive state in neuropathic pain. Nat Neurosci 12, 1364–1366 (2009).

73. Johansen, J. P. & Fields, H. L. Glutamatergic activation of anterior cingulate cortex produces an aversive teaching signal. Nat Neurosci 7, 398– 403 (2004).

74. Yang, J., Xie, Y.-F., Smith, R., Ratté, S. & Prescott, S. A. Discordance between preclinical and clinical testing of Na V 1.7-selective inhibitors for pain. Pain 166, 481–501 (2025).

75. Nath, T. et al. Using DeepLabCut for 3D markerless pose estimation across species and behaviors. Nat Protoc 14, 2152–2176 (2019).

76. Tillmann, J. F., Hsu, A. I., Schwarz, M. K. & Yttri, E. A. A-SOiD, an activelearning platform for expert-guided, data-efficient discovery of behavior. Nat Methods (2024) doi:10.1038/s41592-024-02200-1.

77. Hsu, A. I. & Yttri, E. A. B-SOiD, an open-source unsupervised algorithm for identification and fast prediction of behaviors. Nat Commun 12, 1–13 (2021).

78. Hunskaar, S., Fasmer, O. B. & Hole, K. Formalin Test in Mice, a Useful Technique for Evaluating Mild Analgesics. Journal of Neuroscience Methods vol. 14 (1985).

79. Friard, O. & Gamba, M. BORIS: a free, versatile open-source event-logging software for video/audio coding and live observations. Methods Ecol Evol 7, 1325–1330 (2016).

80. Acuña, M. A., Kasanetz, F., De Luna, P., Falkowska, M. & Nevian, T. Principles of nociceptive coding in the anterior cingulate cortex. Proc Natl Acad Sci U S A 120, (2023).

81. Nagai, Y. et al. Deschloroclozapine, a potent and selective chemogenetic actuator enables rapid neuronal and behavioral modulations in mice and monkeys. Nat Neurosci 23, 1157–1167 (2020).

82. Stegemann, A. et al. Prefrontal engrams of long-term fear memory perpetuate pain perception. Nat Neurosci 26, 820–829 (2023).

83. Qi, X. et al. A nociceptive neuronal ensemble in the dorsomedial prefrontal cortex underlies pain chronicity. Cell Rep 41, (2022).

84. Smith, M. L., Asada, N. & Malenka, R. C. Anterior cingulate inputs to nucleus accumbens control the social transfer of pain and analgesia. Science (1979) 371, 153–159 (2021).

85. Wiech, K. et al. Anterior insula integrates information about salience into perceptual decisions about pain. Journal of Neuroscience 30, 16324– 16331 (2010).

86. Faig, C. A. et al. Claustrum projections to the anterior cingulate modulate nociceptive and pain-associated behavior. Curr Biol 34, 1987-1995.e4 (2024).

87. Meda, K. S. et al. Microcircuit Mechanisms through which Mediodorsal Thalamic Input to Anterior Cingulate Cortex Exacerbates Pain-Related Aversion. Neuron 102, 944-959.e3 (2019).

88. Wilson, H. D., Uhelski, M. L. & Fuchs, P. N. Examining the role of the medial thalamus in modulating the affective dimension of pain. Brain Res 1229, 90–99 (2008).

89. Neugebauer, V., Li, W., Bird, G. C. & Han, J. S. The amygdala and persistent pain. Neuroscientist 10, 221–234 (2004).

90. Cheriyan, J. & Sheets, P. L. Altered Excitability and Local Connectivity of mPFC-PAG Neurons in a Mouse Model of Neuropathic Pain. The Journal of Neuroscience 38, 4829–4839 (2018).

91. Valentinova, K., Acuña, M. A., Ntamati, N. R., Nevian, N. E. & Nevian, T. An amygdala-to-cingulate cortex circuit for conflicting choices in chronic pain. Cell Rep 42, (2023).

92. Beecher, H. K. Experimental Pharmacology and Measurement of the Subjective Response. Science (1979) 116, 157–162 (1952).

93. Roger Chou, M. D., F. S. S. M. D., M. P. H., J. W. M. A., A. Y. A. B. A., R. J. D. Ph., M. P. H. M. A., K. M. M. D., K. D. S. M. D., M. S., Y. Y. M. S., and R. F. Ph. D. Systematic Review on Opioid Treatments for Chronic Pain: Surveillance Report 3. https://www.ncbi.nlm.nih.gov/books/NBK556255/table/ch4.tab1 (2022).

94. Ari, M., Alexander, J. T. & Weyer, G. Prescribing Opioids for Pain. JAMA vol. 329 1789–1790 Preprint at 10.1001/jama.2023.6539 (2023).

95. Volkow, N. & McLellan, T. Opioid Abuse in Chronic Pain — Misconceptions and Mitigation Strategies. N Engl J Med 374, (2016).

96. Volkow, N. D. & Collins, F. S. The Role of Science in Addressing the Opioid Crisis. NEJM 377, 391–394 (2017).

97. Jones, J. et al. Selective Inhibition of Na V 1.8 with VX-548 for Acute Pain. New England Journal of Medicine 389, 393–405 (2023).

98. Yekkirala, A. S., Roberson, D. P., Bean, B. P. & Woolf, C. J. Breaking barriers to novel analgesic drug development. Nature Reviews Drug Discovery vol. 16 545–564 Preprint at 10.1038/nrd.2017.87 (2017).

99. Agarwal, N., Choi, P. A., Shin, S. S., Hansberry, D. R. & Mammis, A. Anterior cingulotomy for intractable pain. Interdiscip Neurosurg 6, 80–83 (2016).

100. Ballantine, H. T., Cassidy, W. L., Flanagan, N. B. & Marino, R. Stereotaxic Anterior Cingulotomy for Neuropsychiatric Illness and Intractable Pain. J Neurosurg 26, 488–495 (1967).

101. R.A. Cohen, R.F. Kaplan, D.J. Moser, M.A. Jenkins, H. W. Impairments of attention after cingulotomy. Neurology 53, (1999).

102. Cohen, R. A. et al. Emotion and personality changes following cingulotomy. Emotion 1, 38–50 (2001).

103. Liberzon, I. et al. mu-Opioid receptors and limbic responses to aversive emotional stimuli. Proc Natl Acad Sci U S A 99, 7084–7089 (2002).

104. Martikainen, I. K. et al. Alterations in Endogenous Opioid Functional Measures in Chronic Back Pain. Journal of Neuroscience 33, 14729– 14737 (2013).

105. Yee, D. M., Crawford, J. L. & Braver, T. S. Dorsal Anterior Cingulate Cortex Encodes the Subjective Motivational Value of Cognitive Task Performance a. bioRxiv 2020.09.20.305482 (2020) doi:10.1101/2020.09.20.305482.

106. Sarafyazd, M. & Jazayeri, M. Hierarchical reasoning by neural circuits in the frontal cortex. Science (1979) 364, (2019).

107. Vázquez, D. et al. Anterior cingulate cortex lesions impair multiple facets of task engagement not mediated by dorsomedial striatum neuron firing. Cerebral Cortex 34, (2024).

108. Becket Ebitz, R. et al. Human dorsal anterior cingulate neurons signal conflict by amplifying task-relevant information. doi:10.1101/2020.03.14.991745.

109. Finkelstein, A., Fontolan, L., Economo, M. N., Li, N. & Romani, S. Attractor dynamics gate cortical information flow during decision-making. bioRxiv (2019) doi:10.1101/2019.12.14.876425.

110. Jones, J. M. et al. A machine-vision approach for automated pain measurement at millisecond timescales. Elife 9, 1–22 (2020).

111. Bohic, M. et al. Mapping the neuroethological signatures of pain, analgesia, and recovery in mice. Neuron 111, 2811-2830.e8 (2023).

112. Zhang, Z. et al. Automated preclinical detection of mechanical pain hypersensitivity and analgesia. Pain 163, 2326–2336 (2022).

113. Wiltschko, A. B. et al. Mapping Sub-Second Structure in Mouse Behavior. Neuron 88, 1121–1135 (2015).

114. Szablowski, J. O., Lee-Gosselin, A., Lue, B., Malounda, D. & Shapiro, M. G. Acoustically targeted chemogenetics for the non-invasive control of neural circuits. Nat Biomed Eng 2, 475–484 (2018).

115. DeNardo, L. A. et al. Temporal evolution of cortical ensembles promoting remote memory retrieval. Nat Neurosci 22, 460–469 (2019).

116. Corder, G. et al. An amygdalar neural ensemble that encodes the unpleasantness of pain. Science (1979) 363, 276–281 (2019).

117. Castro, D. C. et al. An endogenous opioid circuit determines state-dependent reward consumption. Nature 598, 646–651 (2021).

118. Yang, S. H. et al. Neural mechanism of acute stress regulation by trace aminergic signalling in the lateral habenula in male mice. Nat Commun 14, 2435 (2023).

119. Seo, D. et al. A locus coeruleus to dentate gyrus noradrenergic circuit modulates aversive contextual processing. Neuron 109, 2116-2130.e6 (2021).

120. Roth, B. L. Neuron Primer DREADDs for Neuroscientists. doi:10.1016/j.neuron.2016.01.040.

121. Pool, A.-H., Poldsam, H., Chen, S., Thomson, M. & Oka, Y. Recovery of missing single-cell RNA-sequencing data with optimized transcriptomic references. Nat Methods 20, 1506–1515 (2023).

122. Young, M. D. & Behjati, S. SoupX removes ambient RNA contamination from droplet-based single-cell RNA sequencing data. Gigascience 9, (2020).

123. Yao, Z. et al. A taxonomy of transcriptomic cell types across the isocortex and hippocampal formation. Cell 184, 3222-3241.e26 (2021).

124. Cipollari, E. et al. Correlates and Predictors of Cerebrospinal Fluid Cholesterol Efflux Capacity from Neural Cells, a Family of Biomarkers for Cholesterol Epidemiology in Alzheimer’s Disease. Journal of Alzheimer’s Disease 74, 563–578 (2020).

